# Plant-microbe co-evolution: allicin resistance in a *Pseudomonas fluorescens* strain (*Pf*AR-1) isolated from garlic

**DOI:** 10.1101/769265

**Authors:** Jan Borlinghaus, Anthony Bolger, Christina Schier, Alexander Vogel, Martin C. H. Gruhlke, Alan J. Slusarenko

**Affiliations:** RWTH Aachen University

**Keywords:** *Pseudomonas*, antibiotic resistance, horizontal gene transfer, genomic island, ecological niche, microbiota

## Abstract

The antibiotic defense substance allicin (diallylthiosulfinate) is produced by garlic (*Allium sativum* L.) after tissue damage, giving garlic its characteristic odor. Allicin is a redox-toxin that oxidizes thiols in glutathione and cellular proteins. A highly allicin-resistant *Pseudomonas fluorescens* strain (*Pf*AR-1) was isolated from garlic, and genomic clones were shotgun electroporated into an allicin-susceptible *P. syringae* strain (*Ps*4612). Recipients showing allicin-resistance had all inherited a group of genes from one of three similar genomic islands (GI), that had been identified in an *in silico* analysis of the *Pf*AR-1 genome. A core fragment of 8-10 congruent genes with redox-related functions, present in each GI, was shown to confer allicin-specific resistance to *P. syringae*, and even to an unrelated *E. coli* strain. Transposon mutagenesis and overexpression analyses revealed the contribution of individual candidate genes to allicin-resistance. Moreover, *Pf*AR-1 was unusual in having 3 *glutathione reductase* (*glr*) genes, two copies in two of the GIs, but outside of the core group, and one copy in the *Pf*AR-1 genome. Glr activity was approximately 2-fold higher in *Pf*AR-1 than in related susceptible *Pf*0-1, with only a single *glr* gene. Moreover, an *E. coli* Δ*glr* mutant showed increased susceptibility to allicin, which was complemented by *Pf*AR-1 *glr1*. Taken together, our data support a multi-component resistance mechanism against allicin, achieved through horizontal gene transfer during coevolution, and allowing exploitation of the garlic ecological niche. GI regions syntenic with *Pf*AR-1 GIs are present in other plant-associated bacterial species, perhaps suggesting a wider role in adaptation to plants *per se*.

---

Adaptation is the process that tailors organisms to a particular environment and enhances their evolutionary fitness. Plants provide habitats for pathogenic and commensal organisms and generally it is assumed that microorganisms found in association with a given plant host are adapted to that ecological niche as part of the microbiota. Plants produce a vast array of secondary metabolites, many of which are involved in defense against microbes, resulting in a dynamic co-evolutionary arms race in the interaction between plants and their associated microorganisms (Burdon & Thrall, 2009). The organosulfur compounds produced by garlic (*Allium sativum* L.) provide an important example of this scenario. The potent antibacterial activity of garlic was shown in 1944 to be due to mainly to diallylthiosulfinate, which was given the trivial name allicin (Cavallito and Bailey, 1944 a; Cavallito et al., 1944 b). Allicin is formed by the action of alliin-lyase (E.C.4.4.1.4) on alliin (*S*-allyl-L-cysteine sulfoxide) when enzyme and substrate mix after damage to garlic tissues. Allicin is volatile and is responsible for the typical odor of freshly crushed garlic. Alliin-lyase is one of the most prevalent soluble proteins accumulating in garlic bulbs and leaves, indicating a major investment of plant resources into this defense system (Van Damme et al., 1992; Smeets et al., 1997, Borlinghaus et al., 2014). The reaction proceeds rapidly and alliin conversion to allicin is approximately 97 *%* complete after 30 sec. at 23 °C (Lawson & Hughes, 1992). The evolutionary investment of garlic in this mechanism is further emphasized by the fact that a single clove of approximately 10 g fresh weight can liberate up to 5 mg of allicin (Lawson et al., 1991; Block, 2010).

Allicin is an electrophilic oxidant that oxidizes thiols, or more precisely the thiolate ion, in a modified thiol-disulfide exchange reaction (Scheme 1), producing *S*-allylmercapto disulfides (Rabinkov et al., 2000; Müller et al., 2016). Cellular targets are accessible cysteines in proteins, and the cellular redox buffer glutathione (GSH). In this way, allicin can inhibit essential enzymes (Wills, 1956) and shift cell redox balance (Gruhlke et al., 2010). Allicin causes oxidative stress and was shown to directly activate the Yap1 transcription factor which coordinates the oxidative stress response in yeast (Gruhlke et al., 2017). Inded, allicin has been described as a ‘redox toxin’ (Gruhlke et al., 2010).

**Scheme 1.**
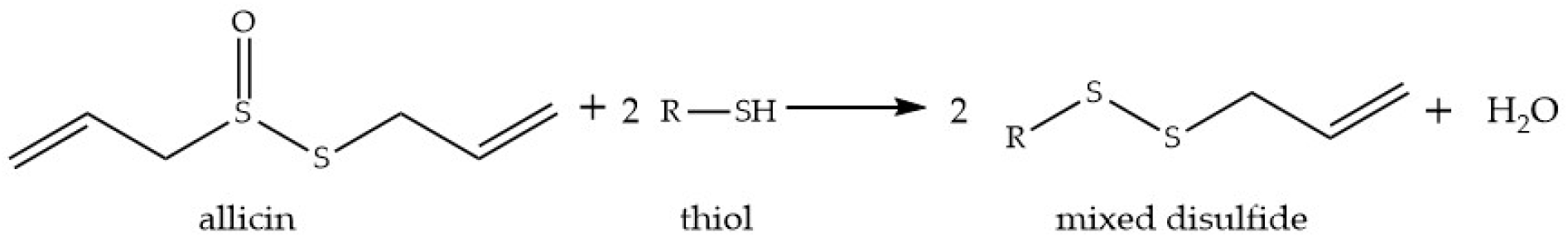
Reaction of allicin with a cellular thiol to produce an S-allylmercapto mixed disulfide.

Thus, allicin has multiple sites and mechanisms of action in cells and is a concentration-dependent biocide active against bacteria, fungi oomycetes and mammalian cells (Borlinghaus et al., 2014). Because of allicin’s thiol reactivity, the cellular redox buffer glutathione (GSH) plays a central role in the resistance of cells at sublethal allicin concentrations (Gruhlke et al., 2010, 2017; Leontiev et al., 2018).

Because of its broad range of cellular targets, it is not easy for an organism to mutate simply to be more resistant to allicin. However, sensitivity to allicin does vary between different bacterial species and isolates (Reiter et al., 2017) and, as described here, we isolated a highly allicin-resistant *Pseudomonas fluorescens* from a clove of garlic. How resistance against allicin might have evolved in *Pf*AR-1 is an intriguing question. One possibility is horizontal gene transfer (HGT), i.e. the sharing of genetic material between organisms that are not in a parent– offspring relationship (Soucy et al. 2015). Genomic islands arising by HGT generally show a different average GC content and codon usage to the rest of the genome. HGT is a widely recognized mechanism for adaptation in bacteria and microbial antibiotic resistance and pathogenicity are often associated with HGT (Maclean & San Milan 2019). Especially large, chromosomally-integrated regions obtained by HGT, that are referred to as genomic islands (GIs), are known to expand the ecological niches of their host bacteria for complex and competitive environments (Soucy et al. 2015).

In the work reported here, we isolated a highly allicin-resistant bacterium from its ecological niche on garlic, an environment hostile to non-adapted microorganisms, and we used a shotgun genomic cloning strategy to functionally identify genes conferring allicin resistance. The annotated gene functions of resistance-conferring genes throws light on the complex molecular mechanisms of resistance of *P. fluorescens* strain *Pf*AR-1 to the defense substance allicin, which has multiple targets and modes of action within the cell. This functional approach was complemented by whole-genome sequencing which revealed unique genomic features in comparison to other Pseudomonads. Both approaches independently identified the same sets of genes, validating the strategy. The complex genomic structures associated with conferring allicin resistance, arose via horizontal transfer and duplication, revealing the evolutionary investment associated with *Pf*AR-1 being able to exploit garlic as an environmental niche.

## Material & Methods

Additional information about bacterial strains, plasmids, primers, chemical allicin synthesis, and allicin analysis are given in the supplementary material (SM).

### *Pf*AR-1 genomic library construction

Genomic DNA of *Pf*AR-1 was extracted and partially digested with Sau3AI to obtain approx. 10 kbp fragments which were subcloned in BamHI digested pRU1097 (SM7, SM8).

### Transposon mutagenesis

Genomic clone 1 plasmid was transposon mutagenized in *Ps*4612 using the transposon IS-Ω-km/hah on plasmid pSCR001. Plasmid DNA was transferred from *Ps*4612 to *E. coli* MegaX to seperate chromosomal and plasmid Tn integrations as described in SM9.

### Overexpression of putative allicin resistance genes

Genes were amplified from genomic clone 1 plasmid DNA with overhangs for NotI/XbaI subcloning in pJABO (SM10).

### Complementation of *E. coli* Δglr

*Pf*AR-1 *glr1* was cloned in vector pJABO5 via homologous recombination in yeast (SM11).

### Inhibition zone assays

Bacteria were freshly grown from an optical density at 600 nm (OD_600_) of 0.05 to OD_600_ = 0.2-0.3. 300 μl liquid culture were seeded in 20 ml 50 °C warm agar medium in round Petri dishes (Ø = 9 cm). Holes (Ø = 0.6 cm) were punched out to apply the test solution. Plates were then incubated over night.

To investigate the effects of different oxidants, bacterial cultures were freshly inoculated from over-night cultures and grown from an OD_600_ of 0.05 to an OD_600_ of 1.0 to 2.0. OD_600_ of all strains was adjusted to an OD_600_ of 1.0 and 125 μl were applied on top of 20 ml solid medium in round Petri dishes (Ø = 9 cm). Bacteria were spread with glass beads (Ø = 3 mm) by soft shaking to avoid heat stress compared to seeded agar. Holes (Ø = 0.6 cm) were punched out to apply the test solution.

### Streak tests

A bacterial colony was harvested from agar plates in liquid medium, mixed, and streaked on 20 ml prepared solid media with holes in the center (Ø = 0.6 cm) to apply test solutions.

### Drop tests

Liquid bacterial cultures grew over night and were subsequently diluted ten-fold from OD_600_ = 1.0 to OD_600_ = 10^−5^. 5 μl of each bacterial dilution was dropped on solid media (LB) containing different amounts of allicin. Plates were incubated at 37 °C over night.

### Protein extraction and glutathione reductase activity assay

Pseudomonads were grown over night in liquid M9JB medium (SM2) to decrease slime production. Crude bacterial cell lysate was prepared from bacteria by vortexing with glass beads. Glutathione reductase activity assay was performed as described in SM 12.

### Genome sequencing of *Pf*AR-1

*Pf*AR-1 was grown in KB medium at 200 rpm and 28 °C over night. For DNA extraction, 15 ml of overnight culture was washed three times in 1×TE with 50 mM EDTA by repeated pelleting at 5000 xg and resuspension by vortexing. The subsequent cell lysis was performed as described in Sambrook & Russel (2001) for gram-negative bacteria. From this material, three Illumina paired-end libraries were created, and run multiplexed in conjunction with other samples, twice as 2×100 paired-end runs on a HiSeq 2000, and once as a 2×311bp paired-end run on a MiSeq. The resulting data was filtered by Trimmomatic V0.32 (Bolger et al., 2014) and assembled using SPAdes V3.5.0 (Bankevich et al., 2012). The resulting assembly was largely complete, with a total size of 6.3 Mbp, but it was still relatively fragmented with 40 scaffolds of 1 kbp or larger, and an N50 of 370 kbp.

In order to fully resolve the genome into one contig, two additional long read datasets were generated on the Pacific Biosciences RS-II platform. For DNA extraction, 15 ml of overnight culture were washed three times in 1×TE with 50 mM EDTA by repeated pelleting at 5000 xg and resuspension by vortexing. The subsequent cell lysis was performed as described in Sambrook and Russel (2001) for Gram-negative bacteria. Further depletion of contaminating polysaccharides was achieved by application of the Pacific Biosciences protocol (Pacific Biosciences, 2019) for gDNA cleanup. The final DNA was eluted in RNAse free water and quality was determined using Nanodrop for purity and Qubit for quantification. Sequencing was performed by GATC. The resulting two datasets, combined with the Illumina datasets described above, were then assembled, using SPAdes 3.5.0, yielding a single contig sequence of ~6.26Mbp.

Self-alignment of this contig revealed that 9,642 bp sequence was duplicated on each end which was then removed from one end. In order to simplify cross-genome comparisons, this sequence was aligned against the *Pf*0-1 reference sequence, and oriented to match, resulting in the 6,251,798 bp *Pf*AR-1 assembly. The completed genome was then submitted to the RAST webserver (Aziz et al., 2008; Overbeek et al., 2014; Brettin et al., 2015) for automatic structural and functional annotation.

### *In-silico* analysis of the *Pf*AR-1 Genome

The low-GC regions identified in the *Pf*AR-1 genome were initially compared manually by cross-referencing the functional annotation of genes. This revealed a list of genes from each region which have a potentially common origin. After removing low confidence protein annotations, which were both unique to a single region and lacking a definitive functional annotation (*Pf*AR1.peg.1058 and *Pf*AR1.peg.1070), the remaining genes were manually reconciled into a putative ancestral arrangement of 26 genes.

### Comparison of Putative HGT Regions across the *Pseudomonas* Genus

A set of bait genes was created based on the putative 26-gene ancestral arrangement described above. Since these 26 groups were generally represented in more than one region, the set comprised 57 sequences in total. All available *Pseudomonas* sequences, comprising 215 complete genomes and 3,132 draft genomes, were downloaded from the Pseudomonas Genome Database website (https://www.pseudomonas.com/), and queried for the bait sequences using BLAST. Similarity was calculated using a sliding window of 40 genes, and regions which exceeded a normalized bit-score total of 5 were selected.

### Inter-Species Codon Analysis

Synonymous codon usage statistics were calculated for the full *Pf*AR-1 genome, the 3 putative HGT regions, the 3347 other available *Pseudomonas* genomes, and 8 representative *non-Pseudomonas* Gammaproteobacteria (*Acinetobacter baumannii* AC29, *Alkanindiges illinoisensis, Azotobacter vinelandii* DJ, *Escherichia coli* K12 MG1655, *Moraxella catarrhalis, Perlucidibaca piscinae, Rugamonas rubra, Ventosimonas gracilis*). After removing methonine and tryptophan, which have only one codon, the remaining codons were analysed using Principle Component Analysis (PCA).

### Phylogenetic Comparson of Whole Genome vs RE-like sequences

Whole genome phylogenetic analysis was performed using OrthoFinder (Emms and Kelly, 2015; version 1.1.8, https://github.com/davidemms/OrthoFinder/releases/tag/1.1.8) to place the newly sequenced *Pf*AR-1 genome in its phylogenetic context, using a subset of 280 *Pseudomonas* genomes supplemented by 4 more distant genomes downloaded from NCBI GenBank, namely *Azotobacter vinelandii* DJ, *Acinetobacter baumannii* AC29, *Escherichia coli* K12 MG1655 and *Burkholderia cenocepacia* J2315 which served as an outgroup. The 280 *Pseudomonas* genome subset consisted of a) all 215 complete genomes, b) the draft genomes showing a substantial hit against the putative-HGT gene set, as described above, and c) 9 *Pseudomonas* genomes with unusual codon usage (*P. lutea, P. luteola*, P. sp HPB0071, *P*. sp FeS53a, *P. zeshuii, P. hussainii* JCM, *P. hussainii* MB3, *P. caeni* and *P. endophytica*).

In a second analysis, the 3 putative-HGT from *Pf*AR-1 were compared against the corresponding regions from other *Pseudomonas* genomes, identified as described above. For this analysis, the sequences from each genomic island region was re-ordered according to the best match against the 26 bait group sequences, concatenated to form a single pseudo-sequence and aligned using MAFFT (version 7, Katoh and Standley 2013). The resulting multiple alignment was accessed using “fitch” from Phylip (version 3.69) and the resulting trees were visualized using FigTree (version 1.4.3, https://github.com/rambaut/figtree/releases/tag/v1.4.3).

### IslandViewer Analysis

For independent confirmation of the HGT analysis, the *Pf*AR-1 genome was submitted to the IslandViewer 4 (Bertelli et al., 2017) website, for assessment regarding horizontal gene transfer events.

### Additional annotation of genomic repeat regions

Gaps in the annotation of genomic repeats with putative horizontal origin indicated incomplete annotation, also implicated by a low gene density (1 gene per 1.3 - 1.6 kbp), which is expected to be one gene per 1 kbp in bacterial genomes (Koonin and Wolf, 2008). Regions were submitted individually without the remaining genome sequence to the RAST webserver, thereby closing annotational gaps (1 gene per 0.90 kbp in average). Remaining DNA regions without annotation were manually curated using NCBI open reading frame finder and BLASTp.

### Dot plot and %GC content analysis

For dot plot analysis and %GC content analysis and comparison, Genome Pair Rapid Dotter (GEPARD, Krumsiek et al., 2007), Artemis Comparison Tool (ACT, Carver et al. 2005), and UGENE (Okonechnikov et al., 2012) were used, respectively.

### Congruent set of genes and copy number analysis

Analysis was performed by batch translation of the coding sequences of the *Pf*AR-1 genomic repeats into peptide sequences using coderet from the emboss suite (Rice et al. 2000), and compared these against all other peptide sequences from the genomic repeats and the remaining genome, respectively. Peptides with a minimal peptide length of ≥ 100 amino acids were compared using BLASTp (Camacho et al. 2009) combined with the graphical user interface visual blast (Melee 2016). Significantly similar sequences were defined by a minimal sequence similarity of ≥ 25 % and with an E-value ≤ 0.0001.

## Results

### An allicin-resistant *Pseudomonas fluorescens* isolated from garlic

We reasoned that if allicin-resistant bacteria were to be found in nature, then it would likely be in association with garlic cloves. Therefore, bacteria isolated from garlic bulbs were tested for allicin resistance in a Petri plate agar diffusion test with bacteria-seeded agar. An extremely resistant isolate was detected that was able to grow right up to the allicin solution in the agar, whereas allicin-sensitive *E coli* DH5α and *Pseudomonas syringae* pv. *phaseolicola Ps*4612 isolates showed large inhibition zones (Fig. 1). The allicin-resistant isolate was putatively identified by Sanger sequencing of the ribosomal ITS (internal transcribed spacer) as a *Pseudomonas fluorescens*, and we therefore named it *Pf*AR-1 (*Pseudomonas fluorescens* Allicin Resistant-1).

**Fig. 1.**
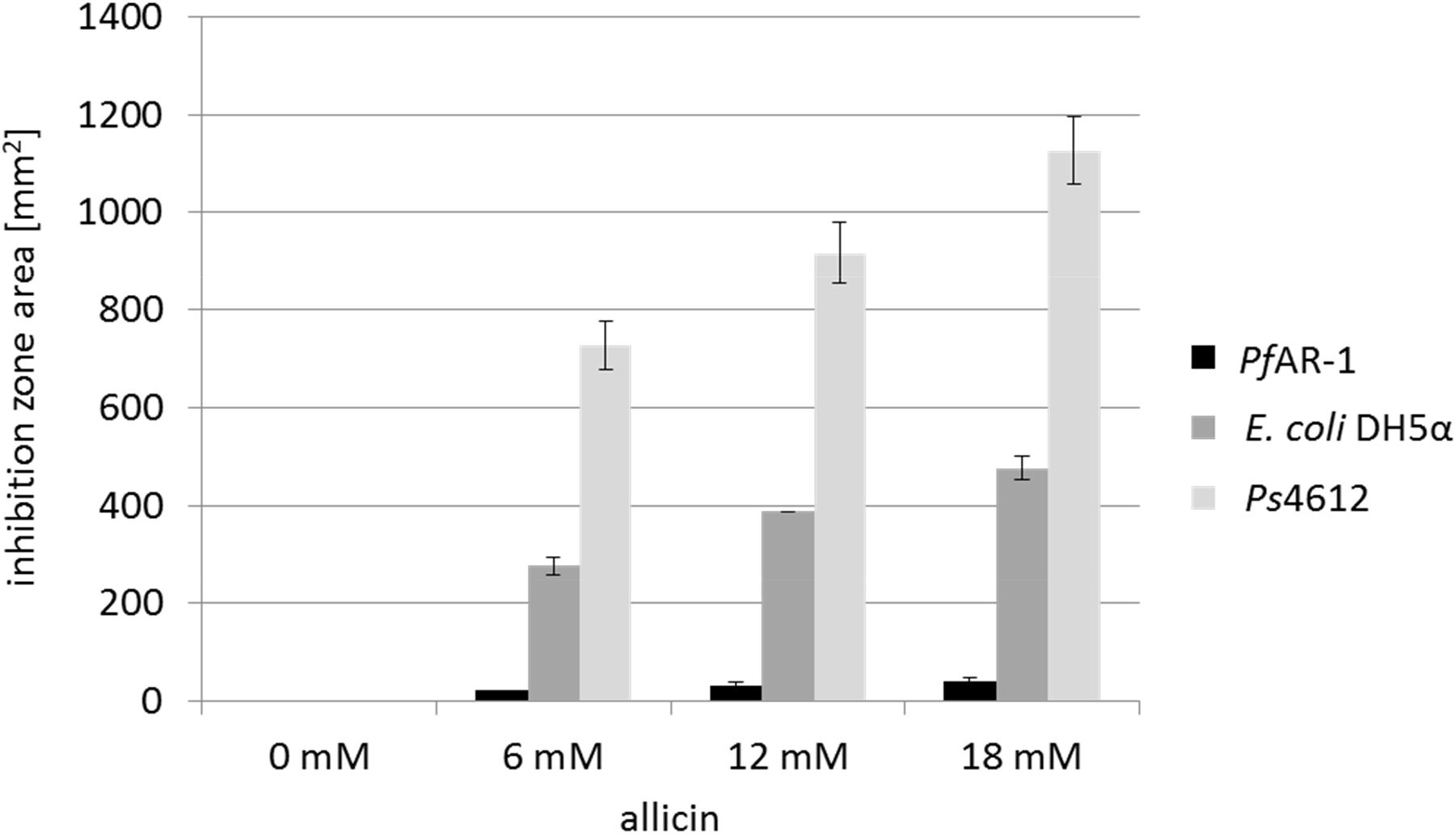
Comparison of the sensitivity of *Pf*AR-1, *E. coli* DH5α and *P. syringae Ps*4612 against allicin. The area of the inhibition zones in an agar diffusion test is shown for 40 μl of a range of allicin concentrations applied centrally to a well in the agar medium. n = 3

### Identification *Pf*AR-1 genomic clones conferring allicin-resistance

*Pf*AR-1 genomic clones were shotgun electroporated into cells of highly-allicin-susceptible *Ps*4612. In all, 1.92 x 10^8^ clones were screened giving approximately 33 x library coverage. Resistant recipients were selected on medium containing 75 μM allicin. No allicin-resistant colonies grew on the control plate of *Ps*4612 cells that had been electroporated with the empty vector. Eight allicin-resistant transformants were confirmed in streak tests (Fig. 2). The degree of allicin resistance conferred by genomic clone 8 was slightly less than that conferred by the clones 1-7, as evidenced by the slightly greater clear zone where growth was inhibited (Fig. 2). Restriction analysis of the genomic inserts revealed that the eight resistance-conferring clones were all approximately 10 kb in size.

**Fig. 2.**
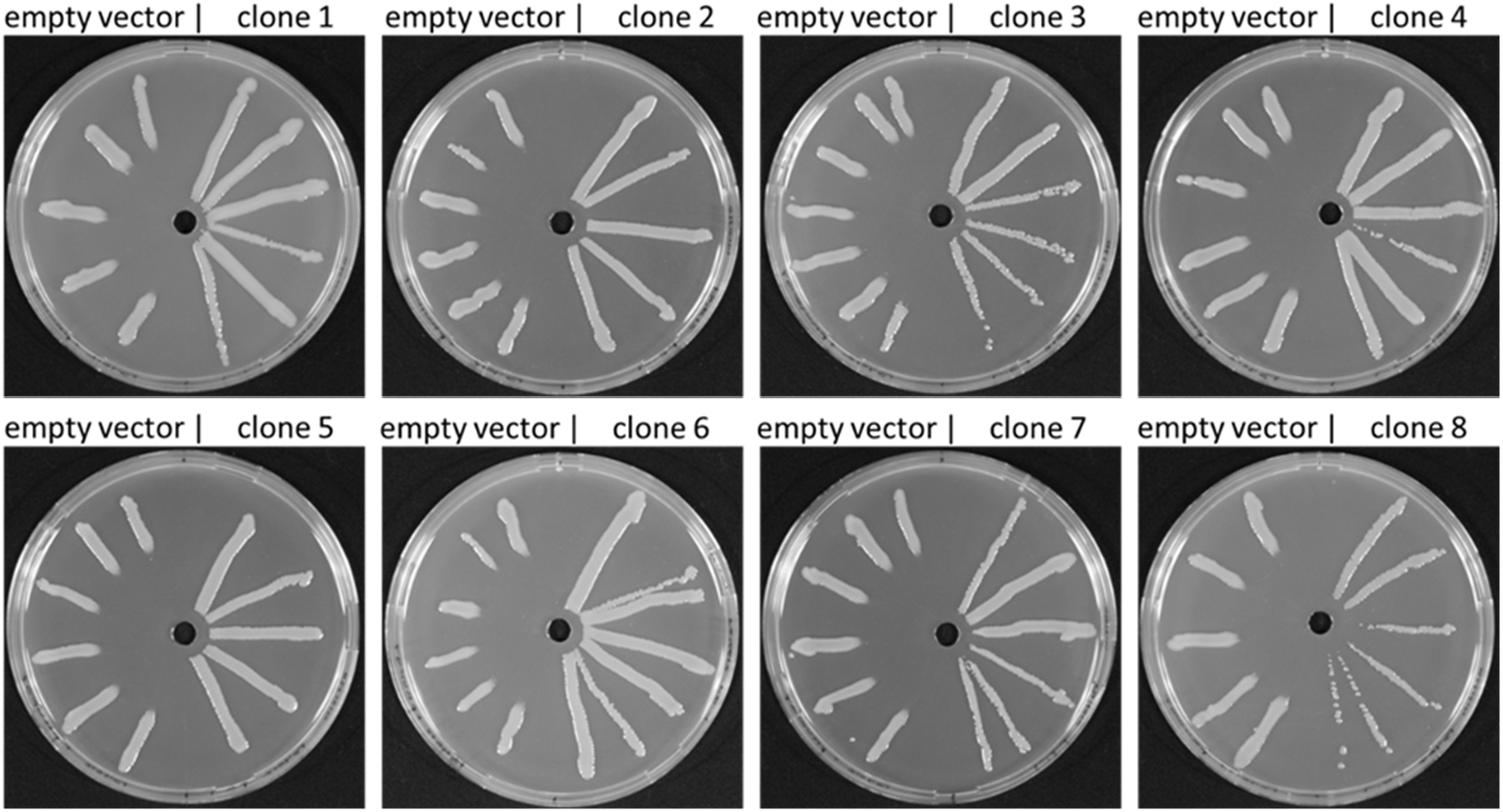
Allicin resistance conferred by genomic clones from *Pf*AR-1 electroporated into *Ps*4612. On the left half of each Petri plate are *Ps*4612 cells transformed with empty vector and on the right half *Ps*4612 cells transformed with vector containing a genomic insert. The central wells contained 30 μl of 32 mM allicin solution.

### Genomic sequencing of *Pf*AR-1

The *Pf*AR-1 genome was sequenced using a combined Illumina and Pacific Biosciences dataset, and assembled into a single chromosome, as described in the Materials and Methods. The resulting genome was 6,251,798 bp, and had an overall GC content of 59.7%. A total of 5,406 putative protein-coding sequences were detected, in addition to 73 tRNAs and 6 rRNA clusters. With an average nucleotide identity (ANI) of 85.94% (SM18), the closest relative to *Pf*AR-1 in the data bases was *Pseudomonas fluorescens* reference strain *Pf*0-1, confirming the prior ITS-based assessment.

### Characterization of resistance-conferring clones

Sanger sequencing of the clone ends was used to identify the origin of the clones within the sequenced *Pf*AR-1 genome. This revealed that clones 1 and 8 had unique origins, whereas clones 2-7 were identical to each other. Thus, 3 relatively compact resistance-conferring genomic regions had been identified. Genes carried on the clones had preponderantly redox-related functions (Fig. 3 *A*, Table 1). The overall arrangement of the genes was highly conserved among the clones. Clones 1-7 contained two sets of genes conserved in the direction of transcription: *osmC, sdr, tetR, dsbA*, and *trx* and the second set being *ahpD, oye, 4-ot, kefF*, and *kefC* (Fig. 3 *C*). Clone 8, which conferred slightly less allicin resistance than the other clones (Fig. 2), lacked the *ahpD* and *oye* genes. The *kefF* and *kefC* genes are part of a glutathione-regulated K+ efflux/H+ influx system and are classified as transporters, although they too are regulated by cellular glutathione, and thus are also redox-dependent.

**Fig. 3.**
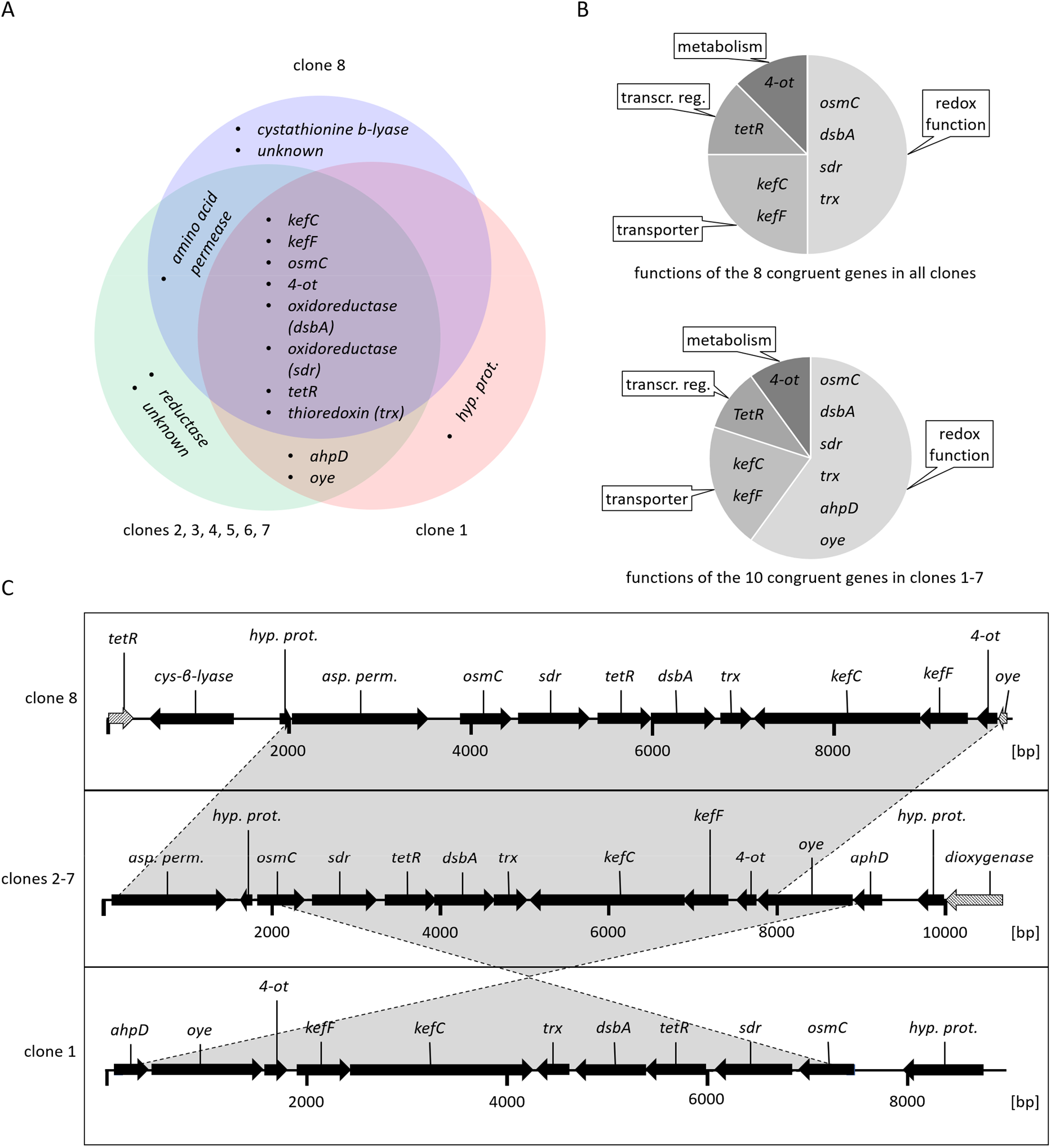
Characteristics of the allicin-resistance-conferring *Pf*AR1 genomic clones. (*A*) Venn diagram showing congruent genes. (*B*): Congruent genes grouped by function. (*C*) Arrows show the direction of transcription. Grey shaded arrows in clones 8 and 2-7 represent truncated genes. (unkn.: unknown i.e. no similar protein or protein domains in databases; transcr. reg: transcriptional regulator; hyp. prot.: hypothetical protein).

**Table 1:** Congruent set of genes identified in the genomic clones 1-7 that conferred allicin resistance to *E. coli* K12 DH10B and *P. syringae* pv. *phaseolicola* 4612.

| **PfAR-1 genes<sup>1</sup>** | **Reported function in other bacteria** | **References** |
| --- | --- | --- |
| *alkylhydroperoxidase (ahpD)* | Part of the carboxymuconolate decarboxylase (CMD) family. NADH-dependent AhpD/AhpC system confers oxidative stress resistance in *Mycobacterium tuberculosis*. | Bryk et al., 2002; Koshkin et al., 2003 |
| *old yellow enzyme (oye)* | OYE protein family contains a diverse set of NADPH-dependent dehydrogenases that reduce $\alpha,\beta$ unsaturated aldehydes and ketones. OYE was reported to be part of the oxidative stress response in yeast and in *Bacillus*. | Stott et al., 1993; Vaz et al., 1995; Trotter et al., 2006; Fitzpatrick et al., 2003 |
| *4-oxalocrotonate tautomerase (4-ot)* | 4-OT converts 2-hydroxymuconate to the $\alpha,\beta$-unsaturated ketone 2-oxo-3-hexendioate. | Whitman et al., 1991; Whitman 2002 |
| *glutathione-regulated potassium-efflux system protein F (kefF)* | KefF is a cytoplasmic regulator of KefC. KefF is activated by glutathione-adducts and subsequently activates KefC. | Munro et al., 1991; Miller et al., 2000; Meury and Kepes, 1982; Elmore et al., 1990; Ferguson et al., 1997; Lyngberg et al., 2011 |
| *glutathione-regulated potassium-efflux system protein C (kefC)* | KefC is a proton import/potassium export antiporter. KefC activity is tightly regulated by glutathione and KefF. Active KefC confers resistance against electrophiles like *N*-ethylmaleimide in *E. coli*. |  |
| *Thioredoxin (trx)* | Trx are dithiol-disulfide oxidoreductases that help maintain the thiol groups of proteins in a reduced state | Holmgren, 2000 |
| *disulfide bond protein A (dsbA) / frnE-like* | DsbA in *E. coli* is responsible for introduction of disulfide bonds in nascent polypeptide chains in the periplasmic space. Other Dsb members show chaperone-like functions. FrnE is a member of the DsbA family and was reported to confer oxidative stress resistance in *Deinococcus radiodurans*. | Bardwell et al., 1991; Kamitani et al., 1992; Khairnar et al., 2013 |
| *transcriptional regulator tetR* | Transcriptional repressors widely distributed among different bacteria. | Ramos et al., 2005 |
| *short chain dehydrogenase (sdr)* | The family contains dehydratases, decarboxylases or simple oxidoreductases. | Kavanagh et al., 2008 |
| *osmotically inducible protein C (osmC)* | The family contains peroxiredoxins which play a role in oxidative stress defense. OsmC confers resistance to organic hydroperoxides such as cumene hydroperoxide in *E. coli*. | Lesniak et al., 2003 |

<sup>1</sup>annotation is based on protein domains and corresponding protein families

### *Pf*AR-1 genomic clones conferred allicin-specific-resistance to *E. coli* and *P. savastanoi* 1448

Clones 1, 5 and 8 were chosen so that each group was represented for the initial functional tests. The genomic clones were electroporated into the allicin-susceptible *E. coli* K12 DH10B and *Pseudomonas savastanoi* (formerly *P. syringae* pv. *phaseolicola*) 1448A strains. *Ps*1448A was chosen for further experiments because, in contrast to *Ps*4612, its genome has been sequenced. Various oxidants were tested (Fig. 4 *A*), and it was found that the genomic clones conferred allicin-specific resistance in both *E. coli* and *Ps*1448A, as evidenced by a reduction in inhibition zone area against allicin, but not the other oxidants tested (Fig. 4 *B*). The degree of allicin-resistance conferred by genomic clones 1 and 5 was similar, but as previously observed in the streak test with *Ps*4612 (Fig. 2), clone 8 was less effective than the other clones (Fig. 4 *B*).

**Fig. 4.**
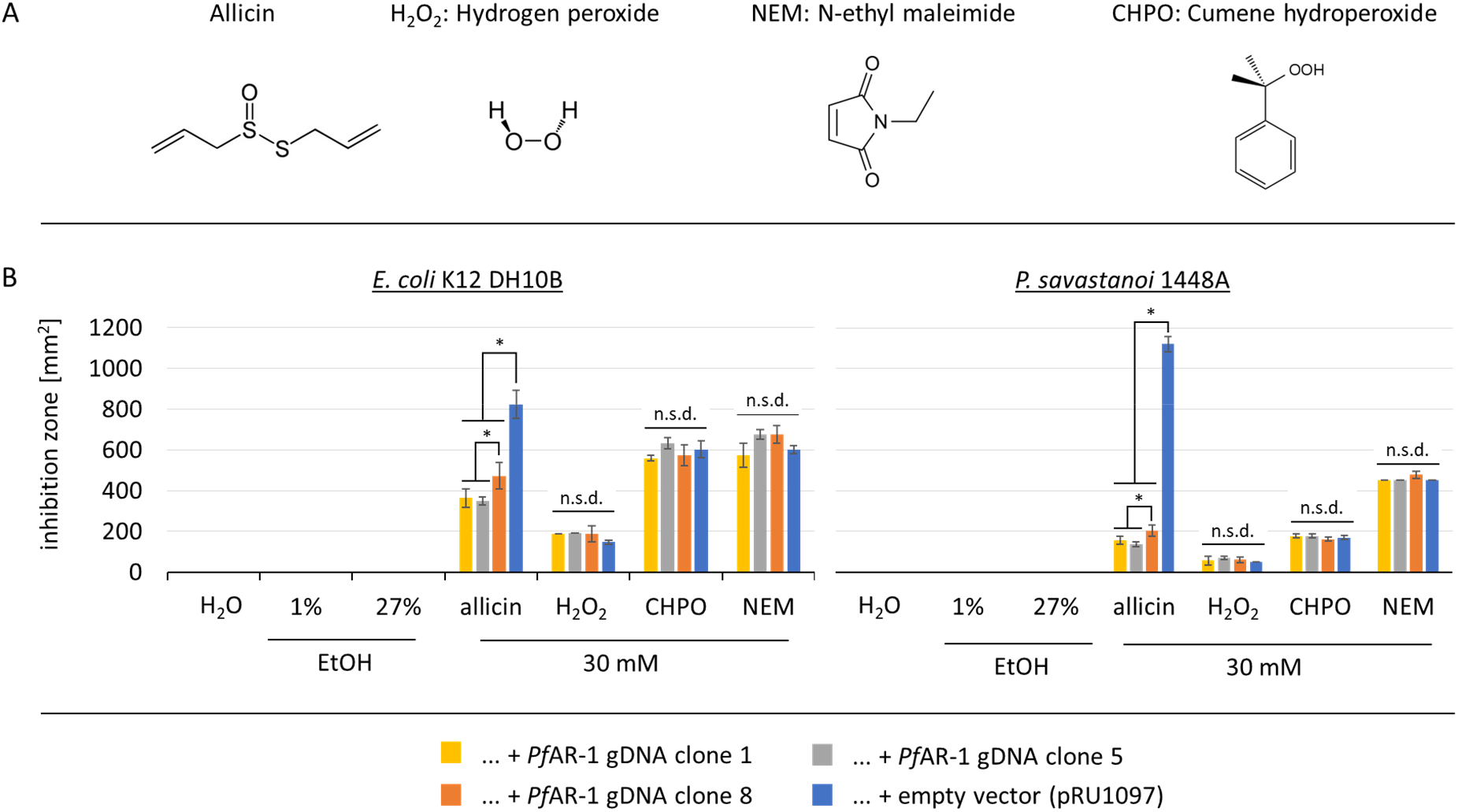
*Pf*AR-1 genomic clones conferred resistance against allicin, but not other oxidants. 40 μl of 30 mM allicin, H_2_O_2_, *N*-ethylmaleimide (NEM) or cumene hydroperoxide (CHPO), were applied to wells cut in bacteria-seeded agar. Ethanol was used as a solvent for NEM and CHPO and 1% and 27% ethanol, respectively, were included as controls. (*A*) Chemical formulas of the oxidants tested. (*B*) Areas of the inhibition zones (mm^2^) are shown for the recipients containing genomic clones 1, 5, 8 or the empty vector. (n = 3 or more, * = p <0.05, Holm-Sidak method for all pairwise comparison), n.s.d. = no significant difference.

### Contribution of individual genes to allicin resistance

The contribution of individual genes to allicin resistance was investigated by transposon mutagenesis of clone 1 in *E. coli* and screening Tn-mutants for loss-of-function. In addition, subcloning and over-expression of individual genes in *Ps*4612 was undertaken, with screening to assess for gain-of-function.

In total, 86 out of the 132 Tn-mutants investigated showed a decrease in allicin resistance compared to non-mutagenized genomic clone 1 in a streak assay. Tn-mutants were examined by sequencing and the positions of Tn insertions are shown in Fig. 5 *A*. No Tn-insertions were found in the *osmC, sdr* or *tetR* genes, but for the majority of the remaining genes, several independent Tn insertion sites were found and these showed a tendency towards decrease in resistance SM13). Tn-mutants for each gene were selected for testing in a more sensitive drop test (Fig. 5 *B*). In the absence of allicin stress, all Tn-mutants grew less well than controls (wt clone 1 and empty vector), as evidenced by the lower colony density visible at the 10^−4^ and 10^−5^ dilutions, respectively. Tn insertions in the vector backbone, or the genes encoding the hypothetical protein *kefF*, or the upstream region of *4-ot*, all had no visible effect on the allicin-resistance phenotype compared to clone 1, at either 150 or 200 μM allicin. In contrast, Tn insertions in either *dsbA, trx, kefC, oye*, or *ahpD*, led to a clear loss of allicin resistance at both allicin concentrations. *ahpD::Tn* showed by far the highest loss of allicin resistance, and resembled the empty vector control. *ahpD* potentially codes for an alkylhydroperoxidase, and the data suggest that this protein plays a major role in being able to confer allicin resistance to *Pf*AR-1. The contributions of the *dsbA* and *trx* genes to allicin resistance were more than those of the *kefC* and *oye* genes, but all of these Tn-mutants showed a clear allicin phenotype, especially at the 200 μM allicin level (Fig. 5 *B*).

**Fig. 5.**
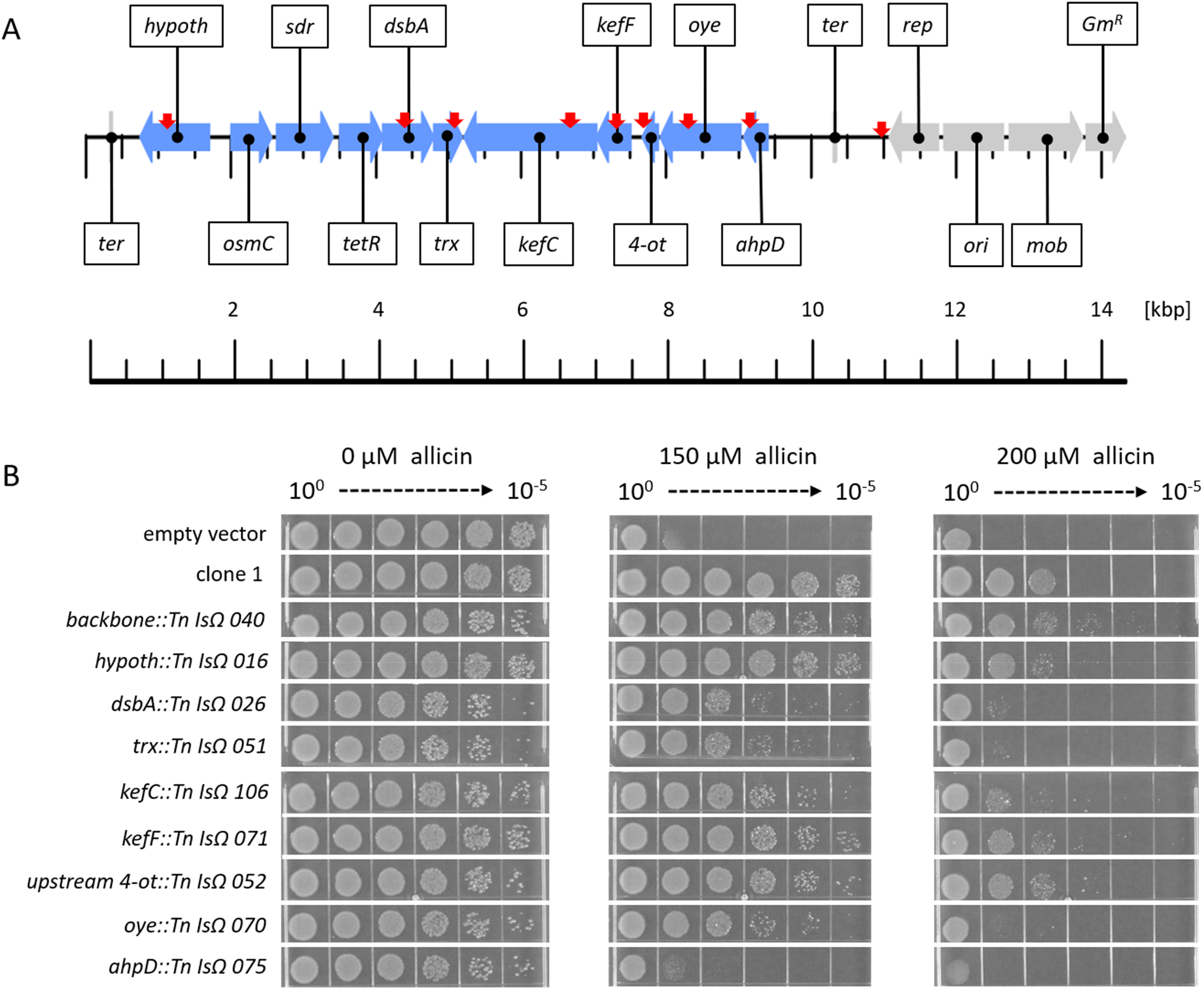
Transposon mutagenesis of genes on clone 1. (*A*) Linear genetic map of *Pf*AR-1 genomic clone 1. *Pf*AR-1 genes are shown in blue, whereas genes on the vector backbone are shown in grey. The position of transposon insertions is indicated by red arrows. (*B*) *E. coli* MegaX DH10B transformed with clone 1, or empty vector, was compared with transposon-insertion mutants with transposon-insertion mutants in drop tests. All cultures were diluted to OD_600_ = 1 (= 10^0^) and 5 μL of a 10^n^ dilution series down to 10^−5^ was dropped onto LB medium supplemented with different allicin concentrations.

### Overexpression of *ahpD* and *dsbA* conferred high allicin-resistance to *Ps*4612

The set of congruent genes on clone 1 were cloned individually in an expression vector to investigate the contribution of each gene to allicin resistance. *Ps*4612 was used for these experiments, because we reasoned that even a small gain in resistance should easily be visible in this highly susceptile allicin isolate. Of the congruent genes from clone 1, only *ahpD* and *dsbA* showed a gain of resistance when overexpressed individually. The resistance conferred by *ahpD* was almost as high as by the intact clone 1. Overexpression of *dsbA* in *Ps*4612 also caused a clear gain of resistance (Fig. 6).

**Fig. 6.**
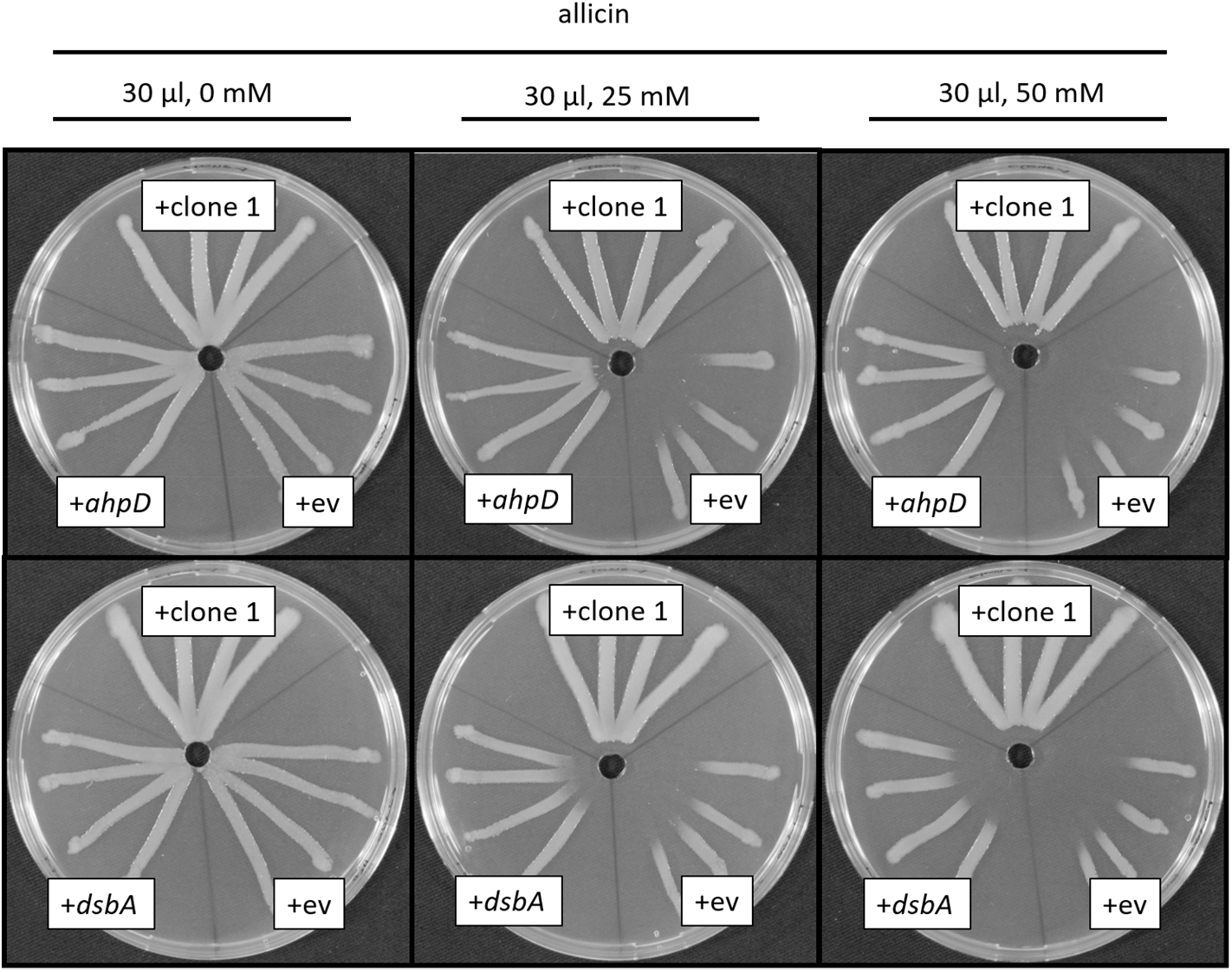
Overexpression of *AhpD* or *DsbA* conferred allicin resistance to *Ps*4612. *ahpD* and *dsbA* were transformed in *Ps*4612. Transformants were streaked from the center to the edges of the plate. Test solutions (30 μl, water, 25 or 50 mM allicin) were applied to each well and the plates were incubated for 48 h at 28 °C.

### *In-silico* analysis of the *Pf*AR-1 *Genome*

As stated above, *Pseudomonas flourescens Pf*0-1 is *Pf*AR-1’s closest previously sequenced relative. Nonetheless, dot matrix alignment of the *Pf*0-1 and *Pf*AR-1 genomes revealed substantial differences. The *Pf*AR-1 chromosome had a central inverted region with respect to *Pf*0-1, and 3 large genomic islands (GI) with lower GC content (<55%), that were absent in *Pf*0-1 (Fig. 7 *A*, *B*). The combination of low GC content and absence from a near-relative genome suggests that these regions might have arisen by horizontal gene transfer. Further analysis revealed that each of the 3 genomic islands (GI1, GI2, and GI3) contained a highly similar region, which we labeled RE1, RE2 and RE3, respectively. These genes within these regions had many annotations in common and a syntenic organization (SM14), suggesting a shared origin.

**Fig. 7.**
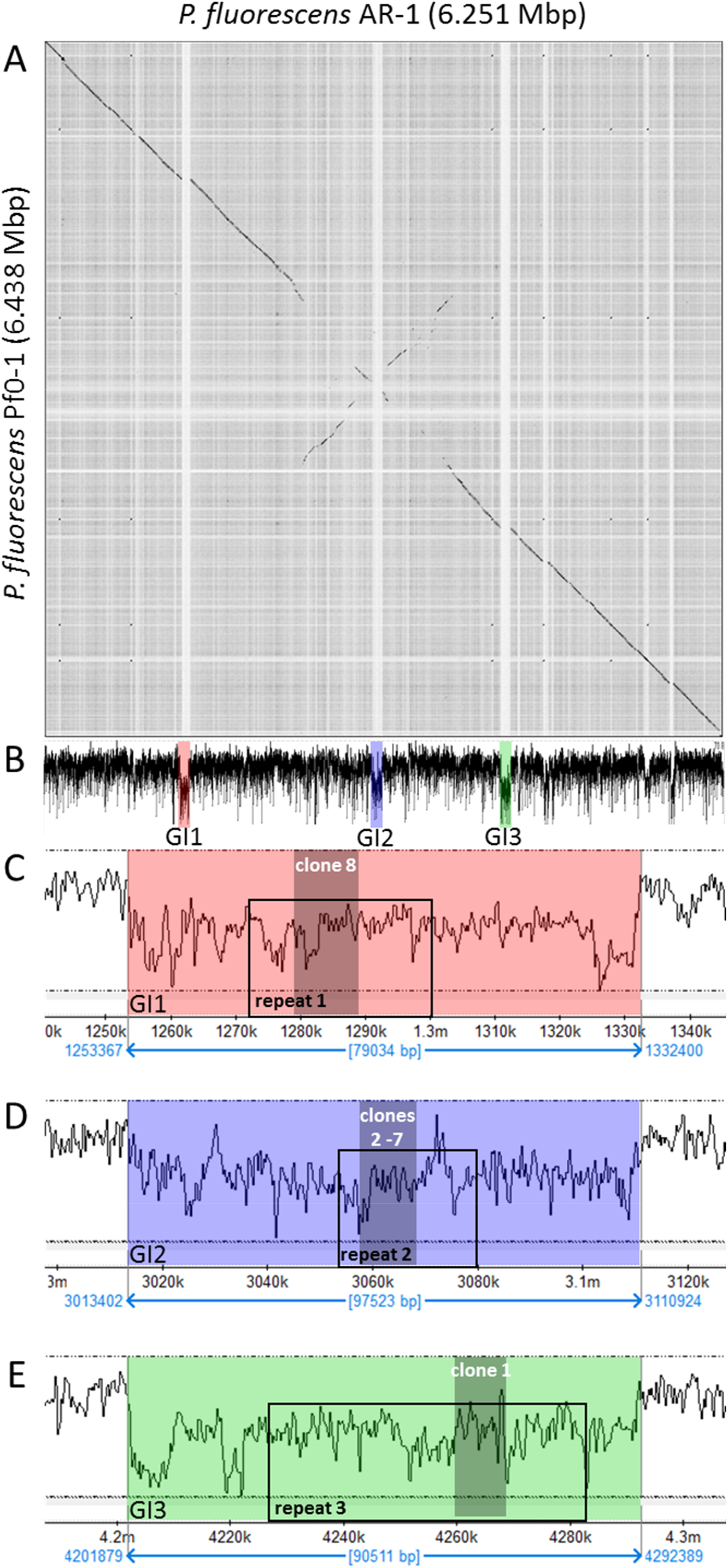
Genomic characteristics of *Pf*AR-1. (*A*) Dot plot alignment of the *Pf*AR-1 and *Pf*0-1 genomes. Numbering is from the putative origin of replication (*oriC*) loci. The disjunctions arising because of inserts in *Pf*AR-1 not present in *Pf*0-1 are clearly visible. (*B*) The GC-content pf the *Pf*AR-1 chromosome with. GI1, GI2 and GI3 marked in red, blue and green, respectively. (*C*) The low GC content region GI1 enlarged to show the position of repeat 1 (RE1) and the location of allicin-resistance-conferring genomic clone 8. (*D*) The low GC content region GI2 enlarged to show the position of RE2 and the location of allicin-resistance-conferring genomic clones 2 −7. (*E*) The low GC content region GI3 enlarged to show the position of RE3 and the location of allicin-resistance-conferring genomic clone 1.

The maintenance of high-similarity regions is generally rare in prokaryotes, and typically requires that such regions offer a substantial evolutionary benefit. Intriguingly, the allicin-resistance-conferring clones found in the functional analysis originated within these 3 regions (Fig. 7 *B, C, D* and *E*), suggesting that the evolutionary benefit may be, in fact, increased allicin resistance. Interestingly, each genomic clone covered almost the complete corresponding repeat region, and thus the genes shared between the RE regions (Fig. 8) matches closely with those shared between the clones (Fig. 3 *B*). Given this strong evidence of importance, possible origins for the putative horizontal gene transfer (HGT) regions into the *Pf*AR-1 genome were investigated more closely.

**Fig. 8:**
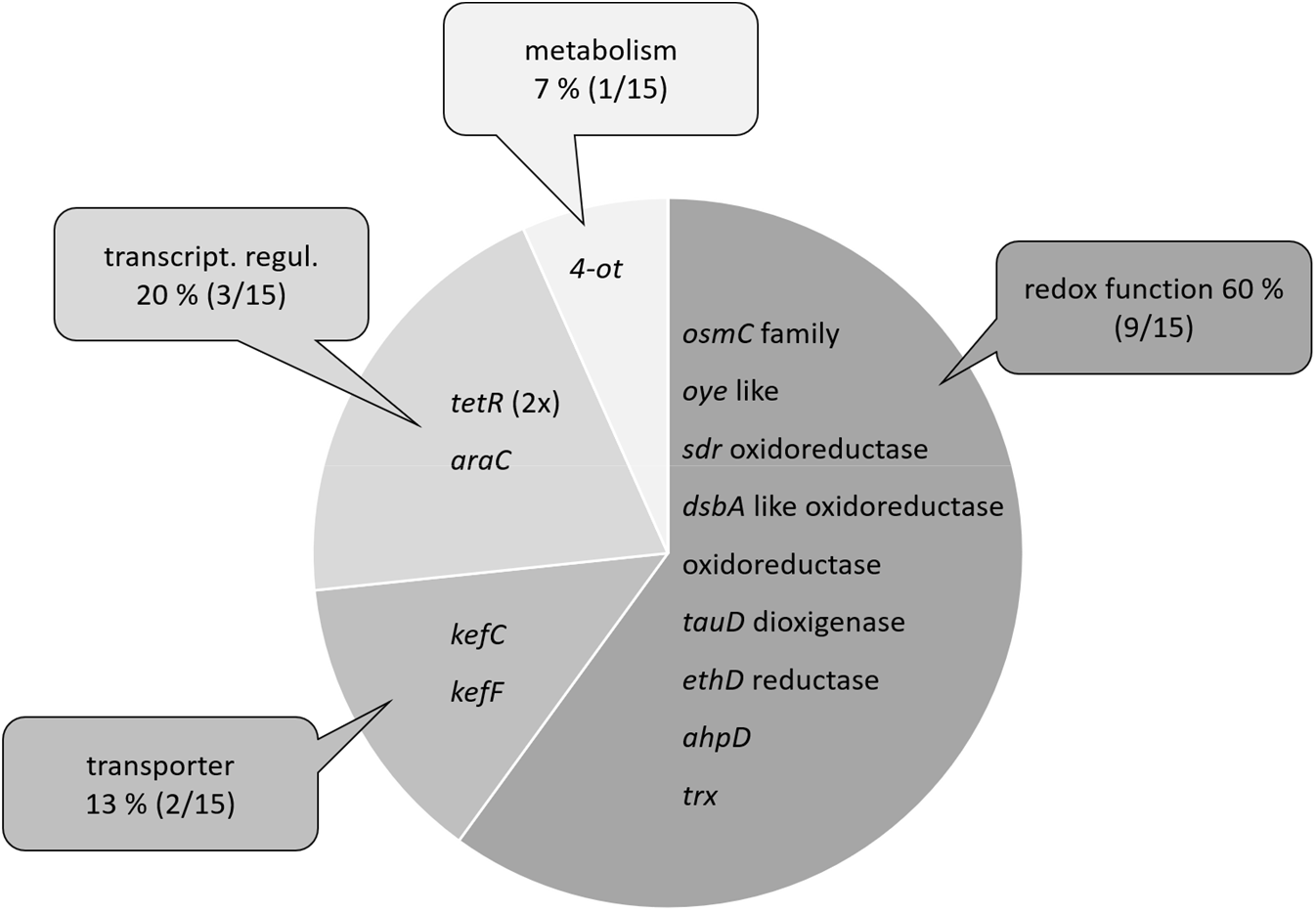
Functional classification of the congruent set of genes from the *Pf*AR-1 genomic repeats RE1, RE2, and RE3 on GI1, GI2 and GI3, respectively. 13 genes are present as a single copy per repeat but the *tetR* transcriptional regulator is present in duplicate on each repeat.

Genes on RE1 and RE2 appeared more closely related to each other than to those on RE3 from both a gene commonality (Jaccard similarity of 90% for RE1 vs RE2, compared to 54.2% for RE1 vs RE3, and 50% for RE2 vs RE3), and amino acid-similarity perspective (97.5%, 87.1% and 87.2%, respectively). This suggested that RE1 and RE2 originated from a more recent sequence duplication and that RE3 resulted from an earlier duplication event from common ancestor of RE1 and RE2.

### Comparison of putative RE regions across the *Pseudomonas* genus

In order to determine potential origins for the RE regions, we arranged these sequences to form a bait set, and compared this against all 3347 available *Pseudomonas* genomes. Similar regions to the bait were detected in 8 of the complete genomes, of which 6 were from plant-pathogenic or plant-associated pseudomonads. Matching regions were also found in 56 of the draft genomes, 8 of which showed two copies of the region. One of the draft genomes had the matching region split across two contigs, although this was presumably due to incomplete assembly, rather than representing a biological signal. These similar regions ranged from effectively complete, with hits from all 26 bait groups, to highly divergent with only 5 of the bait groups found. Of the 56 partial genome sequences, 37 were from plant-pathogenic or plant-associated bacteria (SM15)

### Inter-Species Codon Analysis

Expecting the codon usage of a horizontally transferred gene region to resemble the donor species rather than the current host, we performed a codon usage analysis to complement the bait-sequence analysis described above. For this, we compared for the full *Pf*AR-1 genome, the 3 RE regions, the 3347 other available *Pseudomonas* genomes, and 8 representative *non-Pseudomonas* Gammaproteobacteria. The results were plotted using Principal Component Analysis (PCA), and are shown in Fig. 9.

**Fig. 9.**
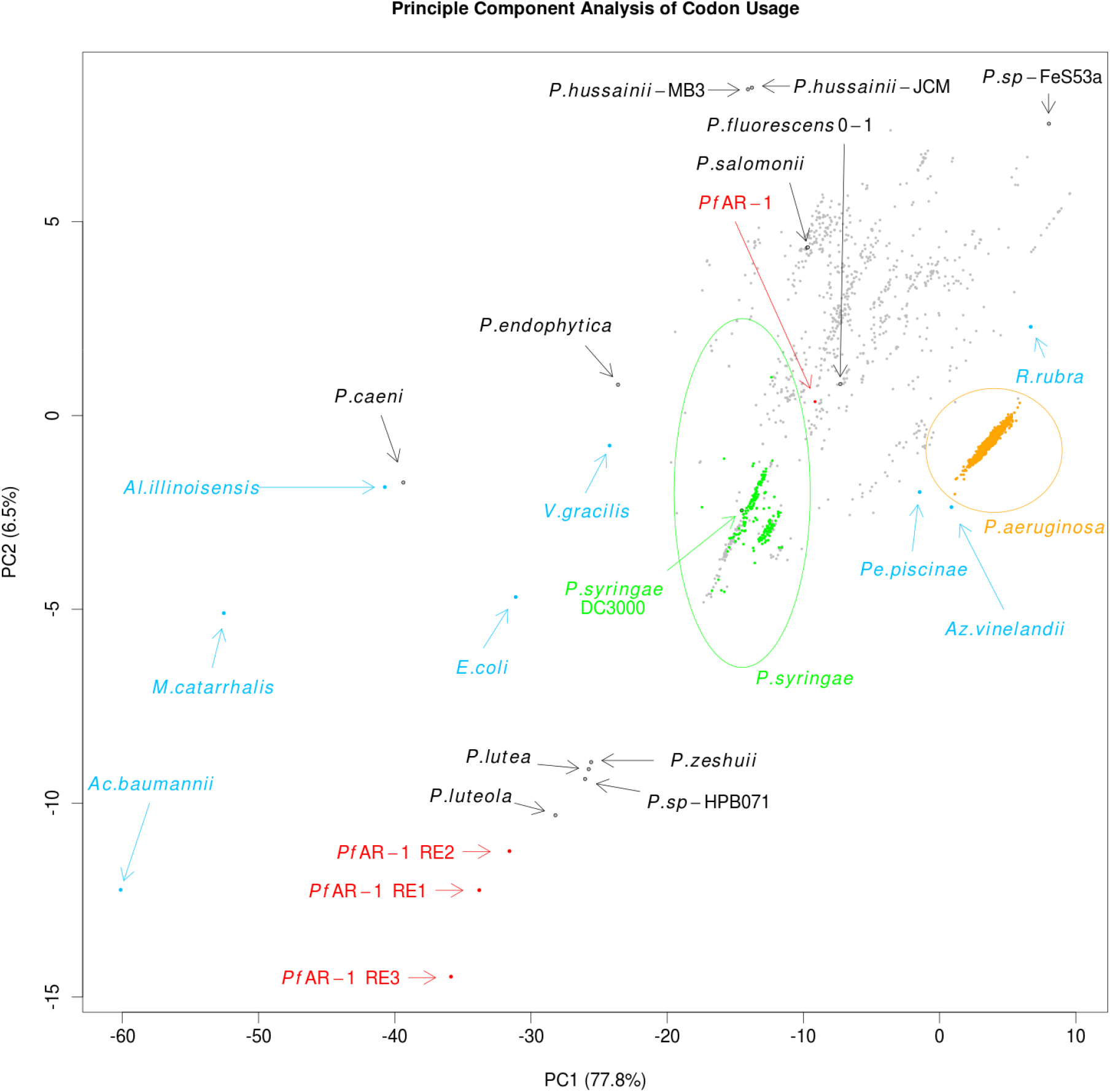
Principle Component Analysis of codon usage in REI, RE2 and RE3 of *Pf*AR-1 compared to the whole genome of *Pf*AR-1, 3347 pseudomonad genomes, and 8 Gammaproteobacteria. The putative HGT regions are clearly separated from the host *Pf*AR-1 genome.

The first principle component, which accounts for almost 78% of the variation, is consistent with GC content, ranging from *Acinetobacter baumannii* with a GC content of 38.9% on one extreme, to *Rugamonas rubra* with 67% GC content on the other, and unsurprisingly, given their usually low GC content, separates the putative HGT regions from not only the *Pf*AR-1 whole genome, but also from the vast majority of other *Pseudomonas* genomes. The second principle component also separates the putative HGT regions from the other genomes, although this component should not be over-interpreted since it accounts for only 6.5% of variation.

The resulting plot loosely clusters the 3 GI regions with 4 sequenced *Pseudomonas* species, namely *P. luteola, P. lutea, P. zeshuii* and *P*. sp. HPB0071. Unfortunately, none of these 4 species were found to contain matches for the bait sequences in the cross-species comparison above, and thus they are unlikely to be the origin of the putative HGT regions.

In addition to the genome-wide analysis, we also did a gene-window analysis of *Pf*AR-1, *Pf*0-1, *Pst*. DC3000, and *P. salomonii* ICMP 14252 (SM16).

### Phylogenetic Comparison of Whole Genome vs RE-like sequences

Regions which have been horizontally transferred have, by definition, an evolutionary history distinct from their host genomes. We therefore created a phylogenetic trees for the RE-like regions across the *Pseudomonas* clade, comprising the 3 RE regions from *Pf*AR-1, plus 72 RE-like regions from other species.

This was then compared to a whole-genome phylogenetic tree of 280 *Pseudomonas spp*. supplemented by 4 more distant genomes, namely *Azotobacter vinelandii* DJ, *Acinetobacter baumannii* AC29, *Escherichia coli* K12 MG1655 and *Burkholderia cenocepacia* J2315 which served as an outgroup. The 280 *Pseudomonas* genome subsets consisted of a) all 215 complete genomes, b) the 56 draft genomes showing a substantial hit against the Repeat Region bait set, as described above, and c) 9 *Pseudomonas* genomes with unusual codon usage (*P. lutea, P. luteola, P. sp* HPB0071, *P. sp* FeS53a, *P. zeshuii, P. hussainii* JCM, *P. hussainii* MB3, *P. caeni* and *P. endophytica*).

It is immediately apparent from comparison of the resulting region and whole genome trees that the RE-like regions have a distinct evolutionary history. For easier visualization of the whole genome trees, the 5 clades covering all species containing the genomic islands were manually split into sub-trees, and plotted separately. The top-level whole-genome tree, with each extracted clade reduced to a single node, is shown in SM17. The RE-like region tree for clade D is shown in Fig. 10 *A*, while the corresponding whole-genome tree is shown in Fig. 10 *B*. The trees for the remaining 4 clades (A, B, C, and E), are shown in SM17.

**Figure 10.**
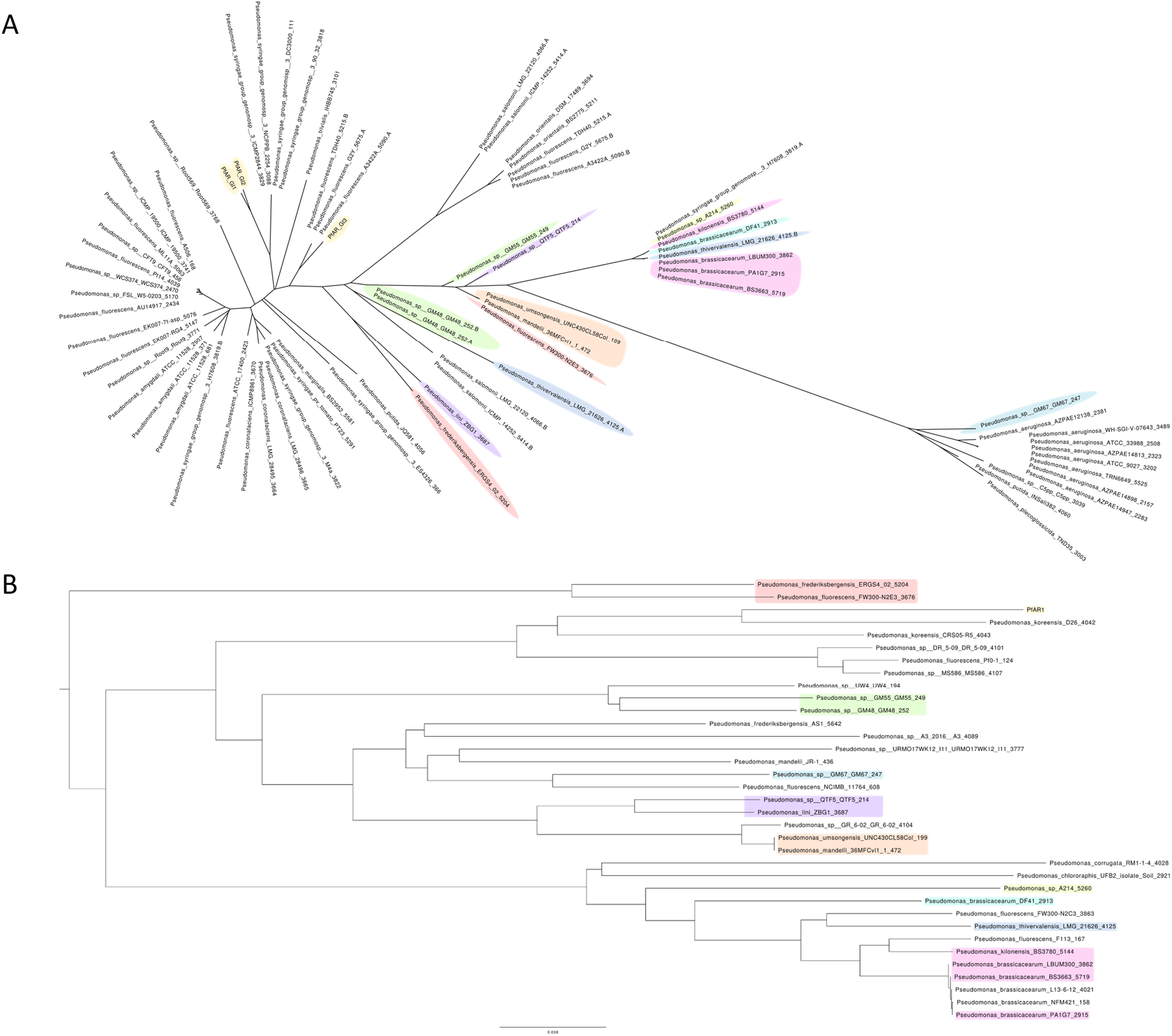
Comparison of RE-like region (*A*) and whole-genome phylogenetic (*B*) trees of a set of *Pf*AR-1 close relatives.

### IslandViewer Analysis

For independent confirmation of the above analysis, IslandViewer 4 (Bertelli et al., 2017) was used to assess the *Pf*AR-1 genome for horizontal gene transfer events. This analysis, shown in Fig. 11, also clearly identifies the 3 putative HGT regions, although additional weaker candidate regions are also indicated.

**Figure 11.**
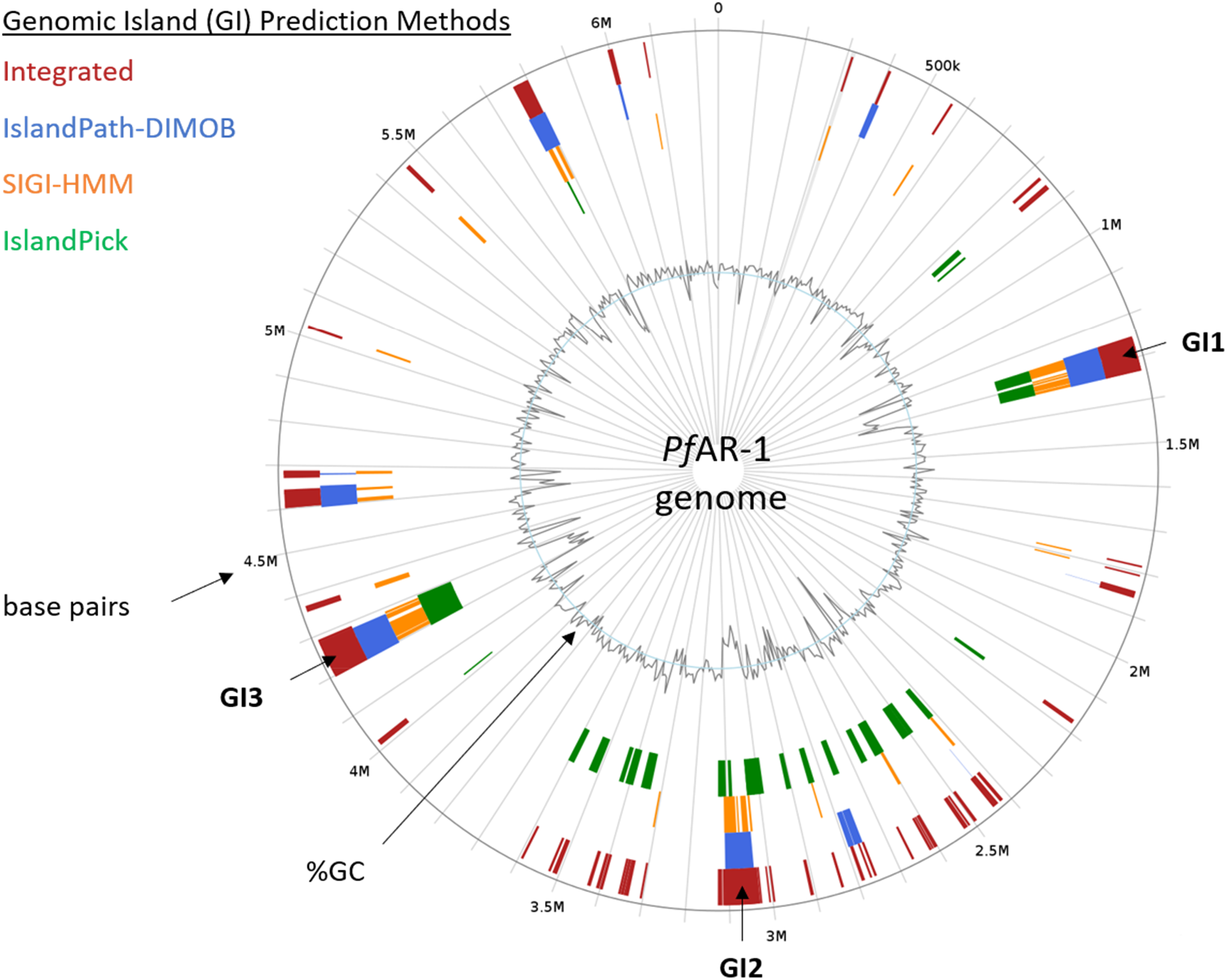
*In silico* prediction of genomic island in the *Pf*AR-1 genome. For *in silico* prediction, *Pf*AR-1 genome was checked with IslandViewer 4 (Bertelli et al., 2017) for genomic islands (GIs). Prediction is based on the three independet methods IslandPath-DIMOB (Hsiao et al., 2003), SIGI-HMM (Waack et al., 2006), and IslandPick (Langille et al., 2008). Results of all methods are integrated (dark red boxes) for island prediction. The position of GI1, GI2, and GI3 corresponds to the genomic regions with lowered GC content that contain the genomic repeat regions and the allicin resistance genes of *Pf*AR-1.

### Syntenic regions to *Pf*AR-1 REs *in P. syringae* pv. *tomato* DC3000 and *Pseudomonas salomonii* ICMP 14252

Data base comparisons revealed that some plant-associated bacteria, for example the garlic pathogen *P. salomonii* ICMP 14252 (Gardan et al., 2002), and tomato and *A. thaliana* pathogen *P. syringae* pv. *tomato* DC3000 (*Pst* DC3000) (Cuppels 1986; Buell et al. 2003), have regions syntenic with RE1 (Fig. 12). *Pf*AR-1 RE1 contains a *glr* gene and two gene groups (from *ahpD* → *kefC* and from *trx* ← *osmC*) that are conserved in RE2 and RE3. These two groups are present in the two syntenic regions in the genome of *P. salomonii* ICMP 14252 and in one syntenic region in *Pst* DC3000 (Fig. 12). In contrast, the French bean (*Phaseolus vulgaris*) pathogen Ps1448A has no genes with significant similarity to any of the allicin-resistance-conferring congruent gene set from *Pf*AR-1 clones. Ps1448A is fully sequenced (Joardar et al., 2005) and is quite similar at the nucleotide level to *Pst* DC3000 with an average nucleotide identity (ANI) of 86.87 %. In comparison, the ANI between *Pf*AR-1 and *Pf*0-1 is 85.94 % (SM18).

**Figure 12:**
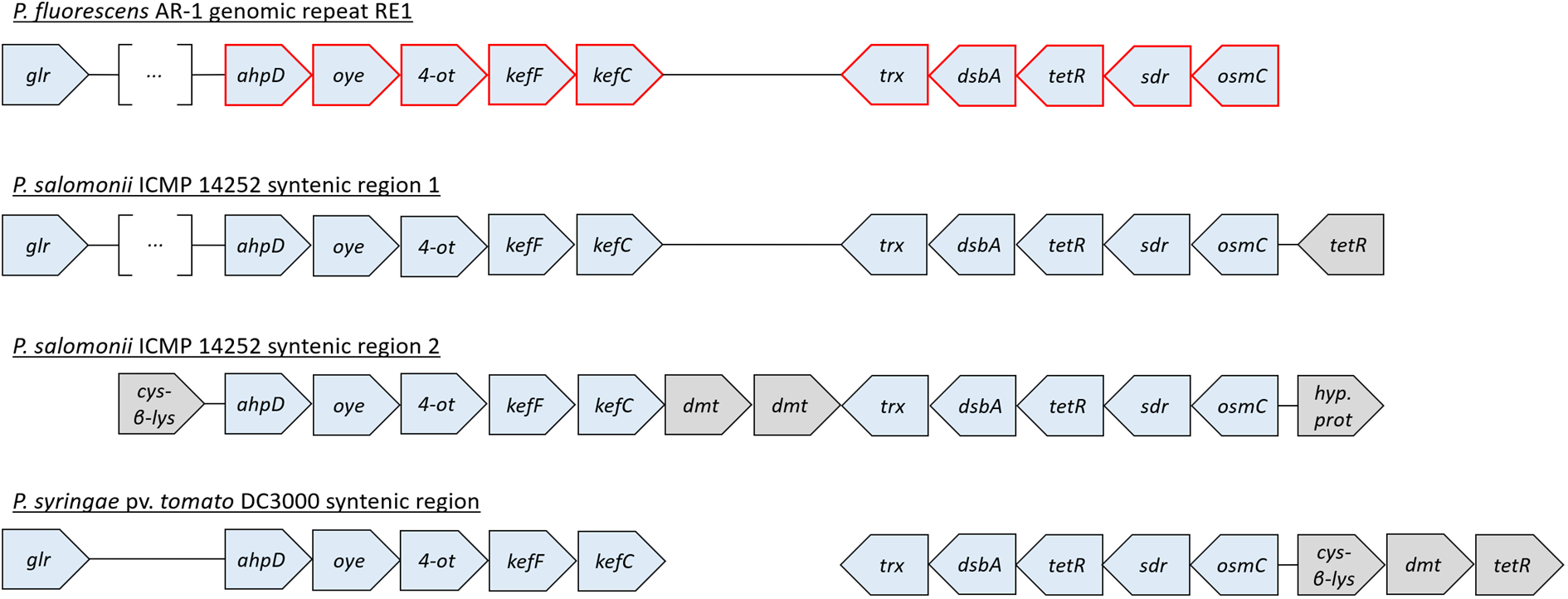
A set of ten genes is conserved in the genomic repeats of *Pf*AR-1 and in syntenic regions of *P. salomonii* ICMP 14252 and in *P. syringae* pv. *tomato* DC3000. *cys-β-lys:* cystathione-β-lyase; *dmt:* Permease of the drug/metabolite transporter (*dmt*) superfamily; the remaining genes are referred to elsewhere in this study. The distance between the different genes does not represent the actual intergenic distances since the gene blocks were graphically aligned to highlight the conservation. In case of *glr* of *Pf*AR-1 RE1 and *P. salomonii* syntenic region 1, these genes are farther upstream of the highlighted genes with several genes in between (represented by the squared brackets with three dots). Red highlighted genes represent the congruent set of genes also found in the resistance-conferring genomic clones of *Pf*AR-1. Coordinates of syntenic regions are given in SM19.

When *Pf*AR-1, *P. salomonii* ICMP14252, *Pst* DC3000 and Ps1448A were tested in a simple streak assay, we observed that *Pf*AR-1 and *P. salomonii* are most resistant against allicin, followed by *Pst* DC3000, then with a much higher susceptibility, by Ps1448A. The transfer of genomic clone 1 of *Pf*AR-1 which contained the core genes described above, raised the allicin resistance of Ps1448A to approximately the same level of allicin resistance observed in *Pst* DC3000 (Fig. 13).

**Figure 13:**
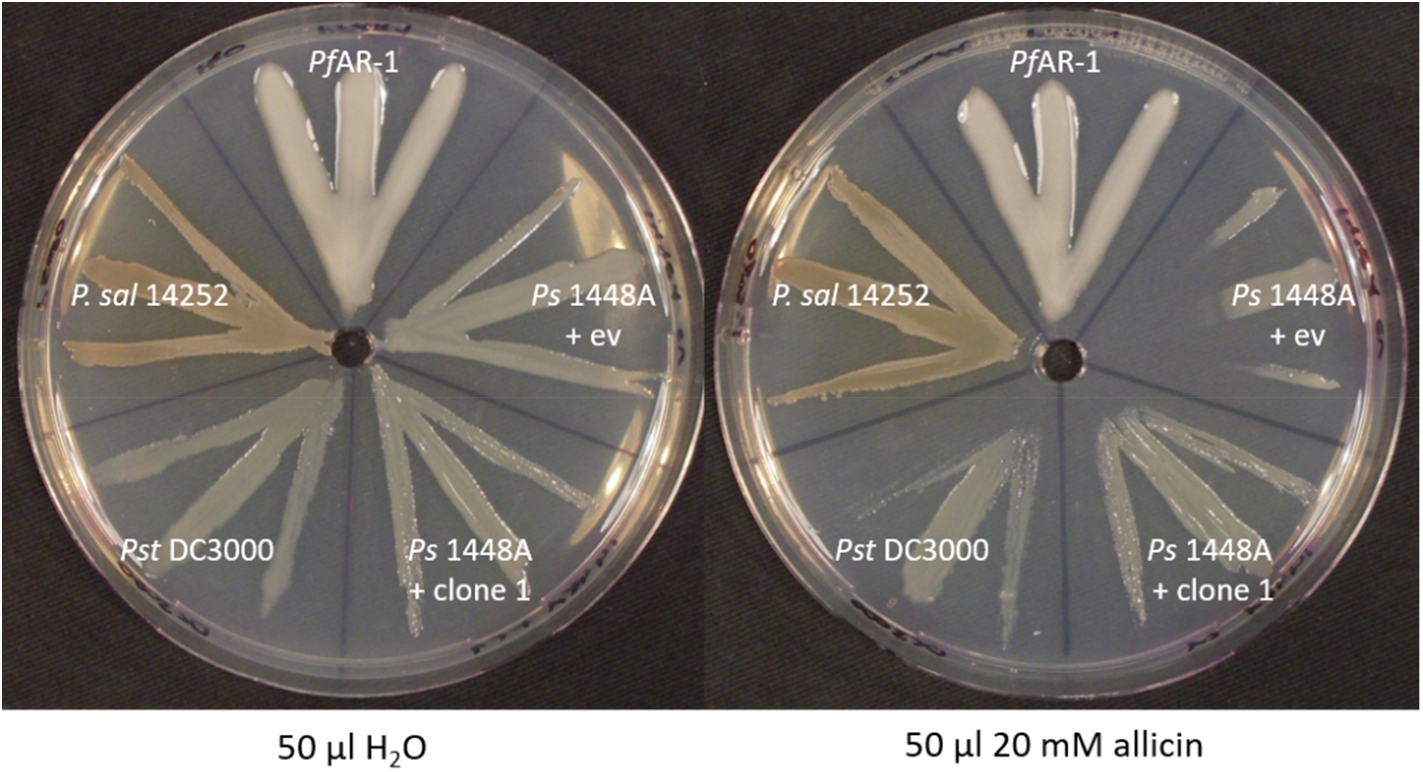
The allicin resistance of different bacteria correlates with the number of gene copies that are syntenic to the core fragment of the genomic clones from *Pf*AR-1. Ps1448 A was either transformed with *Pf*AR-1 genomic clone 1 or pRU1097 (empty vector control), while the other strains were not genetically modified. *Pf*AR-1 has three copies of a set of ten genes that were identified on genomic clones (e.g. genomic clone 1) that confer resistance to allicin in *P. syringae* strain 4612. *P. salomonii* ICMP 14252 has two copies of this set of genes in its genome while *P. syringae* pv. *tomato* DC3000 has one and *P. savastanoi* 1448A none.

### The role of glutathione reductase (Glr) in *Pf*AR-1 allicin resistance

Both GI1 and GI2 have a *glr* gene (*glr2*, *glr3*) outside of the allicin-resistance-conferring genomic clone regions in RE1 and RE2, respectively, and a further *glr* gene (*glr1*) is present on the *Pf*AR-1 chromosome. Furthermore, *Pf*AR-1 also had a two-fold higher basal Glr activity than *Pf*0-1 (Fig. 14). Since allicin targets -SH groups in proteins and GSH metabolism is critical for resistance to allicin, we investigated the potential contribution of Glr to allicin-resistance in *Pf*AR-1.

**Fig. 14.**
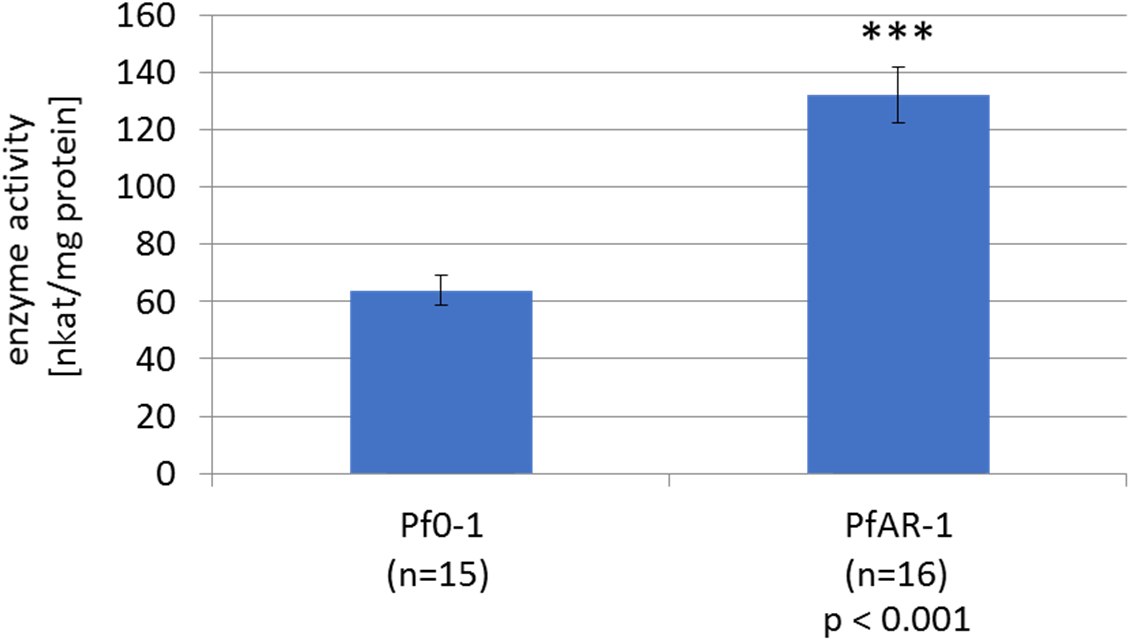
Glutathione reductase activity in protein extracts from *Pf*0-1 and *Pf*AR-1. Significance was tested with student’s t-test (P<0,001).

The importance of GSH metabolism and Glr for allicin-resistance is shown in Fig. 15. The agar-diffusion test showed that deleting the *glr* gene from *E. coli* BW25113 increased its susceptibility to allicin compared to the wt (Fig. 15 *A* and *B*). Allicin-resistance was restored by complementing the BW25113 Δ*glr* strain with the chromosomal *glr1* gene from *Pf*AR-1 (Fig. 15 *C*).

**Figure 15.**
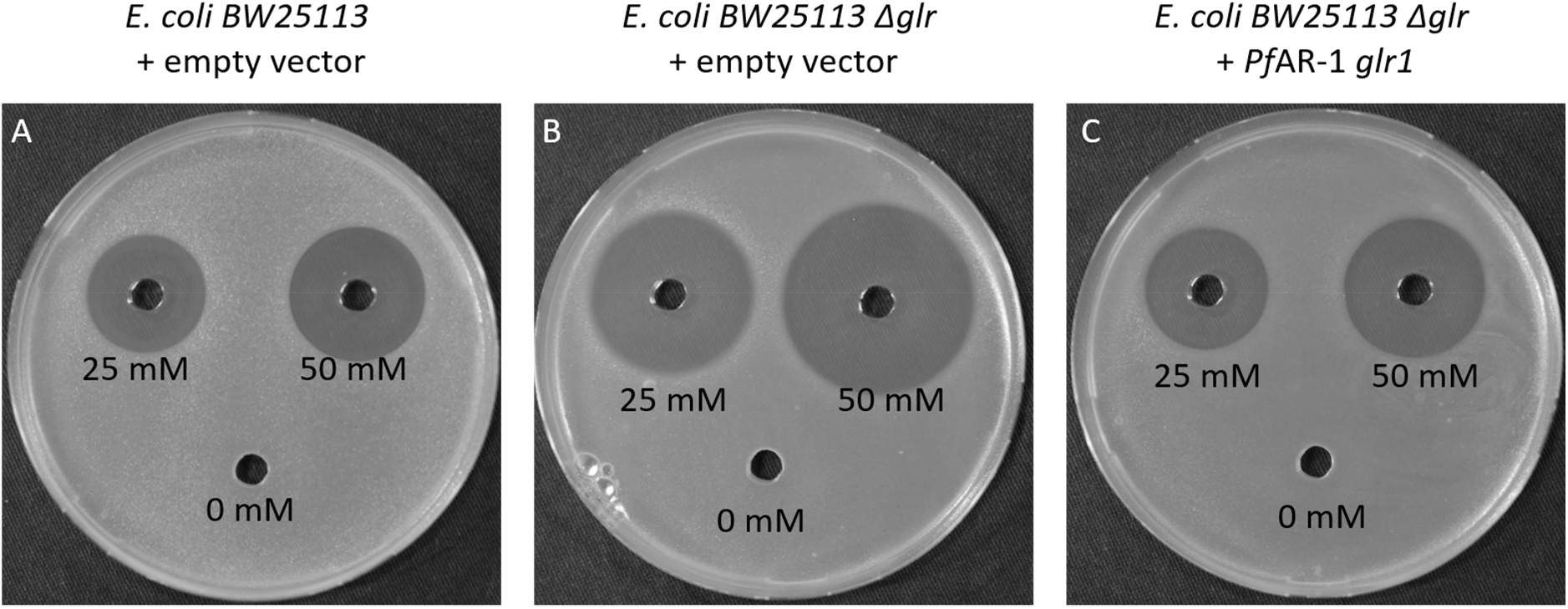
*Pf*AR-1 glutathione reductase (*glr1*) complements *E. coli* BW25113 glutathione reductase deletion mutant (*Δglr*). 40 μL of allicin solution (or water) were pipetted into wells in *E. coli*-seeded medium and the plates incubated over-night at 37°C.

## Discussion

The garlic defense substance allicin is a potent thiol reagent which targets accessible -SH groups in proteins and the cellular redox buffer glutathione (Borlinghaus et al., 2014). Allicin has been shown to *S*-thioallylate several cysteine-containing proteins in bacteria (Müller et al., 2016, Loi et al., 2019), and has been described as a redox toxin (Gruhlke et al., 2010). *S*-thioallylation by allicin is reversible, and sublethal doses suppress bacterial multiplication for a period of time, the length of which is dose-dependent, before growth resumes (Müller et al., 2016). Adding a lethal dose of allicin to a high-density bacterial culture and plating out for survivors, the usual strategy to isolate antibiotic-resistant mutants, has proven ineffective with allicin. Because allicin affects such a broad catalogue of cellular proteins, it is not easy for an organism to adapt to it by simple mutation. Nevertheless, the degree of allicin resistance varies between different bacterial isolates and, although the genetic basis for this variation was unknown, we reasoned that we might find organisms with high allicin-resistance associated with the garlic bulb itself as a niche-habitat. This was indeed the case, and we were able to isolate a highly allicin-resistant bacterium, which we named *P. fluorescens* Allicin Resistant-1 (*Pf*AR-1), from a garlic bulb. In inhibition zone tests, comparison with *E. coli* K12 DH5α or *P. syringae* 4612, *Pf*AR-1 showed an exceptionally high degree of allicin-resistance (Fig. 1). We employed parallel approaches of functional testing of random genomic clones, and whole genome sequencing, to characterize genes conferring allicin-resistance to *Pf*AR-1 (Figs 2 & 3). Interestingly, positive clones conferred allicin-resistance not only to closely-related pseudomonads, but also to distantly related bacteria such as *E.coli* (Figs 4 & 5).

Resistance-conferring *Pf*AR-1 genomic clones contained 16 unique genes in total, with a congruent set of 8 genes shared by all clones (Fig. 3). Since allicin is a redox toxin causing oxidative stress, it was interesting to observe that half of these genes were annotated with redox-related functions (Fig. 3). Moreover, these genes were linked to reports in the literature in the context of oxidative and disulfide stress (Table 1).

Transposon mutagenesis indicated that the *dsbA, trx, kefC*, and *oye* genes worked together, contributing incrementally to confer allicin-resistance to a susceptible recipient. The contributions of the *dsbA* and *trx* genes were greater than those of the *kefC* and *oye* genes. In contrast, the effect of a mutation in *ahpD* alone was major, with transposon mutants showing a similar phenotype to the susceptible parent (Fig. 5). These results are consistent with a multicomponent mechanism of allicin resistance. Since the genes involved do not have annotated genes that code for significantly similar peptides in the *Pf*0-1 reference strain, or *E. coli*, this suggests that the genes might be of external origin and would explain why spontaneous mutation to a gain-of-resistance upon allicin selection was not observed in axenic cultures under lab conditions.

To investigate individual contributions to the resistance phenotype, genes were cloned in a broad-host-range overexpression vector and investigated in *Ps*4612. We observed a strong increase in allicin resistance by the overexpression of *ahpD* or *dsbA* alone, with either gene conferring almost as much resistance as the complete genomic clone (Fig 6). However, we observed no effect for *trx* or *oye* overexpression; although transposon insertions in the complete genomic clone led to a decrease in resistance (Fig. 5). This might indicate that the function of the single gene depends on the function of another gene or genes from the genomic fragment, or that there are downstream effects of the Tn-insertion. Overexpression lines for *osmC* and *kefC* were not found in *Ps*4612, most likely due to toxic effects. In this regard, the activity of KefC is normally tightly regulated by KefF and GSH, and an imbalance could lead to a toxic decrease in cellular pH and loss of potassium, which is needed to maintain turgor and enable cell growth and division (Epstein, 2003). Overexpression of *sdr* and *oye* showed no phenotype (not shown).

In parallel to the gain-of-function approach, genome analysis revealed unique features of the *Pf*AR-1 genome compared to the *Pf*0-1 reference strain. Thus, three large genomic regions (GI1 to GI3, Fig. 7) were identified, with sizes between 79 to 98 kbp, having a lower GC content (Δ%GC approximately 5-10 %). These GI’s contained repeat regions RE1, RE2 and RE3 with the resistance-conferring clones identified in the functional analysis being contained within them (Fig. 3 *B*, 8).

A codon usage analysis showed differences in RE1, RE2 and RE3 compared to the core *Pf*AR-1 genome (Fig. 9); which was a strong indication that these regions were obtained by horizontal gene transfer (HGT), and comparison with other *Pseudomonas* spp. suggested that the origin may lie outside this genus. The HGT hypothesis was strongly supported by our phylogenetic analysis (Fig. 10, SM17), and an independent *in silico* analysis using IslandViewer 4 (Fig. 11). Thus, by current selection criteria, regions RE1, RE2 and RE3, and most likely the complete GI1, GI2 and GI3 regions can be reliably considered to be *bona fide* genomic islands which were obtained by horizontal gene transfer. The preponderance of genes with redox-related functions in the RE regions fitted well with a role in resistance against allicin. It is unusual for multiple copies of genes to be maintained in bacteria because of the genomic instability that arises through homologous recombination leading to genome rearrangements and loss of essential interim sequences (Rocha, 2003). The presence of such large, widely spaced REs in the *Pf*AR-1 genome, suggested that there was a high selection pressure to maintain them. The latter is presumably associated with the allicin-resistance-conferring functions of many of the genes and this fits with the competitive advantage they offer over other bacteria in occupying the garlic niche.

Although the GI donor remains unknown, phylogenetic analysis identified similar syntenic regions to the REs from *Pf*AR-1 in other bacterial genomes (Fig. 12). Thus, the garlic pathogen *P. salomonii* ICMP14252 has two syntenic regions, and the well-described model pathogen *P. syringae* pv. *tomato* DC3000 has one syntenic region. In *P. salomonii* ICMP 14252 and *Pst*. DC3000 the syntenic regions have the set of ten core genes identified in genomic clones 1-7 of *Pf*AR-1 (Fig.12). Furthermore, we observed that the species with multiple copies of the syntenic regions, 3 for *Pf*AR-1 and 2 for *P. salomonii*, showed higher allicin resistance than those with only one or zero copies, namely *Pst* DC3000 and *Ps* 1448A, respectively (Fig 13). This further supports our findings that these regions are putative allicin resistance factors in *Pf*AR-1. *P. salomonii* causes the café-au-lait disease on garlic (Gardan et al., 2002) and its high degree of allicin resistance corresponds well to its niche as a pathogen of garlic. One might expect that a pathogen like *P. salomonii* could be the origin of allicin resistance genes in *Pf*AR-1, but according to our codon usage analysis, the allicin resistance regions in *P. salomonii* ICMP14252 are quite distinct from the remainder of the genome, and therefore were also likely obtained by horizontal gene transfer (Fig 9, 10). *Pst* DC3000 is a model pathogen with a fully sequenced genome (Buell et al., 2003) and is pathogenic on tomato and on the model organism *Arabidopsis thaliana* (Xin and He, 2013). To the best of our knowledge, the genes and their function in allicin resistance have not been described before in this well studied strain. Although our experiments suggest that the resistance conferred by the core-region is allicin-specific (Fig. 4), oxidative stress has manifold causes and some genes in the syntenic region may help to counter its effects. In this regard, it was reported that a transposon insertion in *dsbA* from the core genome of *Pst* DC3000 led to decreased virulence of *Pst* DC3000 on *A. thaliana* and on tomato (Kloek et al., 2000). Based on this study, it seemed that the remaining *dsbA* copy from the syntenic region of *Pst* DC3000 was not sufficient to functionally complement the loss of the *dsbA* in the core genome; perhaps indicating subtly different functions between the two. It is intriguing to speculate that the syntenic region might help to overcome the oxidative burst associated with plant defense, as well as protecting against more specifically redox-active sulfur-containing plant defense substances, like allicin. The oxidative burst in plants is a general defense response to avirulent pathogens (Lamb & Dixon, 1997), and it would be interesting to see if loss of syntenic genes other than *dsbA* in *Pst* DC3000 also lead to a reduction of virulence. Moreover, a recent study reported plasmid-born onion virulence regions (OVRs) in different *Pantoea ananatis* strains that are pathogenic on onion (Stice et al., 2018). The OVR-A region contained a subset of genes that we describe in our present study as allicin-resistance genes. More specifically, *dsbA*, which was annotated in *P. ananatis* as *isomerase* in OVR-A, *oye* (as *alkene reductase), trx, ahpD* (annotated as *alкylhydroperoxidase*), *glutathione disulfide reductase, sdr*, and *osmC*, were all present. Although onion does not produce allicin, it has many sulfur-containing redox-active compounds which may be involved in defense (Imai et al., 2002). Nevertheless, there are several plant pathogenic bacteria, e.g. *P. savastanoi* 1448A, which have no equivalent syntenic region but are successful plant pathogens in their own right. Therefore, there is clearly no absolute requirement for the syntenic region to enable colonization of plants as a habitat *per se*. However, in this regard it should be noted that a comprehensive genomic analysis of plant-associated bacteria to identify protein domains associated with adaptation to growth in or on plants, showed that 7 of the 10 genes we identified in the syntenic region contained plant-associated domains as described by the authors (Levy et al., 2018). Furthermore, of the bacteria showing homology to the syntenic region, 43/64 were from either plant-pathogenic or plant-associated bacteria (SM15). This observation is compatible with a general function of the syntenic region supporting colonization of the plant environment, perhaps acting favorably in an oxidative stress situation, in addition to a specific role in resistance to allicin and other sulfur-containing defense compounds.

Allicin targets *inter alia* the GSH pool in plants, and GSH metabolism has been shown to be important in the resistance of bacteria, yeast and *Arabidopsis thaliana* to allicin (Müller et al., 2016; Gruhlke et al. 2010; Leontiev et al., 2018). Thus, it was interesting to note that *Pf*AR-1 has three copies of *glr*, one each on RE1 and RE2, and one in the core genome. This is quite remarkable since bacteria normally have only one *glr* copy. Exceptions, like *Pst* DC3000 and *P. salomonii* ICMP14252, have an additional *glr* gene that was also very likely obtained by horizontal gene transfer as in *Pf*AR-1. Here, we demonstrated that the higher copy number of *glr* in *Pf*AR-1 correlated with a 2-fold higher basal GLR enzyme activity compared to *Pf*0-1 with only one copy of *glr* (Fig. 14). The importance of Glr activity for tolerance to allicin is clear from the enhanced sensitivity to allicin of a *Δglr* knockout in *E. coli*. Moreover, we were able to complement this phenotype by introducing *glr1* from *Pf*AR-1 (Fig. 15). Glr recycles oxidized glutathione (GSSG) to GSH using NADPH as a reductant. GSH protects cells from oxidative stress, either by direct reaction with pro-oxidants, thus scavenging their oxidative capacity, or by serving as an electron donor for detoxifying enzymes like glutathione peroxidase and glutaredoxins (Meister and Anderson., 1983). It was shown that allicin treatment leads to oxidation of GSH to GSSG in yeast (Gruhlke et al., 2010), and to the formation of *S*-allylmercaptoglutathione (GSSA) (Horn et al., 2018). In yeast, GSSA is reduced by Glr to release GSH, shown by *in vitro* activity measurements with GSSG and GSSA as substrates, and *in vivo* by chemical complementation of the *Δgsh1* phenotype of yeast cells with GSSA (Horn et al., 2018). Therefore, it is clear that Glr is an important resistance factor for *Pf*AR-1 to allicin and this fits with the HGT hypothesis allowing *Pf*AR-1 to exploit its ecological niche. Gram-positive bacteria, such as *Staphylococcus aureus*, have bacillothiol rather than GSH and in an independent investigation we showed that the bacillothiol reductase YpdA, which is the functional equivalent of Glr, reduced *S*-allylated bacillothiol (BSSA) and is important for the resistance of *Staphylococcus* to allicin (Loi et al., 2019). Furthermore, GSH negatively regulates the activity of KefC but GSH-conjugates stimulate KefC activity via KefF (Fig. 16) (Miller et al., 2000, Ferguson et al., 1997). Thus, GSH inhibits K^+^-efflux and *E. coli Δgsh* mutants lose K+-ions similarly to cells stressed with electrophiles such as *N*-ethylmaleimide (NEM) (Meury & Kepes 1982; Elmore et al., 1990). KefC activity acidifies the cytoplasm and protects against oxidative stress caused by electrophiles such as NEM and methylgloxal; presumably because the lowered pH works against thiolate ion formation (Ferguson 1993, 1995, 1996, 1997; Poole, 2015). KefC activation would be expected to protect against oxidative stress caused by the electrophile allicin in the same way (Fig. 16).

**Figure 16:**
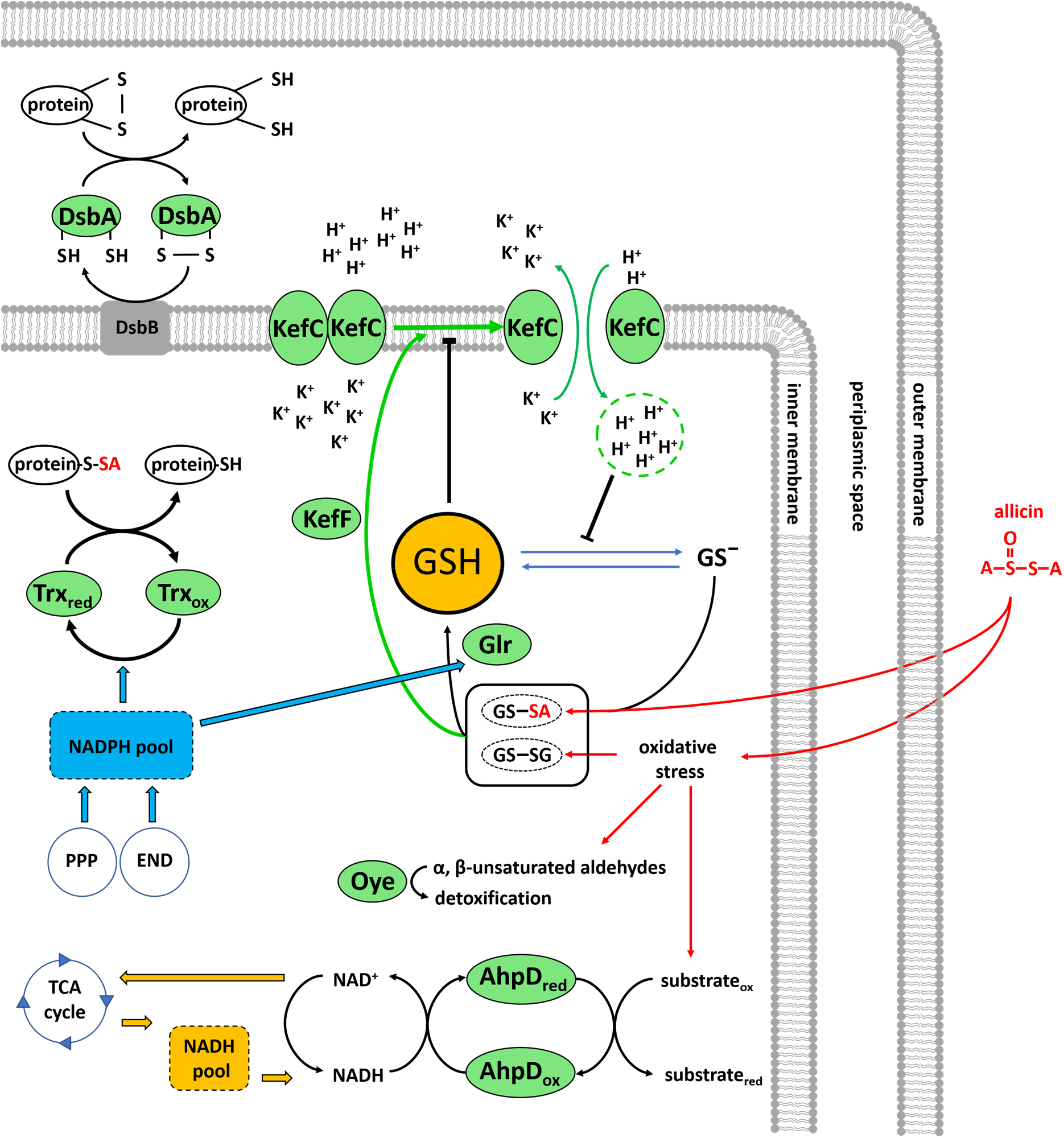
Suggested model for allicin resistance in *Pseudomonas fluorescens* Allicin Resistant-1. Disulfide bond protein A (DsbA), thioredoxin (Trx), glutathione (GSH), *S*-allylmercaptoglutathione (GSSA), Glutathione reductase (Glr), glutathione-regulated potassium efflux system (KefF, KefC), alkylhydroperoxidase D (AhpD), pentose phosphate pathway (PPP), Entner/Doudoroff pathway (END).

Because *Pf*AR-1 lacks the *6-phospho-fructokinase* gene necessary for glycolysis, it uses the Entner Dodouroff Pathway (EDP) to metabolize glucose to pyruvate. During oxidative stress, the EDP is advantageous over glycolysis because, in addition to NADH, NADPH is produced (Conway, 1992). For example, it was shown for *P. putida* that key enzymes of the EDP are upregulated upon oxidative stress (Kim et al., 2008). NADPH is used as reducing equivalents for antioxidative enzymes like glutathione reductase and Oye-dehydrogenases. Thus, cells using the EDP have a further source of NADPH in addition to the pentose phosphate pathway. Moreover, in *Mycobacterium tuberculosis* the AhpD enzyme depends on NADH consumption (Bryk et al., 2002), and thus, *Pf*AR-1 could be able to tap into two pools of reducing equivalents to defend against allicin stress (Fig. 16).

Disulfide bond protein A (DsbA) is located in the periplasm (Shouldice et al., 2011), and based on its protein domain content, in *Pf*AR-1 DsbA might act as disulfide isomerase or as a chaperone. The Dsb system might be supported with reducing equivalents from the cytosol by thioredoxin (Trx) and additional membrane proteins as previously reported for *E. coli* (Katzen and Beckwith, 2000).

How alkylhydroperoxidase D (AhpD) might protect *Pf*AR-1 against allicin is so far unknown. Possibly, as in *Mycobacterium tuberculosis*, it might act by using NADH to reduce oxidized molecules arising from oxidative stress (Bryk et al., 2002).

All these data reinforce the central importance of GSH metabolism and redox enzymes in the resistance of cells to the electrophilic thiol reagent allicin. The maintenance of multiple copies of resistance genes, obtained by HGT, probably facilitates exploitation of the garlic ecological niche by *Pf*AR-1 in competition with other bacteria.

## Acknowledgements

Financial support from the RWTH Aachen University (J.B., A.J.S., MCHG,) is gratefully acknowledged. J.B. was supported by an RFwN Ph.D. stipendium. This research did not receive any specific grant from funding agencies in the public, commercial, or not-for-profit sectors. Nikolaus Schlaich ist thanked for helpful discussions and Ulrike Noll for proof-reading the MS.

