## supplementary material for "Plant-microbe co-evolution: allicin resistance in a *Pseudomonas fluorescens* strain (*Pf*AR-1) isolated from garlic"

### [SM1] Bacteria used in this study

Table S1: List of bacteria used in this study for physiological or genetic experiments.

| organism | reference |
|---|---|
| *E. coli* K12 HB101 | (Boyer and Roulland-Dussoix, 1969) |
| *E. coli* K12 MegaX DH10B $T1^R$ | Thermoscientific |
| *E. coli* K12 DH5α | (Hanahan, 1983) |
| *E. coli* S17-1 λpir | (Simon et al., 1983) |
| *E. coli* MG1655 | (Blattner et al., 1997; Neidhardt and Curtiss, 1996) |
| *E. coli* BW25113 | (Baba et al., 2006; Datsenko and Wanner, 2000) ; Coli Genetic Stock Center Number CGSC#: 7636; Strain BW25113 |
| *E. coli* BW25113 *Δglr* | (Baba et al., 2006); Coli Genetic Stock Center Number CGSC#: 10569; Strain JW3467-1 |
| *Pseudomonas fluorescens* Allicin resistant-1 (*Pf*AR-1) | this study |
| *Pseudomonas syringae* pv. *phaseolicola* strain 4612 | Robin E. Mitchell, Division of Horticulture and Processing, Department of Scientific and Industrial Research, Auckland, New Zealand |
| *Pseudomonas syringae* pv. *tomato* DC3000 | (Buell et al., 2003; Cuppels, 1986) |
| *Pseudomonas savastanoi* pv. *phaseolicola* strain 1448A | (Joardar et al., 2005) |
| *Pseudomonas salomonii* ICMP 14252 | (Gardan et al., 2002) |

### [SM2] Cultivation Methods and Media

**2xYT medium** is a more nutrient-rich version of Lysogeny Broth (LB) full medium (Bertani, 1951) and is frequently used for *E. coli* cultivation.

**King´s B medium** was originally described to cultivate *Pseudomonas* on full medium to search for fluorescent isolates (King et al., 1954). In contrast to the original recipe, $MgSO_4$ was left out of the King´s B medium in this study.

**M9JB medium** was developed during this study for the cultivation of *Pseudomonas* for reduced slime production. This defined medium is based on M9 salts (Maniatis, 1982) with glycerol as carbon source. Additionally, Nitsch vitamin mixture (Duchefa Biochemie, product N0410) are added to complement for *E. coli* auxotrophies, and complete supplement to enrich the media for amino acids (except cysteine) to improve doubling time. The detailed composition can be found in Table S2.

Table S2: Composition of M9JB medium.

| | ingredients | final concentration |
|---|---|---|
| 1x M9 minimal salts | $Na_2HpO_4$ | 33.9 g/l |
| | $KH_2PO_4$ | 1.5 g/l |
| | NaCl | 2.5 g/l |
| | $NH_4Cl$ | 5 g/l |
| | adjusted with NaOH to: | pH 7.5 – pH8 |
| | $MgSO_4$ (added at 50°C after autoclaving) | 2 mM |
| carbon source | glycerol | 1.25% (w/v) |
| 3x Complete Supplement Mixture (CSM) | adenine | 30 mg/l |
| | L-arginine | 150 mg/l |
| | L-aspartic acid | 240 mg/l |
| | L-histidine HCl | 60 mg/l |
| | L-isoleucine | 150 mg/l |
| | L-leucine | 300 mg/l |
| | L-lysine HCl | 150 mg/l |
| | L-methionine | 60 mg/l |
| | L-phenylalanine | 150 mg/l |
| | L-threonine | 300 mg/l |
| | L-tryptophan | 150 mg/l |
| | L-tyrosine | 150 mg/l |
| | uracil | 60 mg/l |
| | valine | 420 mg/l |
| | L-proline | 250 mg/l |
| 1x nitsch vitamin mixture (added at 50°C) | biotin | 0.05 mg/l |
| | folic acid | 0.50 mg/l |
| | glycine | 2.00 mg/l |
| | myo-Inositol | 100.00 mg/l |
| | nicotinic acid | 5.00 mg/l |
| | pyridoxine HCl | 0.50 mg/l |
| | thiamine HCl | 0.50 mg/l |
| (for solid medium) | agar | 1.5 % (w/v) |

### [SM3] Plasmids used in this study

Table S3: Plasmids used in this study

| plasmid | comments | source/reference |
|---|---|---|
| pSCR001 | contains the transposon IS-Ω-km/hah for transposon mutagenesis | (Giddens et al., 2007) |
| pRU1097 | promoter probe vector, broad host range, used to make genomic library from *Pf*AR-1 | (Karunakaran et al., 2005) |
| pJP2neo | modified version of pJP2 (Prell et al., 2002) source of neo-promoter for pJABO | Dr. Jürgen Prell, RWTH-Aachen, Unit of Sil Ecology, BioI |
| pBluescript I KS (-) | standard high copy cloning vector for *E. coli*. Source of MCS used to construct pJABO and pJABO5 | Stratagene Inc., La Jolla |
| pJABO | broad host range expression system, based on pRU1097 backbone | this study |
| pRS426 | yeast high copy expression vector (2µ ori, URA3). Used for construction of pJABO5. | (Mumberg et al., 1995) |
| pJABO5 | broad host range expression system, based on pRU1097 backbone | this study |

### [SM4] PCR and Primer

All DNA cloning steps in this work were based on enzymatic restriction and sticky end (or blunt end) DNA ligation with T4-DNA ligase from Thermoscientific. The necessary restriction sites for PCR fragments were introduced during PCR via primer overhangs if not already present in the DNA template.

For all PCR applications, the Phusion High-Fidelity PCR Master Mix (Thermo Scienctific) was used according to the user manual. All primer sequences are listed in Table S4.

Table S4: Primers used in this study

| Primer | sequence (5'->3') | amplicon | reference |
|---|---|---|---|
| CEKG 2A | GGCCACGCGTCGACTAGTACNNNNNNNNNNAGAG | random | (Manoil, 2000) |
| CEKG 2B | GGCCACGCGTCGACTAGTACNNNNNNNNNNACGCC | random | (Manoil, 2000) |
| CEKG 2C | GGCCACGCGTCGACTAGTACNNNNNNNNNNGATAT | random | (Manoil, 2000) |
| CEKG 4 | GGCCACGCGTCGACTAGTAC | nested primer for CEKG amplicons | (Manoil, 2000) |
| hah-1 | ATCCCCCTGGATGGAAAACGG | IS-Ω-Km/hah h | (Manoil, 2000) |
| P137 | ACTGACGCGGCCGCTTATTTGAGCGATGCAACTGCGTC | hypothetical protein | this study |
| P138 | ATATACTCTAGACGGTAAACATAGCCCACAGCGAAG | hypothetical protein | this study |
| P139 | ACTGACGCGGCCGCTTAGCCCTCGATAACTTCGAGCTT | *osmC* | this study |
| P140 | ATATACTCTAGAGCTTCGCTGTGGGCTATGTTTACC | *osmC* | this study |
| P141 | ACTGACGCGGCCGCTTAGGCGCGTACAGCCATCACCCC | *sdr* | this study |
| P142 | ATATACTCTAGAATTGTTGGTCGTCGGCGGCACAAG | *sdr* | this study |
| P143 | ACTGACGCGGCCGCTTATCCTGCACGGAGGAGTGAAGA | *tetR* | this study |
| P144 | ATATACTCTAGAGTGAGAAAGGGTGCTAGCTGTAGG | *tetR* | this study |
| P145 | ACTGACGCGGCCGCTCAAATTTTGCAGCCATCGATGCC | *dsbA* | this study |
| P146 | ATATACTCTAGAGCCGCAATGCAATCTTCACTCCTC | *dsbA* | this study |
| P147 | ACTGACGCGGCCGCTTAGCTAGCATCTTCCAGCTCGGT | *trx* | this study |
| P148 | ATATACTCTAGAGATAGCTCATTGATGGGCCAAGTC | *trx* | this study |
| P149 | ACTGACGCGGCCGCTCAGCTCCCTGTATCTCTATGCGC | *kefC* | this study |
| P150 | ATATACTCTAGACCGAAATTATGGCGAGCGCCTGACG | *kefC* | this study |
| P151 | ACTGACGCGGCCGCTCAATCTTCCCTCCATGACGTCAG | *kefF* | this study |
| P152 | ATATACTCTAGAGGCTTCTCCTTCAGGTCATTAG | *kefF* | this study |
| P153 | ACTGACGCGGCCGCTTACTGGCCGGCTTGCAGCTTCAA | *4-ot* | this study |
| P154 | ATATACTCTAGAAGCGGTTCGATTGATAAGGGCTAC | *4-ot* | this study |
| P155 | ACTGACGCGGCCGCTCAGGCTTTGGCAGCTGCGTATTC | *oye* | this study |
| P156 | ATATACTCTAGAGACGCGTTCGACAAGTCCTAAATC | *oye* | this study |
| P157 | ACTGACGCGGCCGCTTAGGACTTGTCGAACGCGTCGAG | *ahpD* | this study |
| P158 | ATATACTCTAGACTGCAGGGTGTTAAAGCCCACGTGAG | *ahpD* | this study |
| P159 | ATATACGTGCACCCGGAATTGCCAGCTGGGGCGCCC | Neo prom. | this study |
| P160 | ACTGACCTCGAGATATACCGATCGTAAACGGCTAGCTTGCAGGGCTTCCCAACCTTACCA | Neo prom. | this study |
| P161 | ATATACCGATCGGAGCTCCACCGCGGTGGCGGCCGC | part of *lacZ* with MCS | this study |
| P162 | ATATACGCTAGCGGTACCGGGCCCCCCCTCGAGGTC | part of *lacZ* with MCS | this study |
| P163 | ACTGACGTGCACTATGCTTGTAAACCGTTTTGTGAA | pRU1097 backbone | this study |
| P164 | ACTGACCTCGAGGCTCTACAAATAATGAATTCCCAG | pRU1097 backbone | this study |
| P183 | GGCTGTGGCCGATCTAGGGCTGCA | pRU197 backbone -NotI | this study |

| Primer | sequence (5'->3') | amplicon | reference |
|---|---|---|---|
| P184 | GCCGCAAACGTGGTCTGGTCGCGG | pRU197 backbone -NotI | this study |
| P195 | CAGGCGTAGCACCAGGCGTTTAAG | sequening primer pJABO | this study |
| P213 | ATATACCGATCGGAGCTCCACCGC | pRU1097 backbone -GFP | this study |
| P214 | CAGGCATCAAATAAAACGAAAGGC | pRU1097 backbone .GFP | this study |
| P217 | ACTGACGAGCTCCAGGCATCAAATAAAACGAAAGGC | *rrnB1* | this study |
| P220 | ACTGACCGATCGATAAAACGAAAGGCCCAGTCTTTCGACT | *rrnB1* | this study |
| P323 | ACTACGTCTAGACATATCATCCGCAGTGCTTG | 1st primer for PfAR-1 glr | this study |
| P324 | ACTATAGCGGCCGCTCAAGCAGAGCGTCGAGGAG | 1st primer for PfAR-1 glr | this study |
| P426 | TACTAAGCTGATCCGGTGGATGAC | sequencing primer pJABO | this study |
| P449 | TACAAGCATAAAGCTTGCTCAATCAATCACCGGATAGTGCCACCTGAACGAAGCATCTGT | yeast 2µ & *URA3* marker for pJABO5 | this study |
| P488 | AGGGTTAATTTCGAGCTTGGCGTAATCATGGTCATAGCTGTTTCCTGTGTGAAATTGTTA | lac promoter for pJABO5 | this study |
| P489 | TGTGAGCGGATAACAATTTCACACAGGAAACAGCTATGACCATGATTACGCCAAGCTCGA | lacZ fragment for pJABO5 | this study |
| P490 | GTTTTATTTGATGCCTGGGAATTCATTATTTGTAGTTACAATTTCCATTCGCCATTCAGG | lacZ fragment for pJABO5 | this study |
| P491 | CATGGGTGGAAGAGATGAAG | sequencing of pJABO5 | this study |
| P506 | AATGAGTGAGCTAACTCACATTAATTGCGTTGCGCCCTTACGCATCTGTGCGGTATTTCA | yeast 2µ & *URA3* marker for pJABO5 | this study |
| P507 | CTATGCGGTGTGAAATACCGCACAGATGCGTAAGGGCGCAACGCAATTAATGTGAGTTAG | lac promoter for pJABO5 | this study |
| P524 | TGTGAGCGGATAACAATTTCACACAGGAAACAGCTATGGCCTACGATTTTGATCTTTATG | $2^{nd}$ nested primer for PfAR-1 *glr* | this study |
| P525 | GTTTTATTTGATGCCTGGGAATTCATTATTTGTAGTTAAGCGCTGACCGGCGTACGCATG | $2^{nd}$ nested primer for PfAR-1 *glr* | this study |
| $P_{AmpF}$ | GTCAGAAGTAAGTTGGCCGCAGTG | part of $amp^R$ CDS | this study |
| $P_{AmpfR}$ | CATTTCCGTGTCGCCCTTATTCCC | part of $amp^R$ CDS | this study |
| $P_{GentF}$ | TCACCGTAATCTGCTTGCAC | part of $gent^R$ CDS | this study |
| $P_{GentR}$ | GGCTCAAGTATGGGCATCAT | part of $gent^R$ CDS | this study |
| $P_{M13(-20)35S-Enh2}$ | GTAAAACGACGGCCAGT | sequencing primer | this study |
| $P_{reverse}$ | GGAAACAGCTATGACCATG | sequencing primer | this study |
| Tn*pho*A II | GTGCAGTAATATCGCCCTGAGCA | IS-Ω-Km/hah h | (Manoil, 2000) |

### [SM5] Chemical Synthesis of Allicin

Chemical synthesis of and analysis of allicin was performed as described in Gruhlke et al. (2010) with minor adjustments:

1. A beaker was filled with ice-water and a magnetic stirrer bar and placed on a magnetic stirrer. Then, a magnetic stirrer bar was placed in a 100 ml round-bottomed flask which was fixed within the ice-water bath.
2. While stirring, 2 ml 99 % re-distilled diallyl disulfide (DADS) and 5 ml 100 % acetic acid were added. Then, 3 ml 30 % $H_2O_2$ were slowly added to this mixture.
3. The round-bottomed flask and the beaker were wrapped in aluminum foil and the reaction was incubated with stirring for 30 minutes at 0 °C.
4. The beaker with the ice-water was removed and the reaction allowed to continue with stirring for 2 h at room temperature.
5. After 2 h, the reaction was stopped by the addition of 25 ml $H_2O$

6. Then, allicin and byproducts were extracted twice with 30 ml dichloromethane (DCM) using a separating funnel. The organic phase was retained, and the aqueous phase was discarded.
7. Next, the organic phase was neutralized by the addition of 5 % $NaHCO_3$ until no more gas-development was observed.
8. The aqueous phase was discarded, and the organic phase was washed several times with water to remove remaining sodium acetate completely.
9. The organic phase was transferred to a round bottomed flask, which was then placed in an ice water bath and solvent was removed by rotary evaporation at reduced pressure .
10. Approximately 100 ml $H_2O$ were added to the oily allicin residue, and the solution checked for purity by HPLC analysis.
11. The aqueous phase was washed several times with n-Hexane to remove unreacted DADS, and the solution checked for purity by HPLC analysis. The organic phase was discarded and the allicin in the aqueous phase was extracted with DCM (step 6)
12. DCM was removed by rotary evaporation at reduced pressure (as in step 9) and 40 ml $H_2O$ were added to the remaining oil (allicin).

The diluted allicin was quantified via HPLC analysis and then aliquoted and frozen at -70 °C until use.

### [SM6] High Pressure Liquid Chromatography Analysis (HPLC) of allicin

Synthesized allicin was analyzed by diluting the allicin tenfold in 100 % methanol. 20 µl of this dilution were loaded on the HPLC. Water (A) and Methanol (C) were used as mobile phase for the separation on a reverse phase silica column (C18) at 25 °C with a flow rate of 1 ml/min. The following gradients were used for separation (Table S5). The HPLC was calibrated with an NMR-confirmed allicin standard.

Table S5:Solvent gradient chart for the HPLC analysis of allicin.

| **Time (minute)** | **Solvent A (water)** | **Solvent C (100% methanol)** |
|---|---|---|
| prerun | 56 % | 44 % |
| 10 | 53.2 % | 46.8 % |
| 15 | 7 % | 93 % |
| 30 | 7 % | 93 % |
| 31 | 56 % | 44 % |
| 35 | 56 % | 44 % |

<u>The HPLC equipment setup was as followed:</u>

| | |
|---|---|
| pump block PU-980 | Jasco GmbH Deutschland |
| UV-detector | Jasco GmbH Deutschland |
| Degasser DG-980-50 | Jasco GmbH Deutschland |
| gradient unit LB-980-02 | Jasco GmbH Deutschland |
| precolumn (Multohigh 100, RP18 5 µ) | CS-Chromatographie Service GmbH |
| ProntoSIL C18 | Bischoff Chromatographie |
| column thermostat (CO-2060 plus) | Jasco GmbH Deutschland |

#### [SM7] Protocol for high yield genomic DNA extraction from bacteria

For preparing a genomic DNA library of *Pf*AR-1, a new protocol for high yield DNA extraction was established based on Chen and Kuo (1993) and on Syn and Swarup (2000). A 50 ml bacterial culture was grown overnight in liquid medium in a 500 ml Erlenmeyer flask. Bacteria were harvested by centrifugation (2,500 x *g* for 20 minutes at 4 °C) in a 50 ml reaction tube. The cell pellet was suspended in 20 ml of 1 % NaCl solution (w/v in double distilled water ($H_2O_{dd}$)) for the removal of bacterial EPS. Therefore, the cells were vortexed vigorously in the NaCl solution and harvested again by centrifugation. For removal of NaCl, the bacterial cells were washed twice with 50 ml $H_2O_{dd}$ by vigorous vortexing and harvesting by centrifugation. The cells were finally suspended in 40 ml $H_2O_{dd}$. The cell solution was distributed among 2 ml reaction tubes and harvested at 12,879 x *g* for three minutes at 4 °C. Afterwards, the supernatant was removed to the last drop. The cell pellets were vortexed without addition of buffer to loosen the cells from each other, thereby increasing the available surface for the subsequent lysis step. Bacterial lysis was done by addition of 1.36 ml lysis buffer (40 mM TRIS-HCl pH 7.8, 20 mM sodium-acetate, 1 mM EDTA, 1 % SDS (w/v, = 35 mM)) to each reaction tube and mixing by pipetting up and down. The tubes were then incubated for 60 minutes in a 50 °C water bath for enhanced lysis and DNA yield. Then, 12 µl of RNase I (10 mg/ml) were added to each reaction tube and incubated for 30 minutes at 37 °C. To precipitate cell debris and SDS, 476 µl 5 M NaCl were added to each reaction tube and mixed gently. The cell debris and SDS were then separated from the remaining solution via centrifugation at 20,937 x *g* for 20 minutes at 4 °C.

For further purification, 1.6 ml from the supernatant of each reaction tube was gathered in an autoclaved glass bottle. Afterwards, the bottle was filled up with dilution buffer (40 mM TRIS-HCl pH 7.8, 20 mM sodium-acetate, 1 mM EDTA, 150 mM NaCl) to approximately 200 ml for dilution. The bottle was placed on ice.

For phase extraction, 5 ml of chloroform were added to 40 ml centrifugation tubes, respectively. The tubes were then filled up with the DNA solution which were gathered previously in the glass bottle and inverted 50 times. The phases were separated by centrifugation at 21,000 x *g* for 3 minutes at 4 °C. The supernatant was gathered in a new sterile glass bottle. These extraction steps were repeated for the whole DNA solution in the glass bottle until no interphase was visible any more.

For DNA precipitation, 25 ml of phase-extracted DNA solution was added to 50 ml reaction tubes and mixed with 25 ml isopropanol. Since the lysis buffer and the dilution buffer contained enough salt (not removed during former steps), no further salt addition was needed for precipitation. The DNA-isopropanol/solutions were stored at -20 °C until all the remaining solution was processed to this stage of this protocol.

The DNA was subsequently precipitated into the same tubes at 21,000 x *g* for 15 minutes at 4 °C. The two DNA pellets were washed twice with 70 % ethanol. Last droplets of ethanol were removed via a Pasteur pipette. DNA pellets were dissolved in 10 mM of TRIS-HCl pH 8. The DNA was then aliquoted and stored at -20 °C. A sample of extracted DNA with this protocol is shown in Fig. S1.

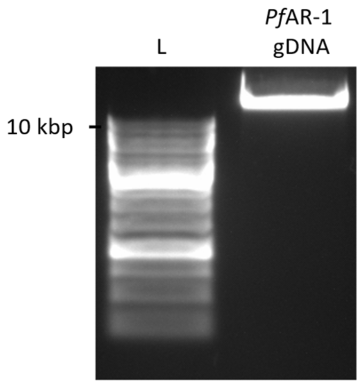

Figure S1: Sample of genomic DNA extracted with a newly established protocol that was based on the combination of the protocols published by Chen & Kuo and by Choong (Chen and Kuo, 1993; Syn and Swarup, 2000). DNA was extracted from 50 ml over-night culture from *Pf*AR-1 grown at 28 °C and 220 rpm in King´s B medium. After DNA extraction, 500 µl pure DNA solution were obtained. From that, 2 µl of DNA was checked via gel electrophoresis. L: GeneRuler 1 kb DNA ladder (Thermo Scientific).

### [SM8] *Pf*AR-1 genomic library construction

Genomic DNA was extracted as described in SM7 and partially digested with Sau3AI FD (Thermo Scientific). Sau3AI FD was diluted 300 fold in 1x FastDigest buffer (Thermo Scientific) and was applied to the reaction mixture for a 3000 fold enzyme dilution. Digested DNA was size-separated via agarose gel electrophoresis and fragments of approx. 10 kbp were extracted and purified using Zymoclean Large Fragment DNA Recovery Kit, subcloned in BamHI digested pRU1097, and electroporated in *E. coli* K12 DH10B MegaX. Plasmid DNA of approx. 14,000 *E. coli* transformant colonies was extracted, representing more than 99.99 % theoretical coverage of the *Pf*AR-1 genome.

### [SM9] Transposon Mutagenesis of genomic clone 1

Transposon mutagenesis of *Pf*AR-1 genomic clones on pRU1097 was done in the *Ps*4612 background. Briefly, pSCR001 carrying transposon IS-Ω-km/hah (Giddens et al., 2007) was transferred from *E. coli* S17 via biparental mating to *Ps*4612 and transconjugants were selected on gentamycin and kanamycin. Since pSCR001 cannot replicate in *Ps*4612, plasmid isolation from the transconjugants yields a Tn-carrying pRU1097 population, that was transformed in *E. coli* MegaX DH10B (Fig. S2). Plasmid DNA of more than 10,000 *Ps*4612 genomic clone 1 transconjugants was extracted and elextroporated electroporated in *E. coli* K12 DH10B MegaX to construct a library of Tn-carrying genomic clone 1.

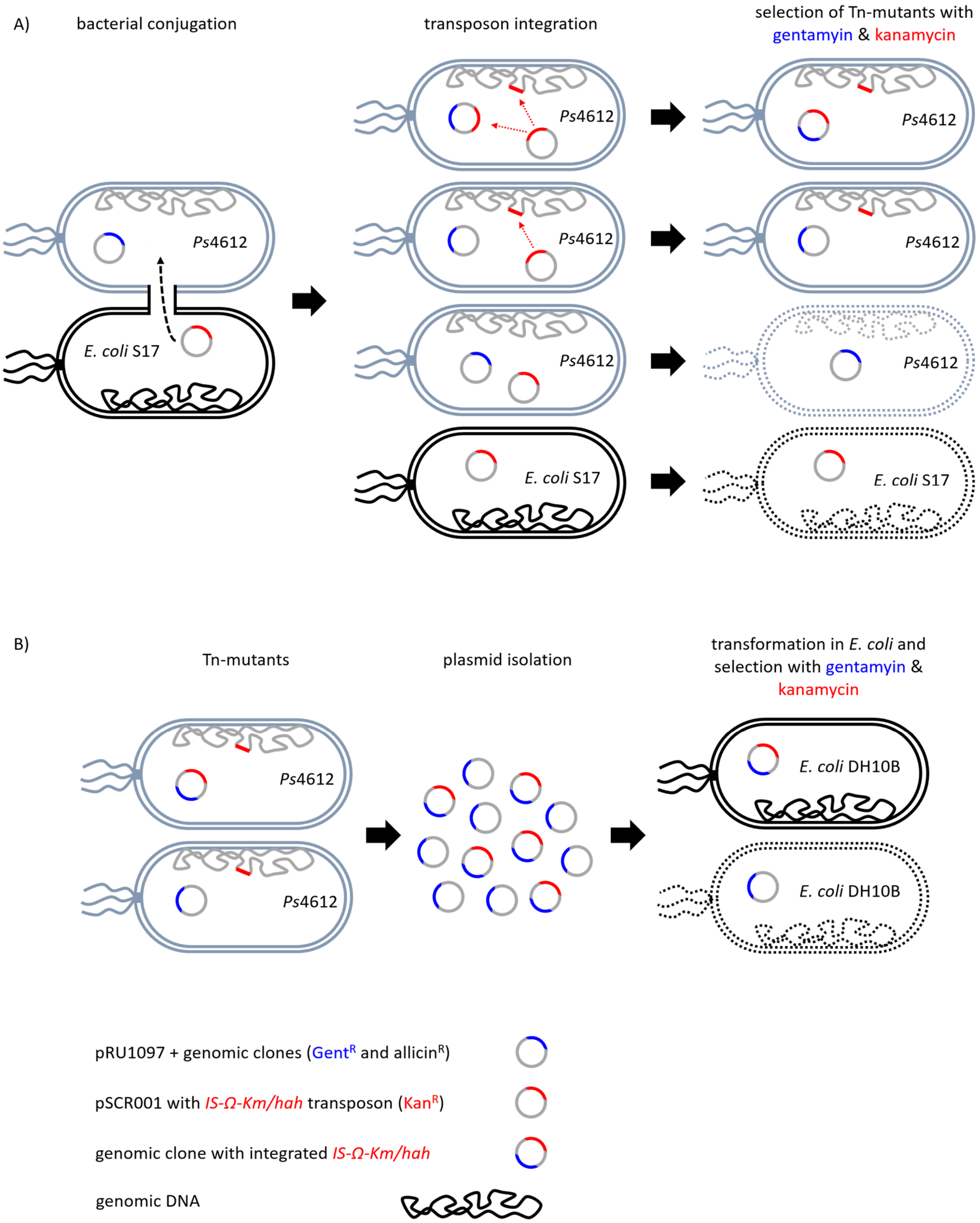

**Figure S2: Strategy for transposon mutagenesis. (A)** mutagenesis and selection in *Ps*4612 **(B)** Plasmid DNA isolated from *Ps*4612 Tn-mutants and transformed into *E. coli* to select for pRU1097::Tn on GentR and KanR.

#### [SM10] Construction of the broad host range expression vector pJABO

The construction of the expression vector pJABO is shown in Figs. S3-S8. Linearized pRU1097 was amplified via PCR with primers P163 and P174; thus adding ApaLI and XhoI restriction sites at the ends. The promoter from the neomycin phosphotransferase gene (Neo prom.) was amplified with the primers P160 and P159; thus adding the restriction sites NheI, PvuI, and XhoI upstream and ApaLI downstream of the promoter, respectively. Both the above PCR products were digested with ApaLI and XhoI and ligated together to give the pRU1097+Neo promoter intermediate (Fig. S3).

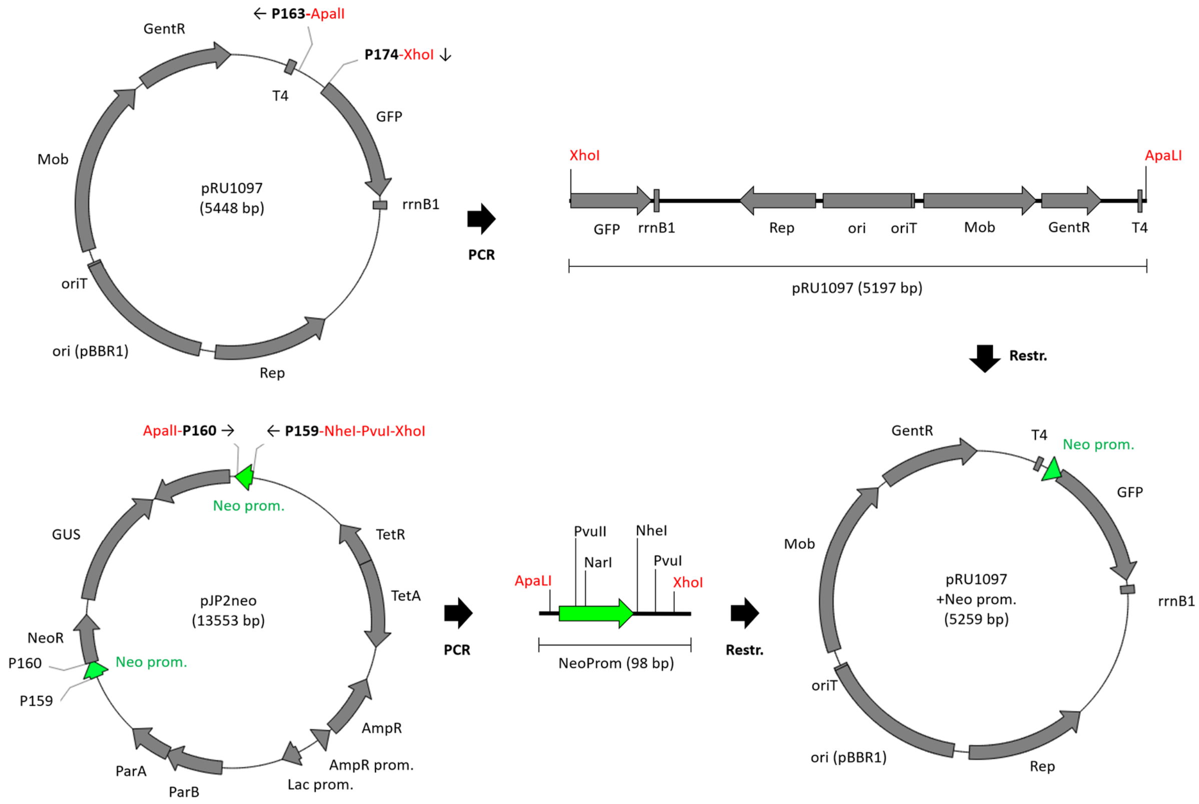

Figure S3: Subcloning of the neomycin phosphotransferase promoter in pRU1097.

Next, the multiple cloning site (MCS) from pBluescript I KS (-) was amplified with the primers P161 and P162; thus adding the restriction sites NheI and PvuI After restriction with PvuI and NheI, this was ligated with pRU1097+Neo promoter to give pRU1097+Neo+MCS (Fig. S4).

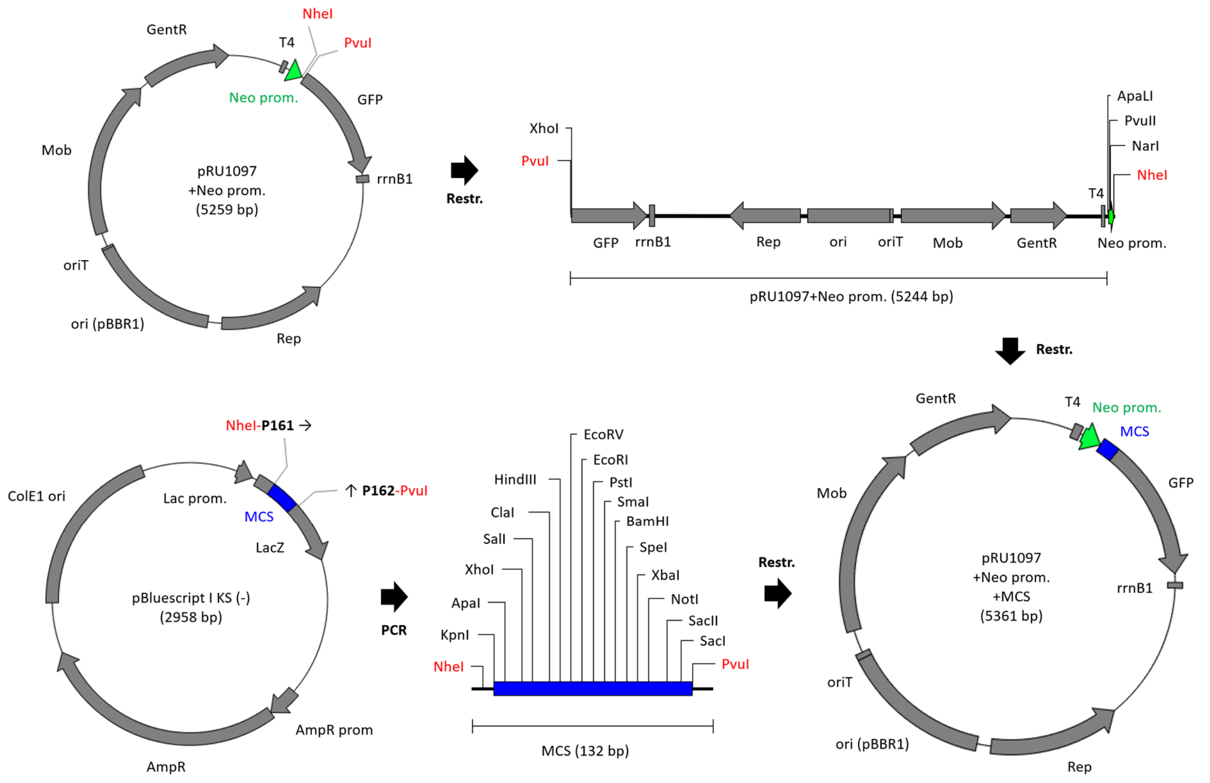

Figure S4: Subcloning of the multiple cloning site of pBluescript in pRU1097+Neo.

The NotI restriction site in the mobilization gene (Mob) from pRU1097+Neo+MCS was removed by whole vector amplification using the primers P183 and P184 and subsequent blunt end ligation. Primer P183 introduces a nucleotide exchange within the recognition sequence for NotI, resulting in the deletion of NotI without changing the encoded amino acid (Fig. S5).

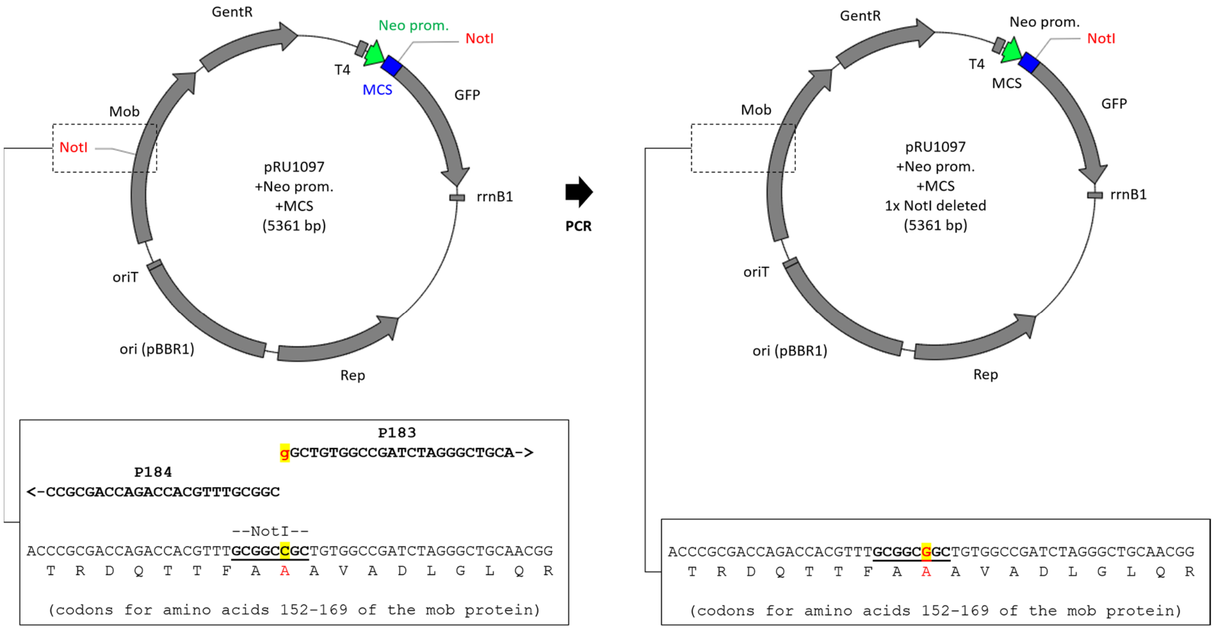

Figure S5: Deletion of NotI restriction site in the mobilization protein sequence of pRU1097.

The elimination of the restriction site was checked via restriction analysis (data not shown) and the constructed vector was analysed by DNA sequencing. Sequencing showed that all components for gene expression except for the rrnB1 terminator sequence were present (Fig. S6).

```
                                                    --------------T4-
Ref.  tgaccttttgaatgacctttaatagattatattactaattaattggggaccctagaggtc
P195  tgaccttttgaatgacctttaatagattatattactaattaattggggaccctagaggtc
P426                                              tggggancctagaggtc

      --------------                                       -------
Ref.  cccttttttattttaaaatttttttcacaaaacggtttacaagcatagtgcacccggaat
P195  cccttttttattttaaaatttttttcacaaaacggtttacaagcatagtgcacccggaat
P426  cccttttttattttaaaatttttttcacaaaacggtttacaagcatagtgcacccggaat

      --------------------NeoProm----------------      -----------
Ref.  tgccagctggggcgccctctggtaaggttgggaagccctgcaagctagcggtaccgggcc
P195  tgccagctggggcgccctctggtaaggttgggaagccctgcaagctagcggtaccgggcc
P426  tgccagctggggcgccctctggtaaggttgggaagccctgcaagctagcggtaccgggcc

      ---------------------------multiple cloning site------------
Ref.  cccccctcgaggtcgacggtatcgataagcttgatatcgaattcctgcagcccgggggatc
P195  cccccctcgaggtcgacggtatcgataagcttgatatcgaattcctgcagcccgggggatc
P426  cccccctcgaggtcgacggtatcgataagcttgatatcgaattcctgcagcccgggggatc

      --------------------------------SacI---PvuI-               -
Ref.  cactagttctagagcggccgccaccgcggtggagctccgatcggtatatcaggcatcaaa
P195  cactagttctagagcggccgccaccgcggtggagctccgatcggtatatc------cca-
P426  cactagttctagagcggccgccaccgcggtggagctccgatcggtatatc---c---ca-

      ------------------rrnB1-Ter----------------
Ref.  taaaacgaaaggctcagtcgaaagactgggcctttcgttttatctgttgtttgtcggtga
P195  ------g------tctttc----gactgagcctttcgttttatttgatgtttgtcggtga
P426  -----------g-tctttc----gactgagcctttcgttttatttgatgtttgtcggtga
```

Figure S6: Sequencing data for the gene regulatory sequences of modified pRU1097. pRU1097 was modified for gene expression. Gene regulatory sequences and features for subcloning and gene expression (*T4 terminator*, *neomycin phosphotransferase promoter* [NeoProm, green], the multiple cloning site of pBluescript [blue] and the *rrnB1 Terminator* [red]) were sequenced using the primers P195 and P426. The sequences were compared against the expected reference sequence from the modified vector. Highlighted SacI and PvuI restriction sites could be used to insert a function *rrnB1 terminator* by a restriction/ligation strategy.

To restore the *rrnB1* terminator somehow lost during the previous steps, the sequence was reamplified from pRU1097 with the primers P217 and P220; adding SacI and PvuI restriction sites for subcloning (Fig. S7).

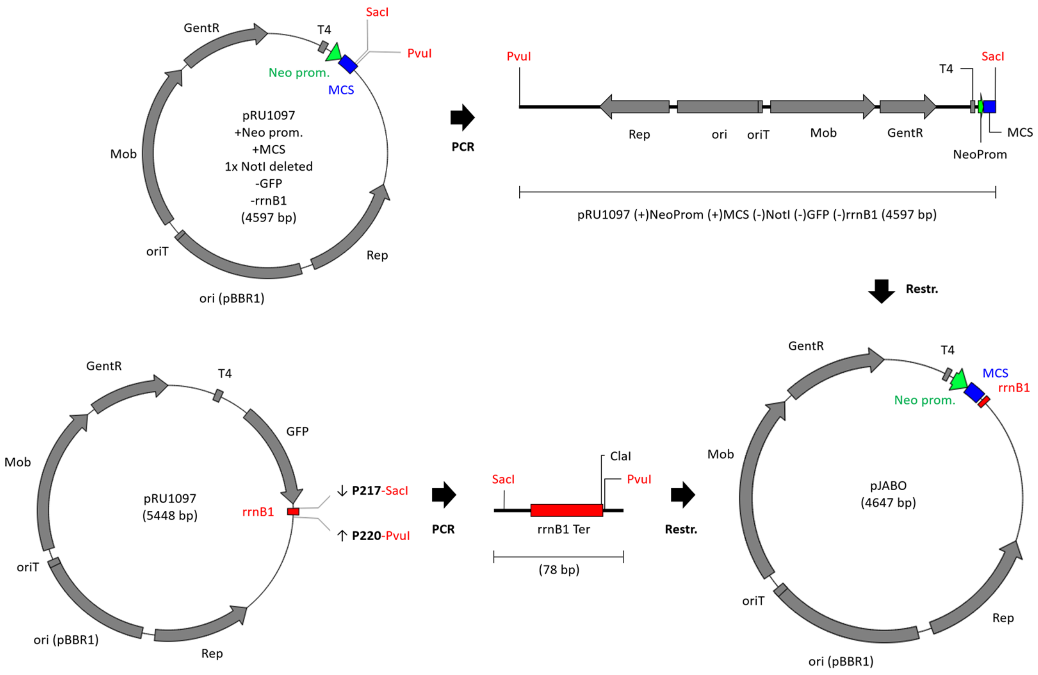

Figure S7: Subcloning of the *rrnB1 terminator* to construct the broad host range expression vector pJABO.

The final vector construct pJABO was verified by restriction analysis and DNA sequencing of the promoter and the multiple cloning site as well as their flanking terminator sequences *T4* and *rrnB1* (Fig. S8).

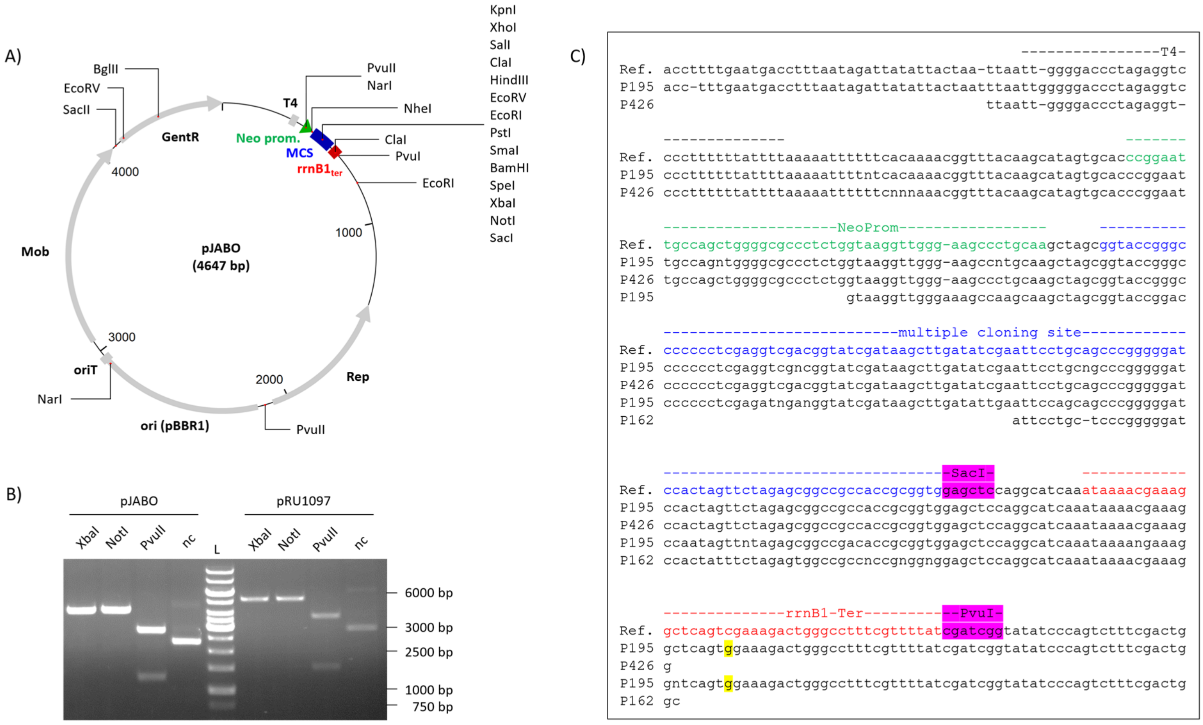

Figure S8: Characteristics of the broad host range expression vector pJABO. (A) Vector map of pJABO. (B) Restriction digest in comparison to the original vector pRU1097. (C) Sequencing of the necessary elements needed for heterologous gene expression. Primers P195, P162 and P426 were used for sequencing and the sequencing data were compared against the expected reference sequence. The restriction sites that were used for *rrnB1 terminator* subcloning are highlighted in purple (SacI, PvuI). Discrepancies to the expected reference sequence are highlighted in yellow. L: GeneRuler 1 kb DNA ladder.

### [SM11] Construction of the broad host range vector pJABO5 and cloning of *Pf*AR-1 glutathione reductase *glr1* gene for inducible expression in *E. coli*

pJABO5, which was used for the expression of the *Pf*AR-1 glutathione reductase in *E. coli*, was constructed by *in vivo*-recombination in yeast. In comparison to pJABO, which was used for overexpression, pJABO5 was designed for induced gene expression based on the inducible lac promoter from *E. coli*.

pRU1097 was digested over night with XbaI and SacI, thereby removing *GFP* from pRU1097. Next, *yeast 2µ ori* and the *URA3* selection marker were amplified from pRS426 via PCR using the primers P449 and P506. The *lac promoter* was amplified from *E. coli* MG1655 genomic DNA with primers P488 and P507, and the *lacZ* fragment was amplified from pBluescript I KS (-) using the primers P489 and P490. The vector backbone fragment of pRU1097 and the PCR products were transformed in *Saccharomyces cerevisiae* BY4742 (Jansen et al., 2005). The vector was extracted from yeast by alkaline lysis and re-transformed into *E. coli* for amplification. The pJABO5 map is shown in Fig. S9.

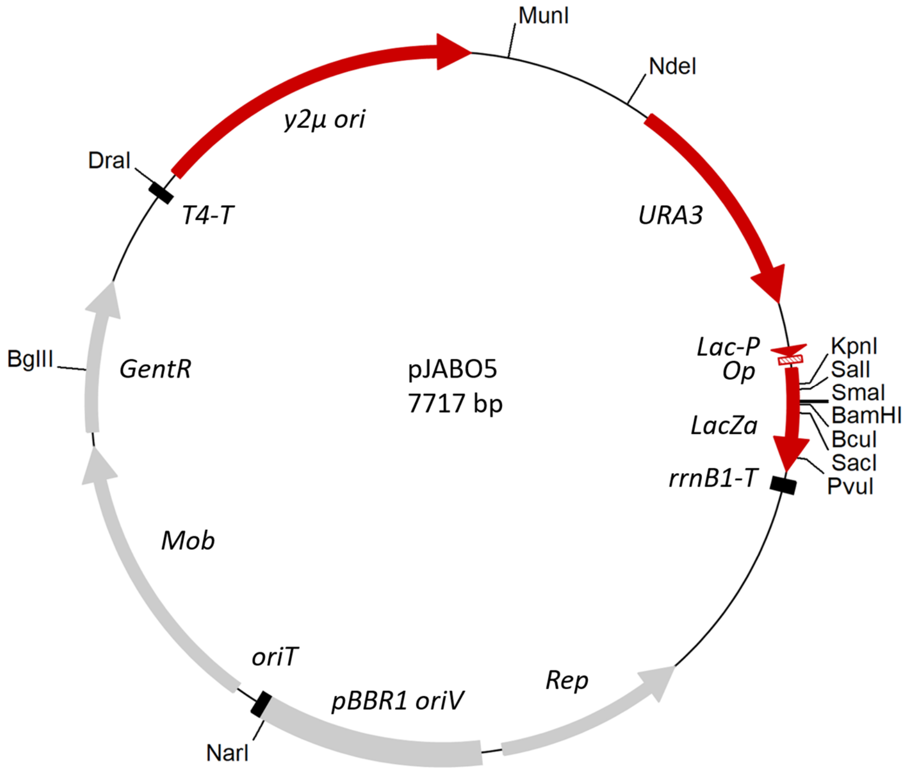

Figure. S9: Vector Map of pJABO5.

For cloning of *Pf*AR-1 *glutathione reductase 1* (*glr1*, fig|294.271.peg.2394), *glr1* had to be amplified via a nested PCR since the different glutathione reductases within the core genome and the horizontally transferred regions were too similar for separate, one-step amplification. Thus, the first PCR amplicon from *Pf*AR-1 genomic DNA was generated with the primers P323 and P324, and used as a template for the amplification of *Pf*AR-1 *glr1* with the primers P524 and P525. The final product was cloned in pJABO5 by *in vivo* recombination in yeast (Jansen et al., 2005). pJABO5 was digested with BamHI and LacZα was replaced by *Pf*AR-1 *glr1*. The recombinant vector was isolated from yeast and directly transformed in *E. coli* BW25113 wild type, or *E. coli* BW25113 Δ*glr*. The presence of the subcloned *glr1* was verified by PCR using the primers P195 and P491.

#### [SM12] Glutathione disulfide reductase enzyme assay

For glutathione reductase activity assays, cells were grown overnight at 28 °C in liquid M9JB medium. Cells from 20 ml over night culture were harvested by centrifugation (3,000 x g at room temperature) and cells were re-suspended in 1 mL phosphate buffer (143 mM Na-Phosphate containing 6.2 mM EDTA, pH 7.5). Bacteria were lysed mechanically by vortexing with 1 mm glass beads three times for 1 minute on ice. Cell debris was removed by centrifugation at 21,000 x *g* for 1 minute at room temperature.

Glutathione reductase activity was measured in a glutathione reductase recycling assay Horn et al., 2018 modified to conditions showing linear dependency of the reaction velocity for enzyme amount, i.e. not substrate-limited. Absorption was followed over 10 minutes at 412 nm using a spectrophotometer (DU800, Beckman Coulter GmbH, Krefeld, Germany). Enzyme activity was calculated assuming a molar extinction coefficient of TNB of 13,600 $M^{-1}cm^{-1}$. (Miron et al., 1998). For calculation of specific enzyme activity, protein content of the sample was measured using the Bradford method (Bradford, 1976).

#### [SM13] List of independent *Pf*AR-1 genomic clone 1 Tn-mutants

To screen for loss-of-function after Tn-mutagenesis, colonies were picked and tested in a streak assay. Positive (*E. coli* MegaX DH10B with non-mutagenized genomic clone) and negative controls (*E. coli* MegaX DH10B with empty vector pRU1097) strains were included on each streak assay plate together with six different transposon mutants.

The data from each plate were not pooled but were evaluated individually, because the controls showed slight variation between plates. The distance between the hole and the first growth of the transposon mutants was inversely proportional to allicin resistance. For each plate, the relative allicin resistance of each mutagenized clone was calculated in relation to clone 1 on that plate. For example, for mutant IsΩ 042, the relative allicin resistance for clone 1 is higher than for IsΩ 042, thereby indicating a loss of allicin resistance due to a transposon insertion:

$$rel.resistance\ clone\ 1 = \frac{clone\ 1}{clone\ 1} = \frac{1.00\ cm}{1.00\ cm} = 1.00$$

$$rel.resistance\ \mathrm{Is}\Omega\ 042 = \frac{clone\ 1}{\mathrm{Is}\Omega\ 042} = \frac{1.00\ cm}{1.70\ cm} = 0.59$$

By setting each dataset in relation to the positive control, all data could be pooled for comparison to get an overview of the number of more resistant, equally resistant and less resistant mutants compared to the non-mutagenized clone 1 (Fig. S10).

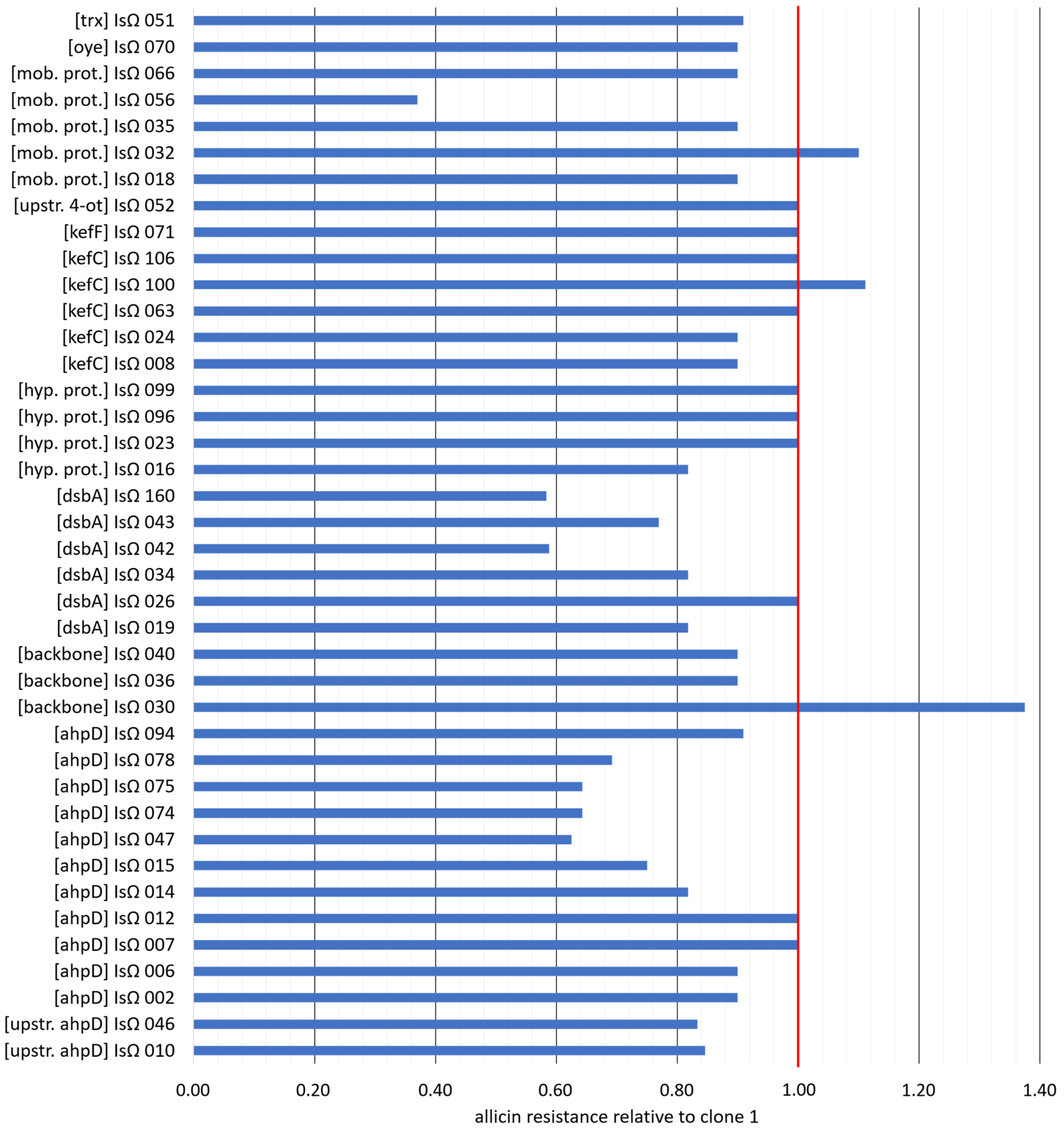

Figure S10: Relative allicin resistance of *E. coli* transformed with transposon mutagenized *Pf*AR-1 genomic clone 1 plasmid DNA. *E. coli* with genomic clone 1 and putative transposon mutants of genomic clone 1 (IsΩ mutants) were investigated via a streak assay on LB media against allicin. The media (20 ml solid LB medium in petri dishes of 9 cm in diameter) contained a hole with 1 cm in diameter. Single colonies were harvested in 100 µl liquid LB medium and mixed. After the different bacterial strains were streaked from the hole to the edge of the plate with an inoculation loop, the hole was filled with 120 µl 10 mM allicin and the plates were incubated overnight at 37 °C. The red vertical line highlights the relative allicin resistance of non-mutagenized clone 1.

### [SM14] Annotation of *Pf*AR-1 genomic repeats (RE)

Table S6: Genes from the three genomic repeats RE1, RE2, and RE3 located in the *Pf*AR-1 genome. Original annotation from RAST was supplemented by individual annotation of the genomic repeats without the *Pf*AR-1 genome and by manual curation (Open Reading Frame Finder (NCBI), BLASTp). Congruent gene set is marked in green, features in addition to the original RAST annotation are marked in orange.

| gene identifier | | | % identity on amino acid level | | | gene annotation | RE core annotation |
|---|---|---|---|---|---|---|---|
| RE1 | RE2 | RE3 | RE1 vs RE2 | RE1 vs RE3 | RE2 vs RE3 | | |
| fig\|294.271. peg.1057 | fig\|294.271. peg.2576 | fig\|294.271. peg.3655 | 97.91 | 67.03 | 68.11 | Transcriptional regulator%2C TetR family | *tetR* |
| | | fig\|RE3_G | | | | hypothetical protein | |
| fig\|RE1_A | | | | | | hypothetical protein | |
| fig\|RE1_B | fig\|RE2_A | | 98.10 | | | Permease of the drug/metabolite transporter (DMT) superfamily | |
| fig\|RE1_C | | | | | | hypothetical protein | |
| fig\|RE1_D | | | | | | Outer membrane protein W precursor | |
| fig\|RE1_F | | | | | | probable short-chain dehydrogenase | |
| fig\|RE1_G | | | | | | Hydrolase, alpha/beta fold family protein | |
| fig\|RE1_H | | | | | | Transcriptional regulator, TetR family | |
| fig\|294.271. peg.1058 | | | | | | hypothetical protein | |
| fig\|294.271. peg.1059 | fig\|294.271. peg.2577 | | 93.81 | | | Cystathionine beta-lyase (EC 4.4.1.8);Ontology_term=KEGG_ENZYME:4.4.1.8 / Other: Cystathionine gamma-lyase (EC 4.4.1.1);Ontology_term=KEGG_ENZYME:4.4.1.1 | |
| fig\|RE1_I | | | | | | hypothetical protein | |
| | fig\|RE2_B | | | | | hypothetical protein | |
| fig\|294.271. peg.1060 | fig\|294.271. peg.2578 | | 96.36 | | | L-asparagine permease / Other: D-serine/D-alanine/glycine transporter | |
| | fig\|RE2_C | | | | | hypothetical protein | |
| fig\|294.271. peg.1061 | fig\|294.271. peg.2579 | fig\|294.271. peg.3654 | 98.91 | 89.67 | 89.67 | OsmC family protein | *osmC* |
| fig\|294.271. peg.1062 | fig\|294.271. peg.2580 | fig\|294.271. peg.3653 | 97.67 | 91.05 | 91.83 | Short-chain dehydrogenase/reductase SDR / Other: 3-oxoacyl-[acyl-carrier protein] reductase (EC 1.1.1.100);Ontology_term=KEGG_ENZYME:1.1.1.100 | *sdr* |
| fig\|294.271. peg.1063 | fig\|294.271. peg.2581 | fig\|294.271. peg.3652 | 98.98 | 86.73 | 87.76 | Transcriptional regulator%2C TetR family | *tetR* |
| fig\|294.271. peg.1064 | fig\|294.271. peg.2582 | fig\|294.271. peg.3651 | 96.98 | 77.16 | 75.43 | isomerase%2C putative / Other: 2-hydroxychromene-2-carboxylate isomerase/DsbA-like thioredoxin domain | *dsbA* |
| fig\|294.271. peg.1065 | fig\|294.271. peg.2583 | fig\|294.271. peg.3650 | 95.52 | 86.96 | 88.06 | Thiosulfate sulfurtransferase%2C rhodanese (EC 2.8.1.1);Ontology_term=KEGG_ENZYME:2.8.1.1 | *trx* |
| fig\|294.271. peg.1066 | fig\|294.271. peg.2584 | fig\|294.271. peg.3649 | 98.52 | 92.98 | 92.98 | Glutathione-regulated potassium-efflux system protein KefC / Other: Glutathione-regulated potassium-efflux system protein KefB | *kefC* |
| fig\|RE1_J | fig\|RE2_D | fig\|294.271. peg.3648 | 93.64 | 77.91 | 79.07 | Glutathione-regulated potassium-efflux system ancillary protein KefF | *kefF* |
| fig\|294.271. peg.1067 | fig\|294.271. peg.2585 | fig\|294.271. peg.3647 | 98.61 | 97.22 | 95.83 | 4-oxalocrotonate tautomerase family protein | *4-ot* |
| fig\|294.271. peg.1068 | fig\|294.271. peg.2586 | fig\|294.271. peg.3646 | 99.18 | 94.29 | 94.02 | NADH:flavin oxidoreductases%2C Old Yellow Enzyme family / Other: N-ethylmaleimide reductase (EC 1.-.-.-);Ontology_term=KEGG_ENZYME:1.-.-.- | *oye* |
| fig\|RE1_K | | fig\|RE3_F | | 77.13 | | Acyl carrier protein phosphodiesterase (EC 3.1.4.14) | |
| RE1_14 | | | | | | hypothetical protein | |
| fig\|294.271. peg.1069 | fig\|294.271. peg.2587 | fig\|294.271. peg.3645 | 100.00 | 98.78 | 98.82 | Possible carboxymuconolactone decarboxylase family protein (EC 4.1.1.44);Ontology_term=KEGG_ENZYME:4.1.1.44 | *ahpD* |
| fig\|294.271. peg.1070 | | | | | | hypothetical protein | |
| fig\|294.271. peg.1071 | fig\|294.271. peg.2588 | fig\|RE3_A | 91.18 | 87.50 | 89.22 | EthD reductase (annotated by RAST as hypothetical protein) | *ethD reductase* |
| | | fig\|294.271. peg.3644 | | | | Transcriptional regulator%2C AsnC family | |
| | | fig\|RE3_E | | | | Cyclohexadienyl dehydratase (EC 4.2.1.51)(EC 4.2.1.91) # Periplasmic precursor | |
| | | fig\|294.271. peg.3643 | | | | amino acid ABC transporter%2C ATP-binding protein / Histidine ABC transporter%2C ATP-binding protein HisP (TC 3.A.1.3.1) | |
| | | fig\|RE3_D | | | | polar amino acid ABC transporter, inner membrane subunit | |
| | | fig\|RE3_C | | | | polar amino acid ABC transporter, inner membrane subunit | |
| | | fig\|294.271. peg.3642 | | | | FAD dependent oxidoreductase | |
| | | fig\|294.271. peg.3641 | | | | Oxidoreductase (EC 1.1.1.-);Ontology_term=KEGG_ENZYME:1.1.1.- | |
| | | fig\|RE3_B | | | | Endoribonuclease L-PSP | |
| | | fig\|294.271. peg.3640 | | | | Cysteine synthase B (EC 2.5.1.47);Ontology_term=KEGG_ENZYME:2.5.1.47 | |
| | | fig\|RE3_A | | | | hypothetical protein | |
| fig\|294.271. peg.1072 | fig\|294.271. peg.2589 | fig\|294.271. peg.3639 | 98.63 | 91.10 | 90.75 | Alpha-ketoglutarate-dependent taurine dioxygenase (EC 1.14.11.17);Ontology_term=KEGG_ENZYME:1.14.11.17 | *tauD dioxygenase* |
| fig\|294.271. peg.1073 | fig\|294.271. peg.2590 | fig\|294.271. peg.3638 | 98.50 | 93.23 | 93.05 | Carotenoid cis-trans isomerase (EC 5.2.-.-);Ontology_term=KEGG_ENZYME:5.2.-.- | *oxidoreductase* |
| fig\|294.271. peg.1074 | fig\|294.271. peg.2591 | fig\|294.271. peg.3637 | 96.97 | 80.99 | 80.99 | Transcriptional regulator%2C AraC family | *araC* |
| | fig\|294.271. peg.2592 | | | | | Transcriptional regulator%2C LysR family | |
| | fig\|294.271. peg.2593 | | | | | Aldo/keto reductase / Oxidoreductase%2C aldo/keto reductase family | |
| fig\|294.271. peg.1075 | fig\|294.271. peg.2594 | | 98.89 | | | Glutathione reductase (EC 1.8.1.7);Ontology_term=KEGG_ENZYME:1.8.1.7 | |
| fig\|RE1_L | fig\|RE2_E | | 98.18 | | | hypothetical protein | |
| fig\|294.271. peg.1076 | fig\|294.271. peg.2595 | | 100.00 | | | Bll2902 protein | |

### [SM15] Comparison of putative RE regions across the *Pseudomonas* genus.

Table S7. Syntenic regions in pseudomonads other tha *Pf*AR-1. The information in this Table is from the Pseudomonas.com database.

| | **synt. regions** | **Location Name** | **isolation source** | **Assembly Accession** | **additional information** |
|---|---|---|---|---|---|
| **complete genomes** | | | | | |
| *Pseudomonas brassicacearum* DF41 | 1 | Manitoba | canola root tip | GCF_000585995.1 | |
| *Pseudomonas brassicacearum* LBUM300 | 1 | Canada: New Brunswick: Bouctouche | soil, Canada | GCF_001449085.1 | |
| *Pseudomonas fluorescens* A506 | 1 | USA: California | pear tree leaf | GCF_000262325.2 | |
| *Pseudomonas fluorescens* FW300-N2E3 | 1 | USA: Oak Ridge, TN | ground water from background well at DOE's FRC site at Oak Ridge | GCF_001307155.1 | |
| *Pseudomonas fluorescens* Pt14 | 1 | India: Tinsukia, Assam | Rhizosphere soil | GCF_001747385.1 | |
| *Pseudomonas frederiksbergensis* ERGS4:02 | 1 | | glacial stream | GCF_001874645.1 | |
| *Pseudomonas syringae* pv. tomato DC3000 | 1 | | | GCF_000007805.1 | host: *Solanum lycopersicum* (tomato) |
| *Pseudomonas trivialis* IHBB745 | 1 | India: Rong Tong, Lahual and Spiti | rhizosphere | GCF_001186335.1 | host: *Hippophae rhamnoides* (sea buckthorn) |
| **draft genomes** | | | | | |
| *Pseudomonas aeruginosa* ATCC 33988 | 1 | USA: Ponca City, OK | fuel tank | GCF_000756575.1 | |
| *Pseudomonas aeruginosa* ATCC 9027 | 1 | Australia: Sydney | | GCF_001294675.1 | host: outer ear infection |
| *Pseudomonas aeruginosa* AZPAE12138 | 1 | USA: New York | | GCF_000796525.1 | host: *homo sapiens* (human, cystic fibrosis) |
| *Pseudomonas aeruginosa* AZPAE14813 | 1 | India: Mumbai | | GCF_000795085.1 | host: *homo sapiens* (human, urinary tract infection) |
| *Pseudomonas aeruginosa* AZPAE14898 | 1 | India: Chennai | | GCF_000791035.1 | host: *homo sapiens* (human, respiratory tract infection ) |
| *Pseudomonas aeruginosa* AZPAE14947 | 1 | China: Beijing | | GCF_000794045.1 | host: *homo sapiens* (human, urinary tract infection ) |
| *Pseudomonas aeruginosa* TRN6649 | 1 | | | GCF_001921175.1 | host: *homo sapiens* (human) |
| *Pseudomonas aeruginosa* WH-SGI-V-07643 | 1 | USA | Hospital | GCF_001452255.1 | |
| *Pseudomonas amygdali* pv. *tabaci* ATCC 11528 | 1 | | | GCF_000145945.1 | This strain was sequenced independently in 3 different labs, with slight variations<br>host: *Nicotiana tabacum* (common tobacco) |
| *Pseudomonas amygdali* pv. *tabaci* ATCC 11528 | 1 | | | GCF_000159835.2 | |
| *Pseudomonas amygdali* pv. *tabaci* ATCC 11528 | 1 | | | GCF_001006455.1 | |
| *Pseudomonas brassicacearum* BS3663 | 1 | | | GCF_900103245.1 | |
| *Pseudomonas brassicacearum* PA1G7 | 1 | France: Finistere | potato rhizosphere | GCF_000800585.1 | host: *Solanum tuberosum* (potato, soft-rot disease) |
| *Pseudomonas coronafaciens* pv. *porri* ICMP8961 | 1 | France | | GCF_001400915.1 | host: *Allium ampeloprasum* (leek) |
| *Pseudomonas coronafaciens* pv. *porri* LMG 28495 | 1 | Belgium: Aarsele | plant | GCF_001275725.1 | host: *Allium ampeloprasum* (leek, leaf yellowing ) |

| | synt. regions | Location Name | isolation source | Assembly Accession | additional information |
|---|---|---|---|---|---|
| *Pseudomonas coronafaciens* pv. *porri* LMG 28496 | 1 | Belgium: Menen | plant | GCF_001275735.1 | host: *Allium ampeloprasum* (leek) |
| *Pseudomonas fluorescens* ATCC 17400 | 1 | USA: California | hen's egg | GCF_000708695.2 | |
| *Pseudomonas fluorescens* AU14917 | 1 | USA | sputum | GCF_000803005.1 | host: *homo sapiens* (human, cystic fibrosis) |
| *Pseudomonas fluorescens* EK007-7t-asp | 1 | not applicable | not applicable | GCF_001931665.1 | |
| *Pseudomonas fluorescens* EK007-RG4 | 1 | | phyllosphere | GCF_001902145.1 | |
| *Pseudomonas fluorescens* ML11A | 1 | | Skin mucus | GCF_001908925.1 | host: *Salvelinus fontinalis* (fish) |
| *Pseudomonas kilonensis* BS3780 | 1 | | | GCF_900105635.1 | |
| *Pseudomonas lini* ZBG1 | 1 | France: Zellenberg | Soil | GCF_001238395.1 | |
| *Pseudomonas mandelii* 36MFCvi1.1 | 1 | | | GCF_000381285.1 | |
| *Pseudomonas marginalis* BS2952 | 1 | | | GCF_900105325.1 | |
| *Pseudomonas orientalis* BS2775 | 1 | | | GCF_900105795.1 | |
| *Pseudomonas orientalis* DSM 17489 | 1 | Lebanon | spring water | GCF_001439815.1 | |
| *Pseudomonas plecoglossicida* TND35 | 1 | India: Tamilnadu, Ottanchathiram | soil | GCF_000764405.1 | |
| *Pseudomonas putida* INSali382 | 1 | Portugal: Lisbon | Vegetable | GCF_001653615.1 | |
| *Pseudomonas putida* JQ581 | 1 | | | GCF_001630725.1 | |
| *Pseudomonas* sp. A214 | 1 | | | GCF_900156295.1 | |
| *Pseudomonas* sp. C5pp | 1 | India: Mumbai | soil | GCF_000814065.1 | |
| *Pseudomonas* sp. CFT9 | 1 | USA: Nyack River | hyporheic zone | GCF_000416255.1 | |
| *Pseudomonas* sp. FSL W5-0203 | 1 | | queso fresco | GCF_001896155.1 | |
| *Pseudomonas* sp. GM55 | 1 | | | GCF_000282395.1 | host: *Populus deltoides* (eastern cottonwood) |
| *Pseudomonas* sp. GM67 | 1 | USA: Tennessee | endopshere | GCF_000282435.1 | host: *Populus deltoides* (eastern cottonwood) |
| *Pseudomonas* sp. ICMP 19500 | 1 | New Zealand | kiwi fruit | GCF_001467145.1 | |
| *Pseudomonas* sp. QTF5 | 1 | China: Tuonamu area in Qiangtang basin | permafrost soil | GCF_000512695.2 | |
| *Pseudomonas* sp. Root569 | 1 | Germany:Cologn e | root | GCF_001427465.1 | host: *Arabidopsis thaliana* (tahle cress) |
| *Pseudomonas* sp. Root9 | 1 | Germany:Cologn e | root | GCF_001429205.1 | host: *Arabidopsis thaliana* (tahle cress) |
| *Pseudomonas* sp. WCS374 | 1 | Netherlands: Flevoland | potato rhizosphere | GCF_000698295.1 | host: *Solanum tuberosum* (potato) |
| *Pseudomonas syringae* pv. *maculicola* 90_32 | 1 | USA: California | | GCF_001293855.1 | host: *Brassica oleracea* |
| *Pseudomonas syringae* pv. *maculicola* ES4326 | 1 | | | GCF_000145845.1 | |
| *Pseudomonas syringae* pv. tomato ICMP2844 | 1 | United Kingdom: Guernsey, Channel Islands | | GCF_001401135.1 | host: *Solanum lycopersicum* (tomato) |
| *Pseudomonas syringae* pv. *maculicola* M4a | 1 | USA | | GCF_001294305.1 | host: *Raphanus sativus* (radish) |
| *Pseudomonas syringae* pv. *persicae* isolate NCPPB 2254 | 1 | | | GCF_900235805.1 | |
| *Pseudomonas syringae* pv. tomato PT23 | 1 | | | GCF_002024925.1 | host: tomato |
| *Pseudomonas umsongensis* UNC430CL58Col | 1 | | | GCF_000620285.1 | |

| | synt. regions | Location Name | isolation source | Assembly Accession | additional information |
|---|---|---|---|---|---|
| *Pseudomonas fluorescens* A3422A | 2 | USA: Alsea Valley Benton Co., OR | rhizosphere | GCF_002022335.1 | |
| *Pseudomonas fluorescens* G2Y | 2 | USA: Linn Co., OR | rhizosphere | GCF_002022365.1 | |
| *Pseudomonas fluorescens* TDH40 | 2 | USA: Philomath Benton Co., OR | rhizosphere | GCF_002022255.1 | host: Poa |
| *Pseudomonas salomonii* ICMP 14252 | 2 | | | GCF_900107155.1 | |
| *Pseudomonas salomonii* LMG 22120 | 2 | France | | GCF_001730645.1 | host: *Allium sativum* (garlic) |
| *Pseudomonas* sp. GM48 | 1 or 2 | USA: Tennessee | root | GCF_000282335.1 | host: *Populus deltoides* (eastern cottonwood) |
| *Pseudomonas syringae* pv. *maculicola* H7608 | 2 | | | GCF_001293925.1 | host: *Brassica campestris* (biennial turnip rape) |
| *Pseudomonas thivervalensis* LMG 21626 | 2 | France: Sexy-les-Bois | rhizoplane | GCF_001637285.1 | host: *Brassica napus* (rapeseed) |

### [SM16] Gene-window analysis of *Pf*AR-1, *Pf*0-1, *Pst*. DC3000, and *P. salomonii* ICMP 14252

**Gene Window Codon Analysis** From the 3347 publicly available genomes, 3 were selected, in addition to *Pf*AR-1, for assessment of local codon usage using a 10-gene sliding window approach. These 3 genomes were: *Pf*0-1, as the reference *Pseudomonas* strain closely related to *Pf*AR-1, although lacking any putative HGT region; *Pst.* DC3000, a well studied plant pathogen, which contained one putative HGT region; and *P. salomonii* ICMP14252, a garlic pathogen which contains two putative HGT regions.

The codon usages for each species were combined, analyzed using PCA, and for clarity, plotted separately. In each plot, a dense 'core' region with gene windows of similar codon usage can be seen in the top-left quadrant (Fig. S11). As expected, the putative HGT regions in *Pf*AR-1, *Pst* DC3000 and *P. salomonii* ICMP14252 are outside of this core region, although *Pf*0-1, which lacks any copy of the target HGT region, shows clear signature of other HGT-derived regions.

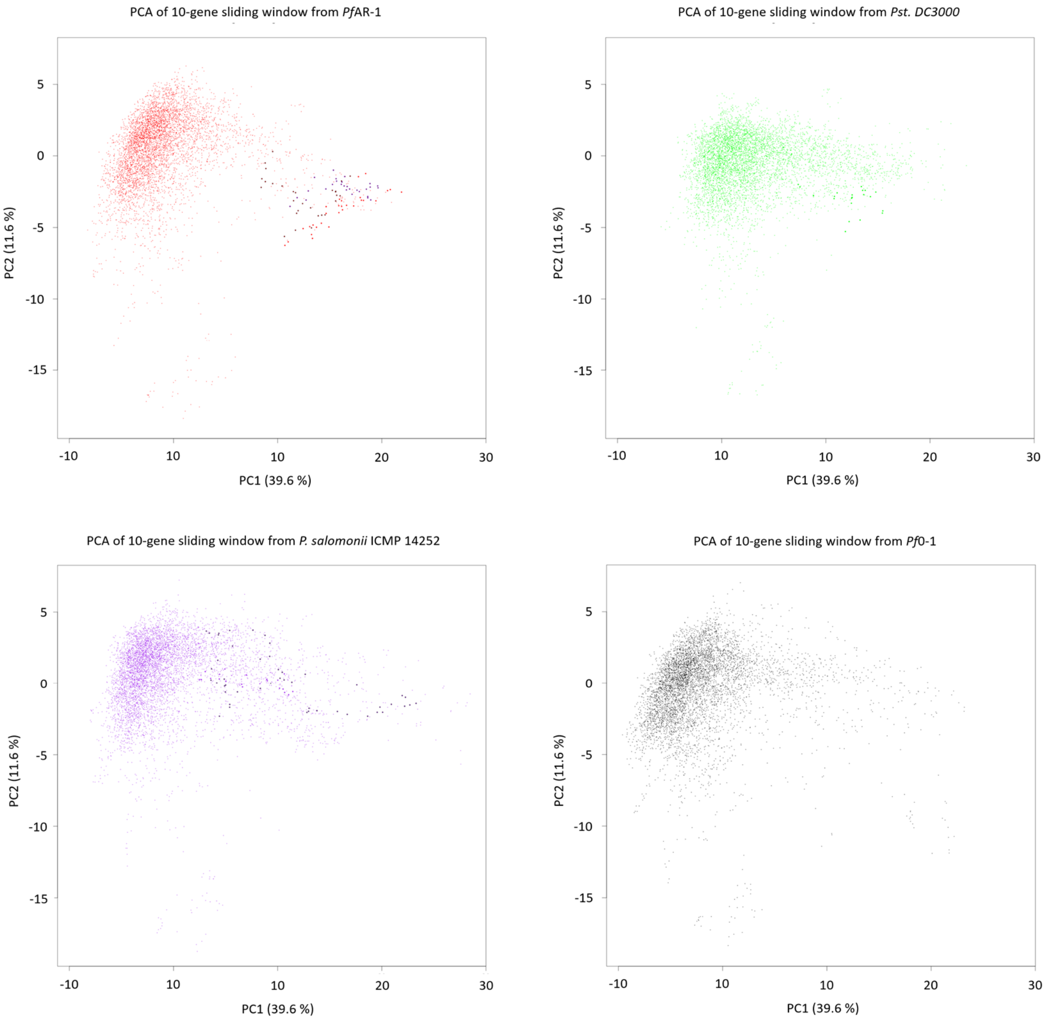

Figure S11: Principle Component Analysis (PCA) for the codon usage of *P. fluorescens* AR-1, *P. syringae* pv. *tomato* DC3000 (*Pst.* DC3000), *P. salomonii* ICMP 14252, and *P. fluorescens Pf*0-1. Bold dots show the codon usage of the putative HGT regions.

### [SM17] Genome trees for phylogenetic comparison

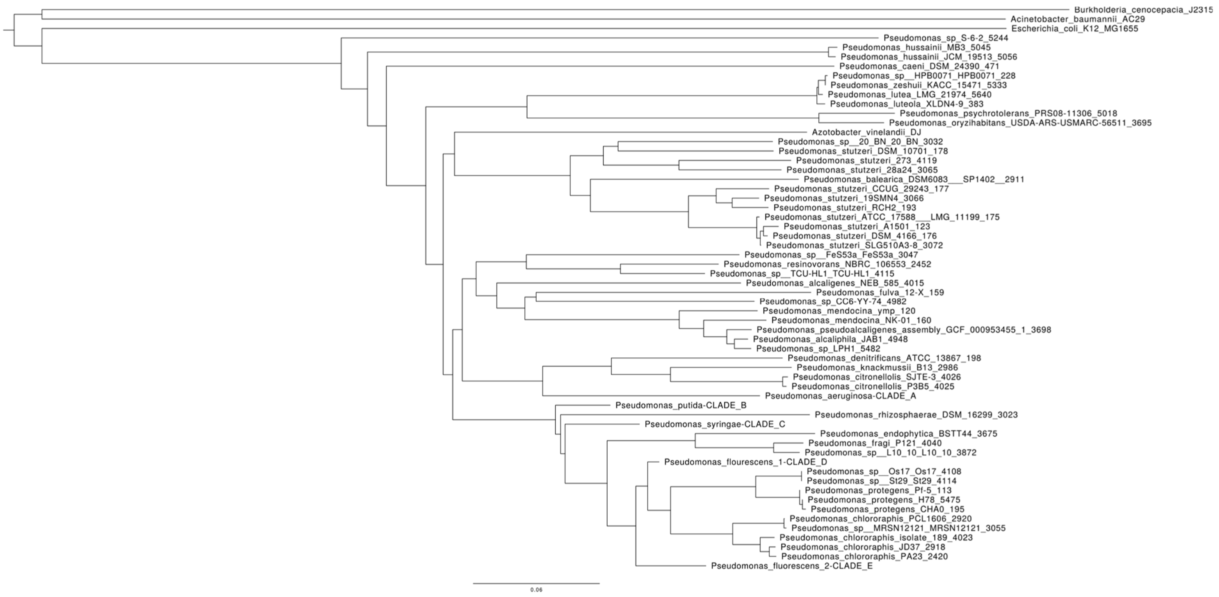

Figure S12: Top level, whole genome phylogenetic tree.

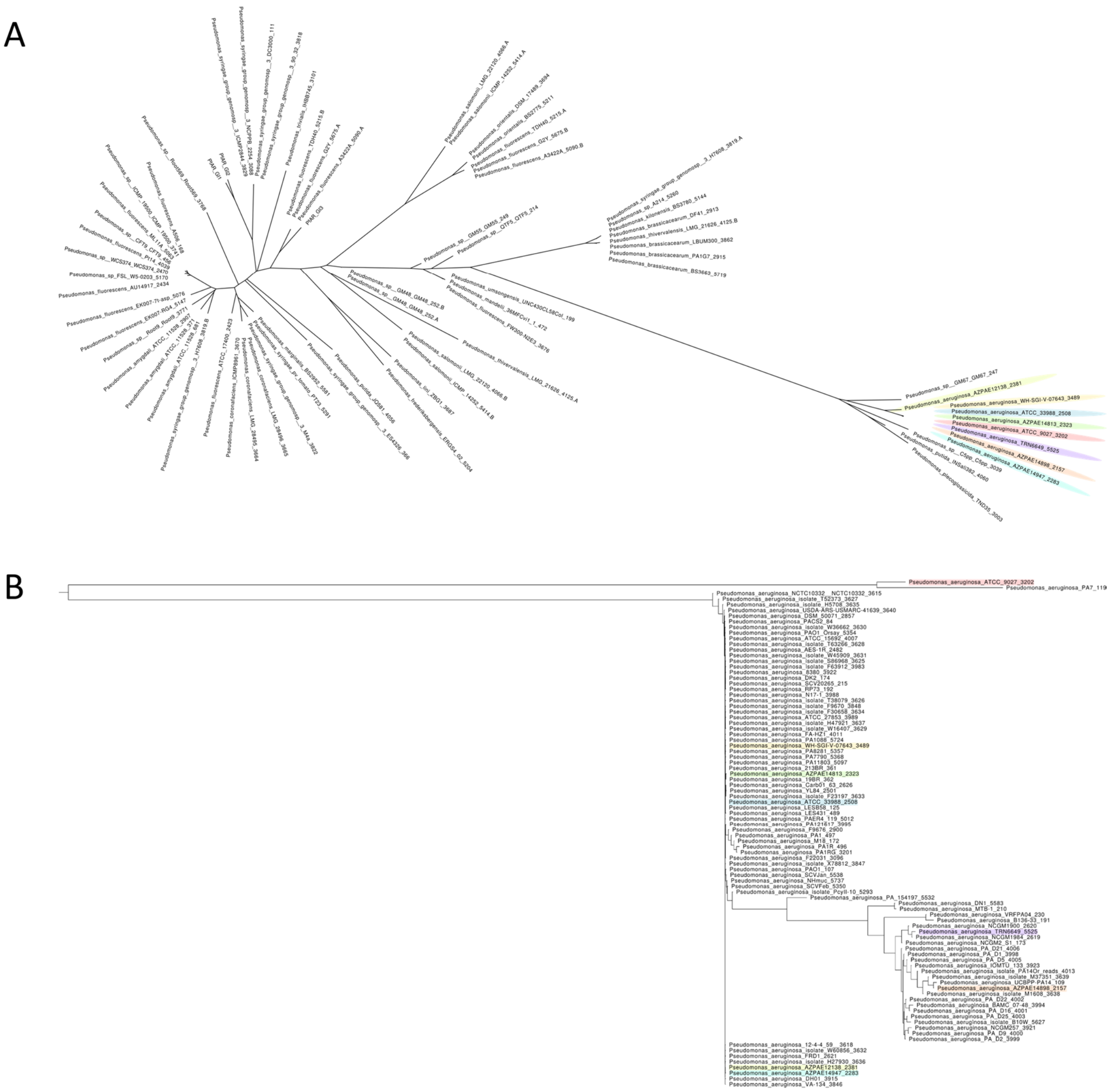

Figure S13: (A) Region specific phylogenetic tree and (B) whole genome phylogenetic tree for Clade A.

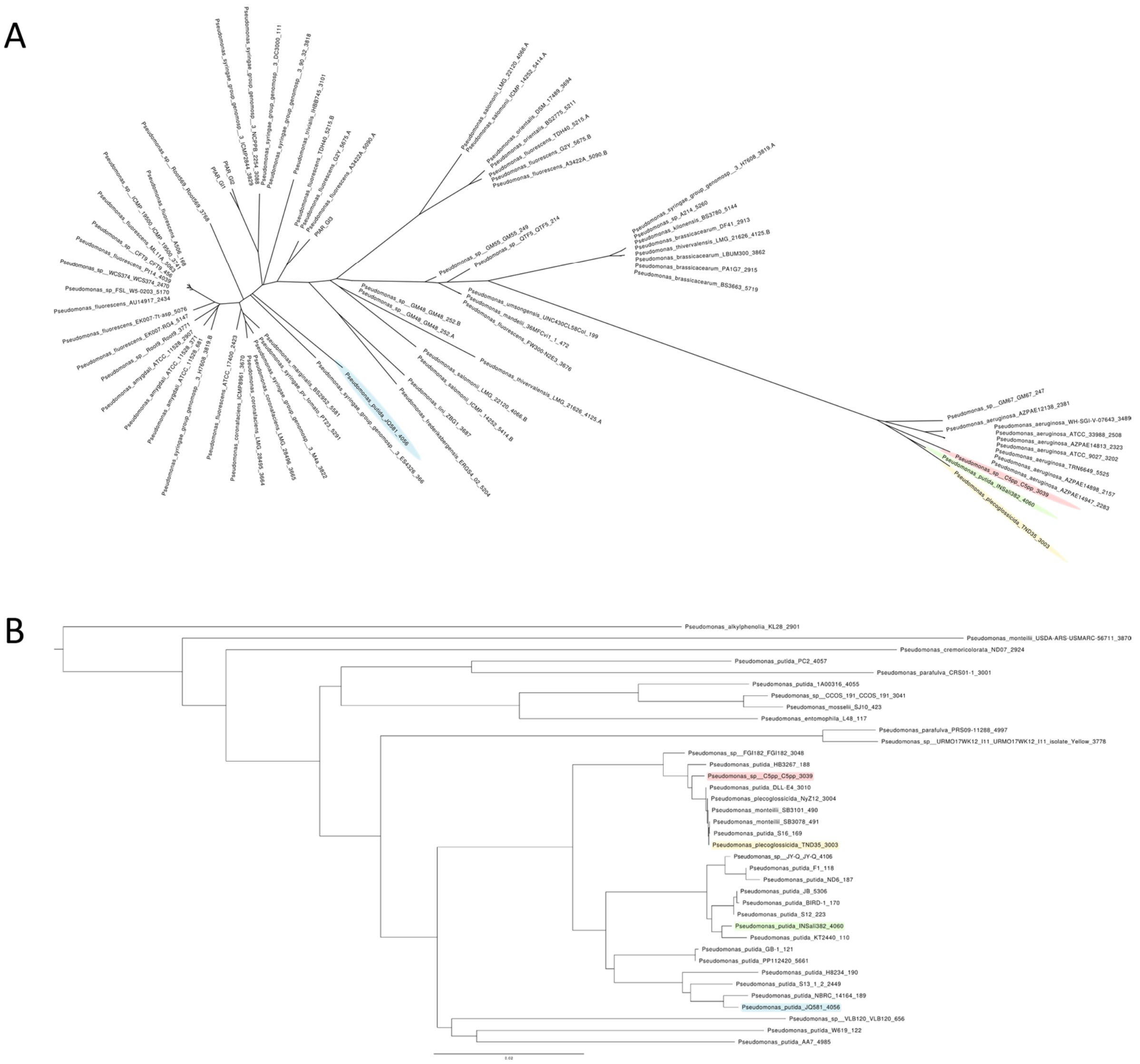

Figure S14: (A) Region specific phylogenetic tree and (B) whole genome phylogenetic tree for Clade B.

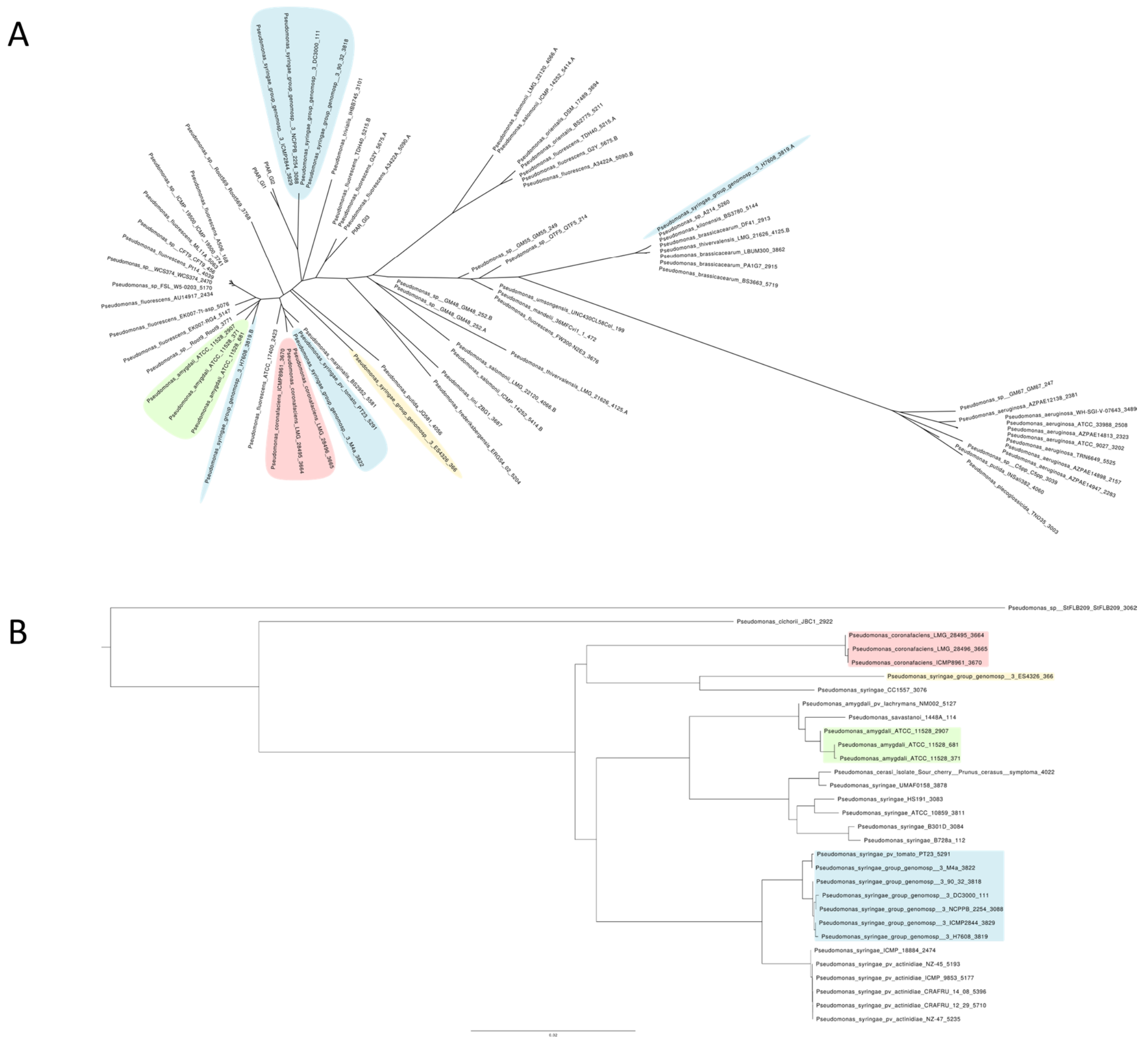

Figure S15: (A) Region specific phylogenetic tree and (B) whole genome phylogenetic tree for Clade C.

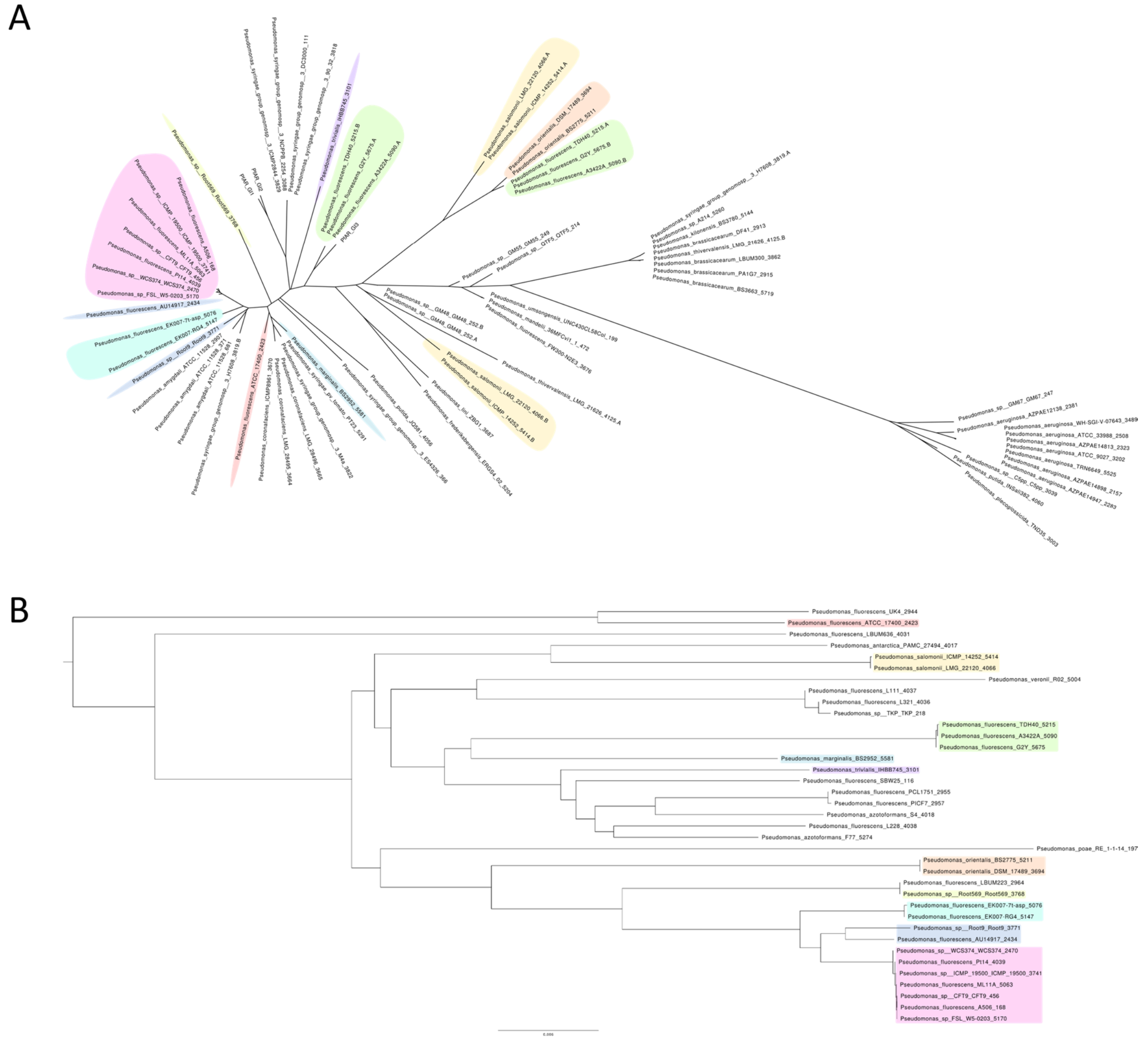

Figure S16: (A) Region specific phylogenetic tree and (B) whole genome phylogenetic tree for Clade E.

### [SM18] ANI-Values

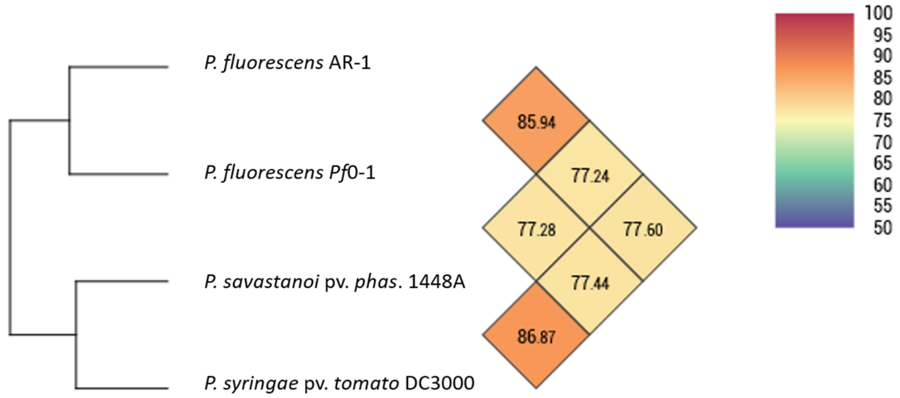

Figure S17: original ANI values calculated with OAT software (Lee et. al 2016).

### [SM19] Position of syntenic regions *Pst* DC3000, *P. salomonii*

Coordinates of syntenic regions are: *P. salomonii* ICMP 14252 (GenBank: FNOX00000000.1) region 1 on contig 102 from position 324,974 to 392,566, and region 2 on contig 114 from position 73,863 to 86,381; and for *P. syringae* pv. *tomato* DC3000 (GenBank: NC_004578.1) from 4,794,584 to 4,807,117.
